# The evolution of gene regulation in mammalian cerebellum development

**DOI:** 10.1101/2025.03.14.643248

**Authors:** Ioannis Sarropoulos, Mari Sepp, Tetsuya Yamada, Philipp S. L. Schäfer, Nils Trost, Julia Schmidt, Céline Schneider, Charis Drummer, Sophie Mißbach, Ibrahim I. Taskiran, Nikolai Hecker, Carmen Bravo González-Blas, Niklas Kempynck, Robert Frömel, Piyush Joshi, Evgeny Leushkin, Frederik Arnskötter, Kevin Leiss, Konstantin Okonechnikov, Steven Lisgo, Miklós Palkovits, Svante Pääbo, Margarida Cardoso-Moreira, Lena M. Kutscher, Rüdiger Behr, Stefan M. Pfister, Stein Aerts, Henrik Kaessmann

## Abstract

Gene regulatory changes are considered major drivers of evolutionary innovations, including the cerebellum’s expansion during human evolution, yet they remain largely unexplored. In this study, we combined single-nucleus measurements of gene expression and chromatin accessibility from six mammals (human, bonobo, macaque, marmoset, mouse, and opossum) to uncover conserved and diverged regulatory networks in cerebellum development. We identified core regulators of cell identity and developed sequence-based models that revealed conserved regulatory codes. By predicting chromatin accessibility across 240 mammalian species, we reconstructed the evolutionary histories of human *cis*-regulatory elements, identifying sets associated with positive selection and gene expression changes, including the recent gain of *THRB* expression in cerebellar progenitor cells. Collectively, our work reveals the shared and mammalian lineage-specific regulatory programs governing cerebellum development.

## INTRODUCTION

Understanding the molecular basis of phenotypic evolution, in particular in the context of the human brain, is a fundamental question in biology. The cerebellum – a brain region involved in sensory-motor control as well as cognitive, affective and social processing (*1, 2*) – has ancient origins and a conserved circuitry (*3*). However, it was shaped by a series of evolutionary innovations. The cerebellar nuclei have repeatedly expanded in number during vertebrate evolution and show additional variation in size and cell type proportions across mammals (*4, 5*). The number of cerebellar neurons increased in parallel to the expansion of the neocortex during mammalian evolution (*6*). In humans, this increase could be related to an expansion of the germinal zones during development (*7*). Additionally, we recently described an increase in the number of early fetal Purkinje cells, skewed towards the early-born subtypes, in the human lineage (*8*).

Mutations in regulatory sequences that affect gene expression have long been considered central to such evolutionary innovations (*9*), because – unlike mutations in coding sequences – they can specifically affect a subset of cell states (*10*). Recently, single-cell transcriptome studies have revealed thousands of gene expression changes between homologous neural cell types in humans and other mammals (*8, 11–14*), providing candidates for further functional investigation. Gene expression evolution is mainly driven by changes in the activity of *cis*-regulatory elements (CREs) (*15*), such as enhancers and promoters, which are bound by transcription factors (TFs) and regulate the transcription of adjacent genes in a cell type-specific manner (*16*). However, the turnover in CRE activity is typically faster than that of gene expression (*17–19*), suggesting that changes in CREs are often buffered by redundancy or compensated by the emergence, loss and repurposing of other CREs. The growing availability of high-quality mammalian genomes (*20, 21*) has opened the door to the study of mammalian CRE evolution (*22–26*). Until recently, the fast evolutionary turnover of CREs and our incomplete understanding of how their activity is encoded in their sequences severely limited the insights obtainable from genomic sequence comparisons alone. However, recent advances in machine learning have shown great promise in deciphering the regulatory code and predicting CRE accessibility from DNA sequences (*27–33*), thereby adding a powerful new tool to the investigation of CRE evolution (*14, 34–37*).

In this study, we combined previously published (*8, 38*) and newly generated single-cell measurements of gene expression and chromatin accessibility in developing cerebellar cells for six mammalian species – human, bonobo, macaque, marmoset, mouse and opossum – with machine-learning models to infer gene regulatory networks, decode the sequence grammar of CRE cell type-specificity, and reconstruct the evolutionary histories of human CREs.

## RESULTS

### Multiomic atlases of cerebellar development across mammals

To explore the gene regulatory programs underlying the development of cerebellar cell types in mammals, we extended our previous single-nucleus gene expression datasets of human, mouse and opossum (*8*) and chromatin accessibility datasets of mouse and opossum (*38*) with newly profiled developmental stages and adult white-matter enriched samples for human and mouse (Fig. 1A and table S1). Additionally, we generated a developmental atlas of chromatin accessibility in humans, and jointly profiled gene expression and chromatin accessibility from the same nuclei for three non-human primates, covering pre- and postnatal development in marmoset and early postnatal development in rhesus macaque and bonobo (Fig. 1A and table S1). To ensure optimal mapping of single-nucleus sequencing reads and minimize biases in cross-species analyses, we refined the existing genome annotations for opossum and non-human primates using publicly available (*39*) and newly generated bulk RNA-seq data (fig. S1, A and B, and table S2). Collectively, our single-nucleus datasets encompass six species and span up to eleven developmental stages, comprising 444,543 gene expression profiles and 338,985 chromatin accessibility profiles.

**Figure 1.**
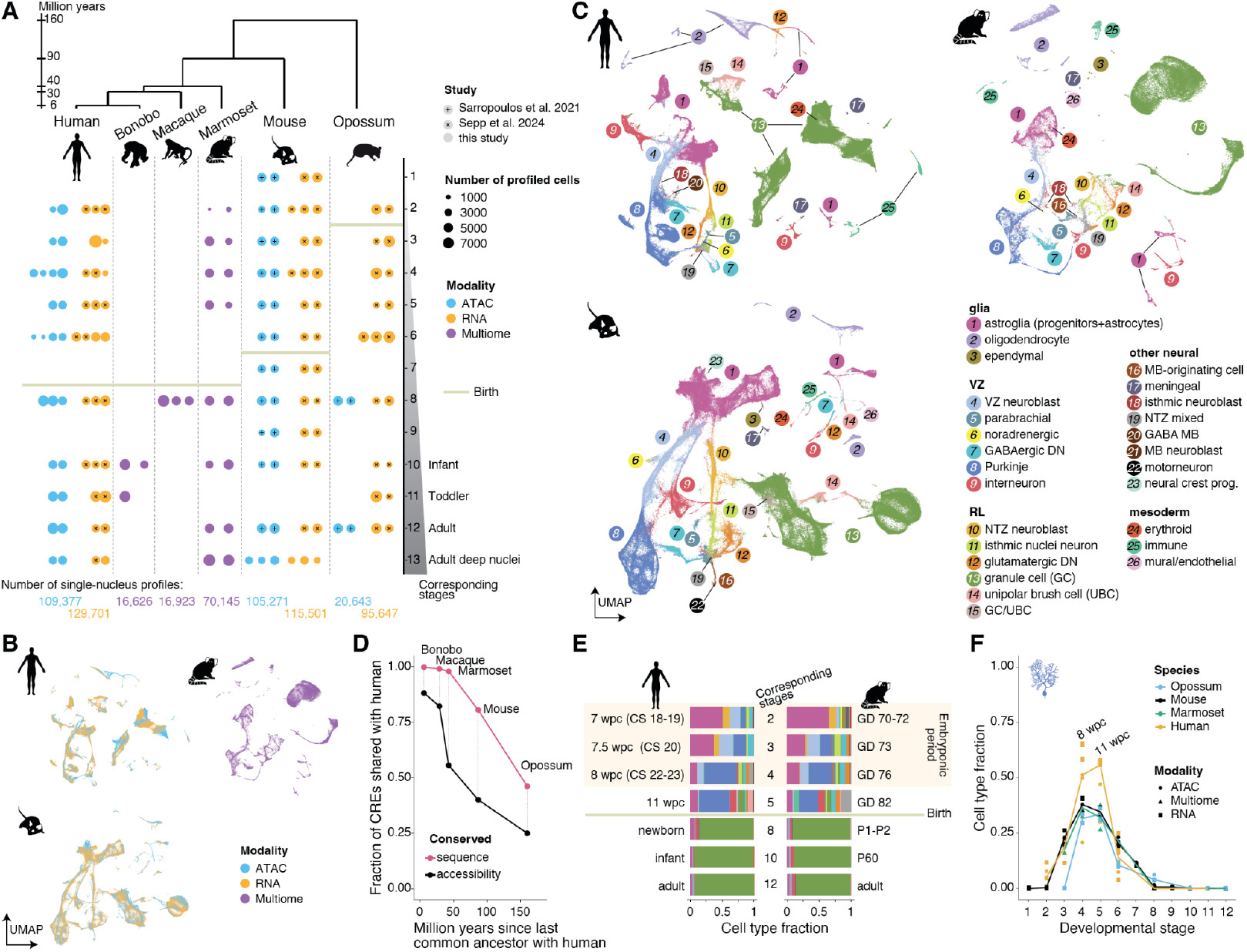
Multiomic atlases of mammalian cerebellum development. (A) Overview of the species and developmental stages included in the dataset. Dots represent individual libraries. Developmental stage correspondences as in fig. S4. (B, C) Uniform manifold approximation and projection (UMAP) of human, marmoset and mouse single-nucleus profiles integrated across developmental stages and data modalities, coloured by modality (B) or cell type (C). (D) Fraction of CREs from non-human species with an orthologous locus (sequence-conserved) or an orthologous CRE (accessibility-conserved) in the human genome. (E) Relative cell type abundances in the human and marmoset datasets at corresponding developmental stages. Colors as in (C). (F) Relative abundances of Purkinje cells within each sample (excluding non-cerebellar cells) across aligned developmental stages from four species. CS, Carnegie stage; GD, gestational day; P, postnatal day; wpc, weeks post conception.

We integrated the datasets across modalities, developmental stages, and species (Fig. 1B and fig. S2), combining label transfer approaches with manual curation to systematically annotate cells based on our previously reported (*8*) hierarchical system of cell types (level 1), differentiation states (level 2) and – for some cell types – subtypes (level 3). With the exception of rare populations of contaminating cell types from adjacent regions, samples from the newly profiled species were composed of the same cell types as detected previously in the other species (Fig. 1C, and figs. S2 and S3). For each species, we grouped cells across the finest level of our cell type annotation (level 3), developmental stage, and biological replicate, and identified peaks of open chromatin as a proxy for CREs. We identified a total of 102,197 to 554,237 reproducible CREs in each species, most of which are intronic (41-54%) or intergenic/distal (22-34%; fig. S1C). Pairwise identification of orthologous CREs between species by reciprocal syntenic alignments revealed a decrease in the fraction of human CREs shared with other species with increasing evolutionary distance as expected (Fig. 1D and fig. S1, D and E).

To establish developmental correspondences between species, we applied dynamic time warping on metrics of pseudoage inference (*40*), cellular composition, gene expression and chromatin accessibility. These analyses reaffirmed our previously inferred developmental correspondences between human, mouse, and opossum (*8*) and allowed us to infer correspondences for the non-human primate species in our dataset (Fig. 1A, fig. S4, and table S3). For the marmoset, we mapped samples from gestational days (GD) 70-76 (10-11 weeks post-conception, wpc) to human embryonic stages (7-8 wpc), while GD82 (12 wpc) shows the highest similarity to human fetal stages (11 wpc; Fig. 1E). This is in line with the delayed embryogenesis in marmoset compared to humans (*41*) even though their gestational period is only half as long (21 weeks versus 38 weeks). Newborn and early postnatal marmoset, macaque, and bonobo samples most closely resembled human samples of similar ages, in line with approximately equivalent maturation of primates around birth (*42*).

We previously identified an increase in the relative abundances of Purkinje cells in human at 8-11 wpc compared to mouse and opossum (*8*). However, we were unable to resolve when this change occurred during the approximately 90 million years of evolution separating human and mouse. Here, we used our marmoset dataset, a non-human primate for which we had access to fetal samples, to better resolve the timing of this change. We found that the relative abundances of Purkinje cells in marmoset closely resemble those in mouse and opossum (Fig. 1F), suggesting that the increase observed in humans occurred at some point during the past 40 million years.

### Gene regulatory networks of mammalian cerebellar cell types

We next used our integrated measurements of gene expression and chromatin accessibility to infer the gene regulatory networks (GRNs) underlying the development of cerebellar cell types in each species, employing SCENIC+ (*43*) to link transcription factors (TFs) to their putative CREs and target genes (Fig. 2A). We focused this analysis on human, marmoset, and mouse, for which our datasets span the entire cerebellum development. To account for the unequal representation of cell types throughout development, we grouped the cells across the three annotation levels with the goal of optimizing both the cell coverage and the resolution for distinguishing corresponding types/states across species, hereafter referred to as cell groups (Fig. 2B). We inferred 251, 220, and 198 regulons (TFs and their target CREs and genes) in human, marmoset, and mouse, respectively (tables S4 to S6). To assess the quality of the inferred GRNs, we focused on the mouse, for which more functional genomics datasets relevant to cerebellum development are available. For all examined TFs, we observed an enrichment of ChIP-seq signals in predicted target CREs (fig. S5A and table S7). CREs and the transcription start site (TSS) of their target genes are significantly more likely to fall within the same topologically associating domain (TAD), irrespective of the CRE-TSS distance (fig. S5B). Finally, our GRN performed significantly better in predicting gene expression differences between held-out mouse samples (mean R^2^ = 0.57), compared to a permuted GRN (R^2^ = 0.04) or permuted gene expression fold changes (R^2^ = 0.01; fig. S5, C and D). Collectively, these analyses support the reliability of our inferred GRNs.

**Figure 2.**
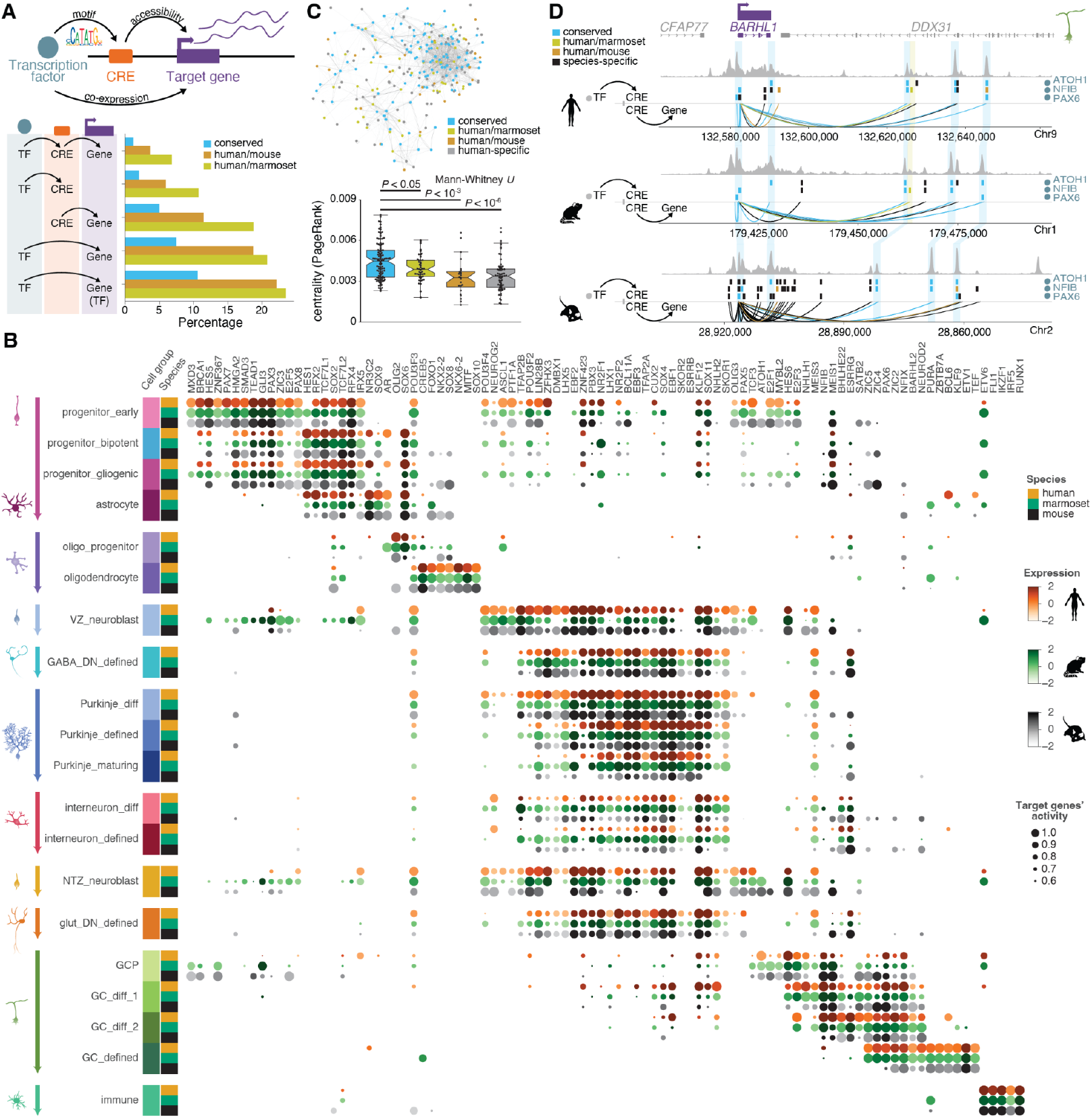
Gene regulatory networks (GRNs) of cerebellar cell types. (A) Percentage of GRN links shared between species. (B) Expression and regulon specificity of TFs with conserved activity in human, marmoset, and mouse. (C) Network of TF-TF regulatory interactions (top) and centrality per conservation group (bottom). Dots represent TFs, colors indicate conservation group, and edges depict links between TFs and TF genes. Boxes represent the interquartile range and whiskers extend to extreme values within 1.5 times the interquartile range from the box. (D) Regulatory interactions in the *BARHL1* locus in granule cell progenitors across species. Boxes represent CREs linked to the TFs shown on the right, arcs depict CRE-gene links and colors indicate conservation levels. Histograms display chromatin-accessibility profiles. CRE, cis-regulatory element; TF, transcription factor.

Focusing on orthologous TFs, we identified 114 regulons present in the GRNs of all three species. Human TFs in shared regulons exhibit higher network centrality than TFs in human-specific regulons (Fig. 2C), indicating that TFs with many regulatory interactions are more likely to remain conserved during evolution. Among the shared regulons, the conservation of connections decreases from TF-gene to CRE-gene and then to TF-CRE links (Fig. 2A), a trend also seen in the adult cerebral cortex (*43*). This finding supports the notion that individual CREs and TF binding sites are rapidly gained and lost during evolution, with the majority of these changes being compensated rather than leading to changes in gene expression (*17, 44, 45*). Indeed, despite this rewiring, the aggregated activities of regulons in each cell type are highly correlated between species (r = 0.78-0.88; fig. S5, E and F). Further supporting the high constraint in TF activity, we observed higher conservation in the regulation of TF genes (TF-TF links) compared to all genes (Fig. 2A).

We next focused on TFs with conserved cell type-specific expression and activity to prioritize core cell fate regulators in the cerebellum (Fig. 2B). This analysis revealed established regulators, including ATOH1 and PAX6 for granule cells (*46*) and SOX2 and HES1 for astroglia, alongside less characterized factors such as PURA (*47*) and SATB2 (*8*) for granule cells, as well as novel regulators like TEF in granule cells and steroid hormone receptors NR3C2 and AR in astroglia. We found that TFs with conserved cell type-specific activity control the expression of other TFs sharing the same expression specificity (fig. S6). However, in line with our previous analyses (Fig. 2A), most (91.3%) of the conserved cell-type-specific TF-gene links are mediated by CREs that are species- or lineage-specific (Fig. 2D and fig. S6). Collectively, our analyses revealed that despite the fast turnover in individual CRE sequences, the core transcription factor networks driving cell type-specific expression programs in the developing cerebellum have remained conserved throughout mammalian evolution.

### Sequence-based models of CRE spatio-temporal specificity

Building on our findings regarding the conservation of core TFs and the well-known preservation of short sequence motifs recognised by TFs throughout animal evolution (*48, 49*), we next asked whether there are specific sequence features shared between CREs accessible in the same cell type regardless of their species of origin or evolutionary relationships. To this end, we first summarized the accessibility of human and mouse CREs across corresponding cell groups (Fig. 2B) and developmental stages. We used non-negative matrix factorisation (NMF) to project human and mouse CREs into a set of 18 factors that capture cell state- and time-specific accessibility patterns (Methods; Fig. 3A, fig. S7, A to C, and table S8). Human and mouse CREs are evenly distributed across these factors (fig. S7D), suggesting that the major axes of biological variation in CRE accessibility are shared across mammals.

**Figure 3.**
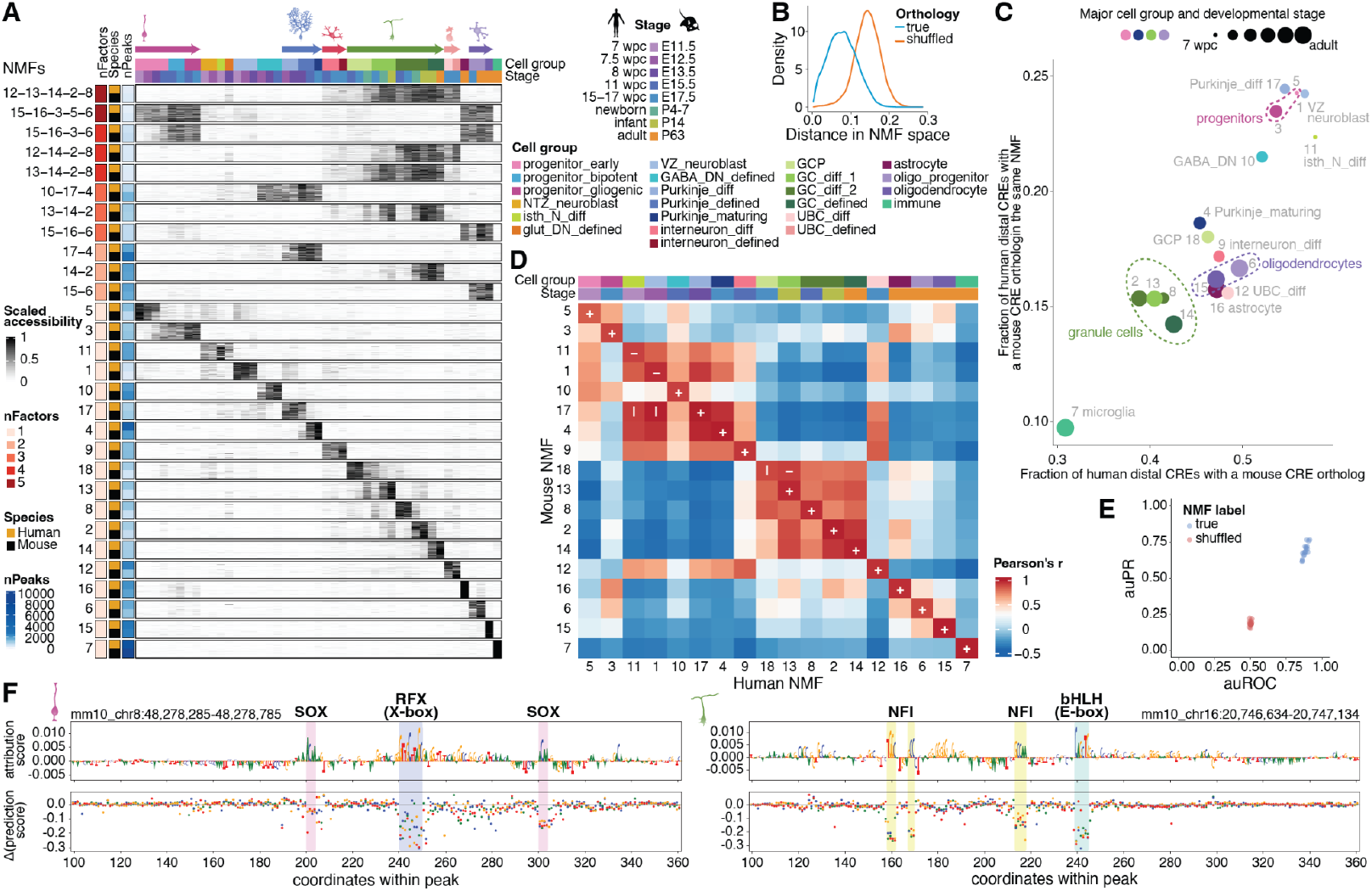
Sequence-based models of CRE accessibility. (A) CRE accessibility (quantile-normalized CPM, max-scaled) across human and mouse cell groups and developmental stages for CREs assigned to one (bottom) or multiple (top; only most frequent combinations shown) NMFs. 100 CREs from each group and species were randomly selected for visualization. (B) Euclidean distance in NMF loadings between true and shuffled human and mouse CRE orthologs. (C) Fraction of human distal CREs with conserved accessibility (x-axis) and specificity (y-axis) in mouse across NMFs. (D) Pearson’s *r* in TF motif enrichment scores for mouse and human CREs assigned to each NMF. (E) Accuracy of sequence-based model in NMF assignment prediction of held-out mouse CREs (blue) versus the same CREs with shuffled labels (red). (F) DeepExplainer and *in silico* mutagenesis profiles of mouse CRE sequences assigned to NMF 5 (early progenitors) and 18 (granule cell progenitors). Motifs of relevant TFs are highlighted.

One third of CREs were assigned to one NMF (fig. S7E), reflecting their cell state- and time-specificity, in agreement with our previous observations for mouse CREs (*38*). In addition, we detected substantial numbers of CREs that contribute to two or more NMFs, which frequently belong to the same or developmentally related cell types, or to the same developmental period (fig. S7F). Orthologous CREs between human and mouse show significantly higher similarity in the NMF space compared to shuffled orthology pairs (Fig. 3B), recapitulating the notion that a sizable subset of CREs has retained the same spatiotemporal activity across cerebellar cell types during mammalian evolution (*38*). However, overall CRE turnover is fast, with 31-58% of orthologous distal CREs accessible in both species, and only 9-24% of these assigned to the same NMF (Fig. 3C). NMFs associated with earlier stages of cerebellar development show higher conservation in both metrics, whereas divergence is highest for the NMF associated with immune cells (Fig. 3C), consistent with our prior inferences from mouse and opossum (*38*). Although most CREs are specific to one species, human and mouse CREs assigned to the same NMF show similar patterns of TF motif enrichment (Fig. 3D), indicating that they share sequence features.

To gain deeper insights into the sequence grammar of cerebellar CREs, we made use of recent advances in applying deep learning models for predicting CRE accessibility from DNA sequence (*27–29, 31–33*). Starting with mouse, we trained a multi-class multi-label classifier based on a hybrid architecture that combines convolutional and recurrent neural networks, akin to previous work (*29, 32*). The resulting sequence-based model was able to predict the assignment of mouse CREs to NMFs (i.e., their spatiotemporal specificity) with high accuracy (auROC:0.88, auPR: 0.69; Fig. 3E and table S9): it effectively assigned low prediction scores to sequences lacking known regulatory activity (median: 0.04) and high prediction scores to CREs accessible in multiple NMFs (median: 0.95; fig. S8A). CREs specific to a subset of NMFs were assigned lower prediction scores (median: 0.40) but significantly higher than CREs not associated with that NMF, which were similar to those observed for random intergenic regions (median: 0.05; fig. S8A). Feature attribution analyses (Methods) revealed that these predictions were driven by biologically relevant features, as the model identified relevant TF motifs in the mouse CRE sequences (Fig. 3F and fig. S8B). For example, the model learned to recognise SOX and RFX motifs in CREs specifically accessible in early progenitors, such as mm10_chr8:48,278,285-48,278,785, while predictions in CREs specific to granule cell progenitors, such as mm10_chr16: 20,746,634-20,747,134, rely on bHLH TFs (e.g., ATOH1) and NFI motifs (Fig. 3F). Thus, deep learning models are able to learn the regulatory codes of cerebellar cell types and infer the accessibility of unseen sequences.

### The sequence grammar of cerebellar CREs is conserved across mammals

Having projected human and mouse sequences in the same NMF feature space, we next assessed the conservation of the sequence grammar of cerebellar CREs by applying the mouse-trained model to human CREs. Strikingly, the model achieved almost as high accuracy across species (median auROC: 0.864) as within species (median auROC: 0.883), and the same was true for a model trained on human CREs (within-species median auROC: 0.8764; across-species median auROC: 0.8758; Fig. 4A and fig. S9A). Additionally, the human- and mouse-trained models showed high correlation in their predictions of marmoset CRE accessibility (median Pearson’s r = 0.83 for the same NMF versus r = 0.48 for different NMFs), suggesting they have learned similar sequence features (Fig. 4B and fig. S9B). To evaluate whether training models on CREs from multiple species would lead to an improved performance, as suggested by previous studies (*50*), we combined human and mouse CREs within a single model. We first kept the size of the training set constant by using 50% of CREs from each species, observing a slight increase in prediction scores (Fig. 4C and fig. S9C). We then combined all human and mouse CREs, effectively doubling the model’s training set, and reached a median auROC of 0.88 across both species, comparable to that of single-species models in their species of training. To evaluate the performance of this model, which we termed DeepCeREvo (Deep-learning of Cerebellar Regulatory Evolution), across even larger evolutionary distances, we used our snATAC-seq data for the marsupial opossum (*38*), which separated from eutherian (“placental”) mammals (such as human and mouse) about 160 million years ago (MYA). Differentially accessible regions (DARs) across opossum cerebellar cell groups exhibited significantly higher predictions in matching NMFs (Fig. 4D and fig. S9D) Thus, our analyses show that the CRE sequence codes that define cerebellar cell types have remained conserved for at least 160 million years of mammalian evolution.

**Figure 4.**
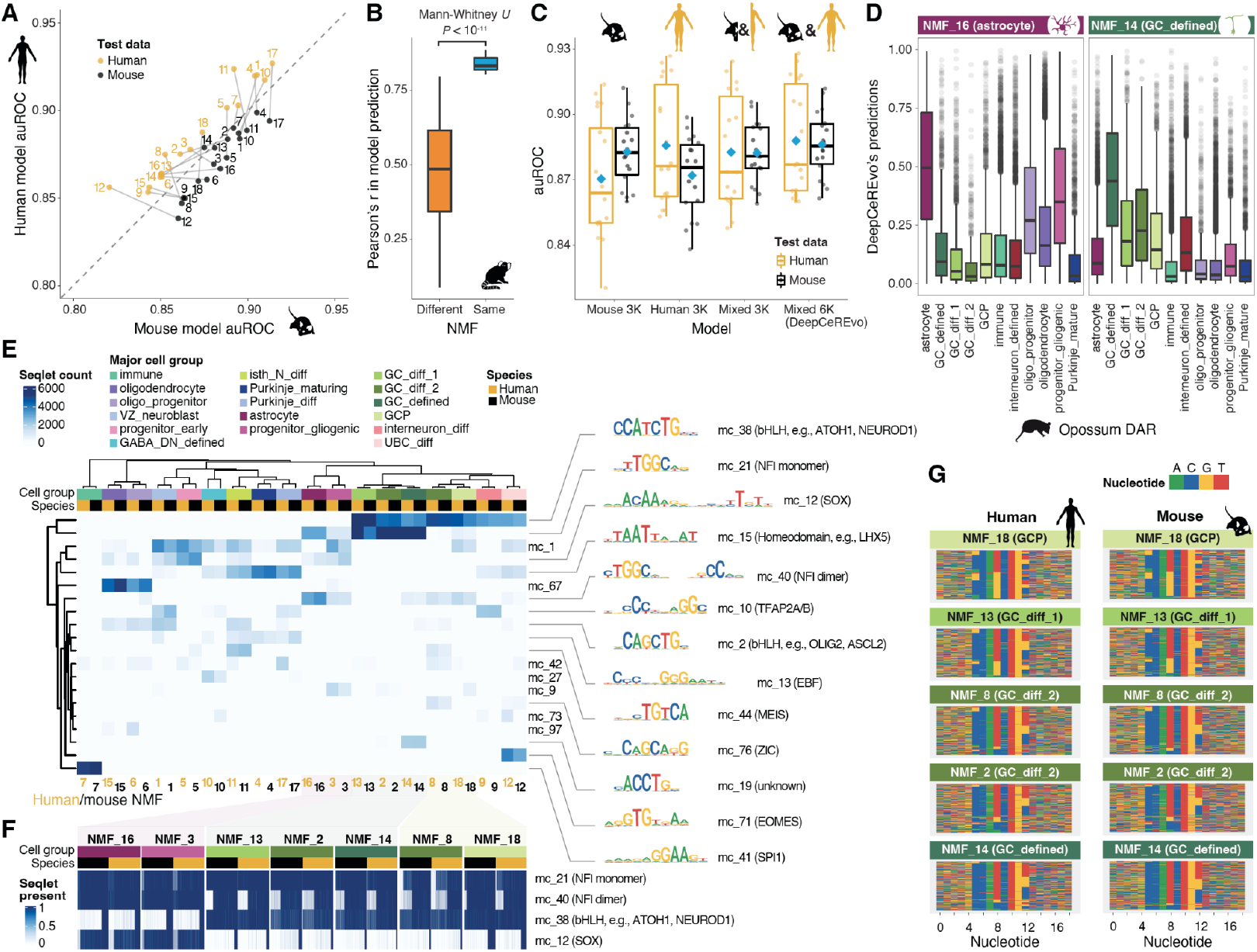
Conserved CRE sequence grammar across mammals. (A) Accuracy (auROC) of sequence-based models trained on human or mouse sequences in predicting the accessibility of unseen sequences from both species. (B) Pearson’s correlation coefficients (r) of human and mouse model predictions for marmoset sequences across the same or different NMF pairs. (C) DeepCeREvo achieves improved performance by training on both human and mouse sequences. Boxplots show auROC metrics per NMF class in human and mouse unseen sequences for models trained on CREs from human, mouse or both species. (D) Representative examples of DeepCeREvo’s predictions for opossum DARs. Full figure shown in fig. S9D. (E) Hierarchical clustering of human and mouse NMFs based on the number of motif instances with high attribution scores (seqlets) for each NMF and motif cluster (mc). Motif clusters with at least 1000 seqlets are shown, along with their respective consensus logos. (F) Presence of one or more NFI (mc_21, mc_40), E-box (mc_38) or SOX (mc_12) seqlets in human and mouse CREs with high predictions for NMFs associated with astroglia and granule cells. (G) Nucleotide composition of human and mouse E-box seqlets (mc_38) in CREs with high predictions for NMFs associated with granule cell differentiation. Cell group annotations in D-F refer to the group with the highest loading for each NMF. In B-D, boxes represent the interquartile range and whiskers extend to extreme values within 1.5 times the interquartile range from the box.

We next sought to identify the sequence features driving DeepCeREvo’s predictions. We used TF-MoDISco (*51*) to identify short sequence instances that are important for the model’s predictions (seqlets), separately for each human and mouse NMF. Clustering these groups based on the identified patterns, we observed the highest similarity between human and mouse CREs assigned to the same NMF, further demonstrating the high conservation of CRE sequence grammar (Fig. 4E and fig. S10, A and B). The identified patterns include the motifs of major TF regulators of cerebellar cell types, such as motifs of SOX TFs for progenitors (NMF 3 and 5), EOMES for UBCs (NMF 12), MEIS for neurons located at the nuclear transitory zone (NMF 11), and SPI1 for microglia (NMF 7). We also observed differences in motif importances between factors representing distinct differentiation states of the same cell type. For example, while E-box and NFI motifs are important throughout granule cell differentiation, GC-rich SP1 motifs primarily contribute to accessibility in granule cell progenitors, whereas the importance of the ETV1 motif increases in mature granule cells (fig. S10C). Similarly, while homeodomain motifs are important throughout Purkinje cell differentiation, TFAP2A motifs contribute to accessibility in differentiating but not mature neurons (fig. S10D).

Finally, we identified patterns that are important across multiple factors, most notably E-box motifs (NMF 2, 8, 9, 13, 14, 18) and NFI motifs (NMF 2, 3, 8, 13, 14, 16, 18), in line with previous observations (*38, 52*). Cell type-specificity in these cases arises from the combinatorial presence of TF motifs in the same CRE sequence. For example, when considering the 2,500 CREs with highest prediction in NMF 3 (bipotent and gliogenic progenitors), NFI motifs co-occur with SOX motifs in 87% of human and 94% of mouse CREs, a combination rarely found in CREs accessible during granule cell differentiation (Fig. 4F and fig. S10E). By contrast, 76-99% of CREs accessible in granule cells combine NFI motifs with an E-box motif (Fig. 4F and fig. S10E). CRE specificity is further achieved through subtle differences in motif sequences. While E-box motifs are used throughout granule cell differentiation (motif cluster 38), we observed a shift from a mixture of CAGCTG and CATCTG motif instances in granule cell progenitors (NMF 18) towards a strict requirement for CATCTG in CREs specific at later stages of differentiation (Fig. 4G). Based on our GRN analysis (Fig. 2B), we hypothesize that this might reflect the shift from ATOH1 to NEUROD2, with both TFs recognising E-box motifs. Although subtle, these differences are conserved between human and mouse and detected by DeepCeREvo, suggesting that they play important roles in the regulation of TF binding to CREs.

### Inferring the evolutionary histories of human CREs accessible in cerebellar cell types

Determining the evolutionary histories of human developmental CREs is challenging due to their high evolutionary turnover (*17*) and to the limited access to non-human primate developmental samples. We reasoned that the high accuracy of DeepCeREvo’s predictions and the overall conservation of the CRE sequence grammar for cerebellar cell types would allow us to predict the accessibility profiles of human CREs in other mammals and thus infer their evolutionary histories. We identified orthologous sequences of human CREs in 240 eutherian mammals (*21*) and used DeepCeREvo to predict their accessibility profiles across cerebellar cell types for each species (Fig. 5A and table S10). To differentiate CREs potentially linked to evolutionary innovations from those subject to high turnover through neutral processes, we decided to focus on CREs that, after emergence, were preserved in a majority of species in a clade. For this, we compared prediction scores of orthologous sequences for each NMF within and outside the clade and applied a phylogenetic approach (*53*) to determine statistical significance (fig. S11 and table S11). Out of 554,237 human CREs, we could assign 64,279 (11.6%) to distinct evolutionary clades in at least one NMF (Fig. 5B, fig. S12, and table S12). Eutherian-shared CREs (n = 34,097) include elements that emerged in the common ancestor of eutherians ~100 MYA or earlier during evolution and have since retained their accessibility in at least one NMF. Among younger CREs, 3,427 are shared within primates, 8,591 within simians, 7,562 within catarrhines, and 8,877 within great apes. Finally, we pinpointed 3,018 human CREs that likely emerged within the last 6.5 million years and are specific to the human lineage (Fig. 5B and fig. S12). Of note, assignment rates in our approach are higher for sequences with a minimum age of at least ~100 million years or less than ~29 million years: the sequences in the former group include those predating eutherians and are likely under strong evolutionary constraints, while the latter group is characterized by more relaxed preservation criteria, given the limited number of species in these clades (Fig. 5C).

**Figure 5.**
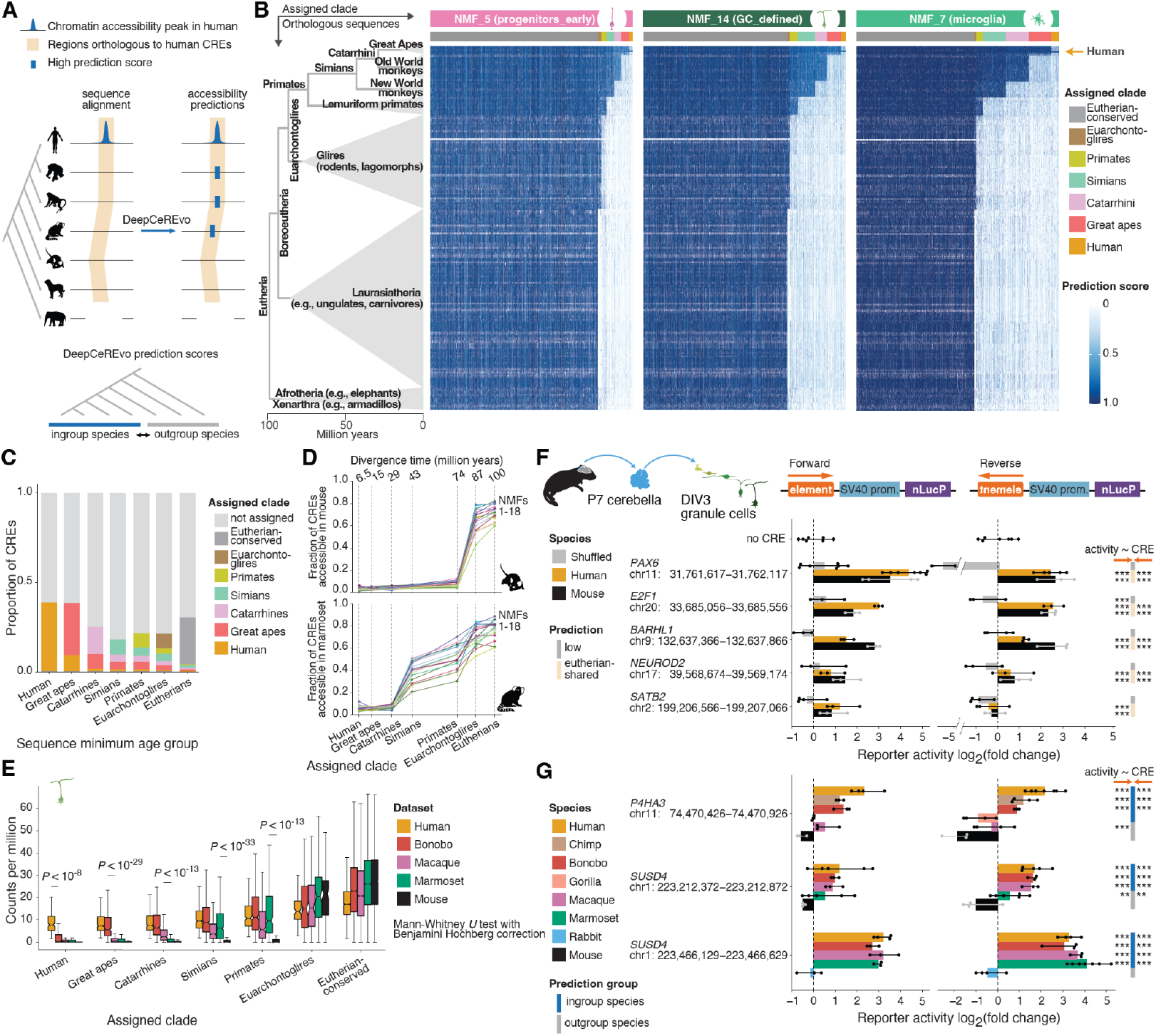
Evolutionary classification of human CREs. (A) Scheme on assignment of human CREs to different evolutionary clades using DeepCeREvo. (B) Human CREs inferred as preserved in different clades based on DeepCeREvo prediction scores on orthologous regions across 227 mammalian species for selected NMFs. (C) Proportions of CREs assigned to different clades, grouped by minimum age of the sequence. (D) Fraction of human CREs in each NMF across different evolutionary clades, with orthologous regions accessible in mouse (top) or marmoset (bottom). (E) Normalized fragment counts in mature granule cells (GC_defined) for regions that are orthologous to human CREs in NMF_14. Regions are grouped according to their predicted evolutionary histories. Boxes represent the interquartile range and whiskers extend to extreme values within 1.5 times the interquartile range from the box. (F, G) Luciferase reporter assays in mouse primary granule cells testing the enhancer activity of CREs predicted to be eutherian-shared (F) or having emerged in the last 43 million years (G). CREs were placed in front of the SV40 promoter in forward (left) or reverse (right) orientation. Bars and error bars display the mean normalized and scaled reporter activity and its range; points denote biological replicates. *P*-values relative to the constructs without an enhancer were estimated using linear mixed models, corrected for multiple testing using the Benjamini-Hochberg method, and are shown for each orientation only for bars with log_2_(fold change) ≥ 0.5. ***, *P* < 0.001; **, *P* < 0.01.

To validate the inferred evolutionary histories, we examined the accessibility profiles of human CREs in their orthologous loci in the mouse, marmoset, macaque, and bonobo datasets. We observed a good concordance between the experimentally measured accessibilities and the inferred evolutionary history of each CRE group (Fig. 5, D and E). For example, amongst human CREs predicted to have emerged between 43 and 87 MYA in the common ancestor of simians or primates, an average of 39-52% are accessible in the corresponding cell types in marmoset but only 4.8-7.5% in mouse (Fig. 5D). Furthermore, among the human CREs specific to mature granule cells (NMF 14) – a cell group with good coverage across all species – those predicted to be eutherian- or euarchontoglires-conserved show similar levels of accessibility across species. On the other hand, CREs predicted to have emerged between 29 and 43 MYA in the common ancestor of catarrhines are significantly more accessible in human, bonobo, and macaque compared to those in marmoset and mouse (Fig. 5E).

To assess whether our predictions about the evolution of CRE accessibility extend to enhancer activity, we performed luciferase reporter assays in primary cultures of mouse granule cells. Single-nucleus multiome profiling of cells isolated from P7 mouse cerebella and cultured *ex vivo* for 3 or 6 days confirmed that in these cultures (i) most (94-95%) cells are granule cells at various stages of differentiation, and (ii) the majority (71-88%) of highly variable CREs observed *in vivo* in the granule cell lineage remain accessible (CPM > 5; fig. S13, A to F). We first tested, in both orientations, human and mouse sequences for five eutherian-shared CREs (table S13) accessible in granule cell progenitors or differentiating granule cells using 3-day cultures that effectively capture these differentiation states (fig. S13, C to E). For four of these CREs, human and mouse sequences exhibited significant enhancer activity (log_2_(fold change) ≥ 0.5, *P* < 0.05, linear mixed model) in both orientations, while none of the shuffled sequence controls showed orientation-independent activity above the signals from constructs without an enhancer (Fig. 5F and table S13). We then considered 12 human CREs predicted to have emerged in the last 43 million years. Of these, five showed enhancer activity in reporter assays in at least one orientation (fig. S13G). The lower enhancer activity of the recently-emerged CREs compared to eutherian-shared CREs is consistent with similar trends observed for accessibility (Fig. 5E). For the three human CREs with orientation-independent enhancer activity, we additionally tested orthologous sequences from 4-5 species in the constant *trans* environment of mouse granule cells (table S13). A CRE in a *P4HA3* intron, predicted to have great-ape-specific accessibility, shows orientation-independent enhancer activity for sequences from 3 out of 4 great ape (ingroup) species, and not for sequences from the 2 outgroup species tested (Fig. 5G), consistent with model predictions and feature attribution scores (fig. S14A). Additionally, for two CREs near the *SUSD4* gene, the ability to robustly enhance reporter expression is confined to the sequences from the Catarrhini and simian species, respectively, in line with their predicted accessibilities (Fig. 5G and fig. S14, B and C). Thus, applying DeepCeREvo across mammalian genomes facilitates the study of enhancer evolution.

### Sequence features of human CREs across ages

We next asked how incorporating DeepCeREvo’s predictions improved our inference of CRE evolutionary histories compared to traditional sequence-based phylogenetic methods. Eutherian-shared distal CREs show significantly higher sequence conservation in comparison to other categories (Fig. 6A), in agreement with the long established association between non-coding sequence constraint and conservation of regulatory activity (*21, 54, 55*). However, DeepCeREvo’s prediction scores averaged across eutherian mammals and primates substantially outperformed sequence conservation in predicting the conservation of human CRE accessibilities in mouse and marmoset respectively (Fig. 6B and fig. S15), demonstrating the power of deep learning models to complement multiple sequence alignments for the identification of conserved regulatory regions.

**Figure 6.**
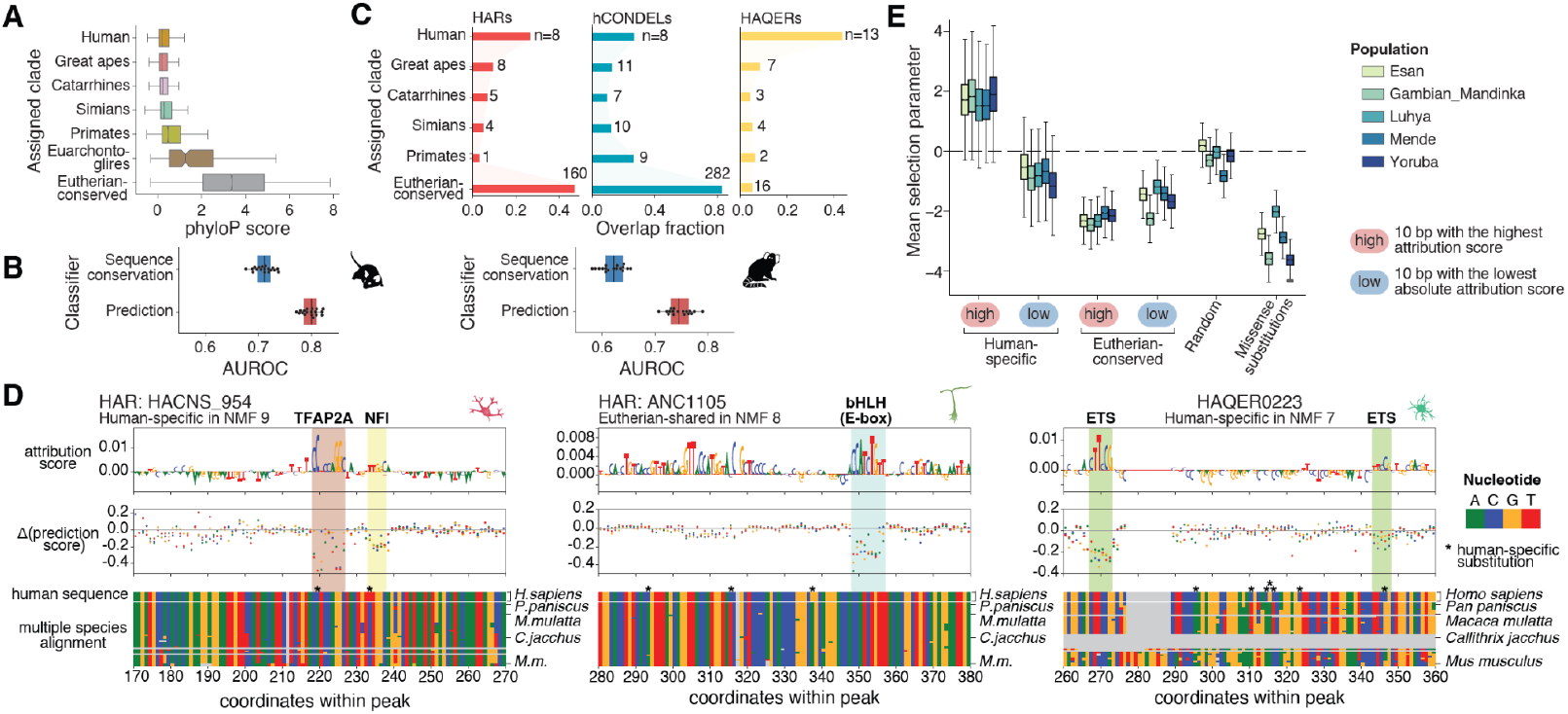
Sequence features of human CREs across age. (A) Sequence conservation of human CREs assigned to different clades. (B) Area under the receiver operating curve (AUROC) for classifiers based on DeepCeREvo prediction or sequence conservation scores averaged across eutherians/primates for assigning highly variable CREs per NMF in mouse (left) and marmoset (right). Each point corresponds to an NMF. (C) Fractions of human CREs from different evolutionary clades that overlap with regions with accelerated substitutions (HARs), deletions (hCONDELs) specific to human, or human ancestor quickly evolved regions (HAQERs). (D) DeepExplainer and *in silico* mutagenesis profiles, along with multiple sequence alignment across 43 primates and mouse, for HACNS_954 in interneurons, ANC1105 in differentiating granule cells, and HAQER0223 in microglia. (E) Posterior distribution of mean selection parameters for sequences with high and low attribution scores in human-specific and eutherian-shared CREs, inferred from segregating sites in individuals from five African populations. In A,B,E, boxes represent the interquartile range and whiskers extend to extreme values within 1.5 times the interquartile range from the box.

Previous studies have identified non-coding regions associated with evolutionary innovation in humans based on elevated sequence substitutions or deletions (*22–25, 56, 57*). Regions showing accelerated substitutions (HARs) or deletions (hCONDELs) in the human lineage, but that are conserved across other vertebrates, show only modest overlaps with our inferred human-specific CREs (Fig. 6C). Conversely, DeepCeREvo classified many HARs and hCONDELs as conserved CREs across eutherian mammals, suggesting that many of the human-specific sequence alterations in these otherwise conserved regions do not affect accessibility within cerebellar cell types. Consistently, by examining attribution profiles of individual sequences, we observed that accelerated regions with predicted human-specific accessibility experienced substitutions in positions important for the model’s predictions, such as the emergence of TFAP2A and NFI motifs in HACNS_954 accessible in GABAergic interneurons (Fig. 6D). Conversely, changes in regions with predicted conserved activity, such as ANC1105 in differentiating granule cells, fall outside of the relevant TF motifs and are thus unlikely to affect CRE accessibility in the examined cell type (Fig. 6D). However, given that conserved CREs are often used in many cell types (*38*), we cannot rule out human-specific gains or losses of accessibility in cell types of other brain regions or organs.

In contrast to HARs and hCONDELs, human ancestor quickly evolved regions (HAQERs), which represent the fastest evolving regions in the human genome without a requirement for conservation (*24*), are significantly enriched amongst the CREs we identified as human-specific (Fisher’s exact *P* < 10^−6^; Fig. 6C). For instance, one of the human-specific substitutions in HAQER0223 gave rise to an ETS motif, (e.g., SPI1 motif), explaining the predicted human-specific accessibility in microglia (Fig. 6D).

To investigate whether human-specific sequence changes underlying the predicted accessibility innovations were driven by positive selection in the human lineage, we inferred the type and extent of selective pressures shaping cerebellar CRE sequences in human populations, following a previous approach (*24, 58*). As CRE activity can be encoded in only a subset of its sequence, we considered the 10 bp sequences with the highest and lowest attribution scores for each CRE. We computed derived allele frequency spectra (dAFS) for the inferred human-specific and eutherian-conserved CREs using segregating polymorphisms in five African populations, and evaluated mean selection parameters acting on each population. For the eutherian-conserved CREs, nucleotides with both high and low attribution scores have been under negative selection (Fig. 6E), suggesting that the majority of conserved CRE sequences are relevant for their activity and have thus remained under constraint. By contrast, nucleotides with low attribution scores in the inferred human-specific CREs evolve neutrally whereas those with high attribution scores show signs of positive selection in (Fig. 6E). These findings suggest that sequence substitutions contributing to the predicted human-specific accessibilities have driven evolutionary innovations in humans.

### Human CREs associated with evolutionary innovation in gene expression

We next sought to assess the impact of CRE evolution on the gene expression programs of cerebellar cell types. We adapted our previous approach (*8*) to identify genes with major changes in expression between human and mouse within cell groups assigned to each NMF, considering both absolute and relative expression levels to limit the effect of technical confounders (fig. S16). Across all NMFs, we identified 1079 genes that are highly expressed exclusively in a human NMF and 812 of such genes in the mouse (fig. S17, A and B, and table S14). We additionally used the marmoset and opossum datasets to track, when possible, the direction and evolutionary timing of each gene expression change (Fig. 7A). For cell types shared with other brain regions (such as microglia, astrocytes, and oligodendrocytes), we observed concordant differences in expression levels in independent datasets (*11, 59*) (fig. S17C). Genes with divergent expression between humans and mice are significantly enriched for encoding membrane-localized proteins, particularly those involved in signaling and synaptic function (fig. S17D and table S15), suggesting that some cerebellar cell types may have altered their connectivity patterns during mammalian evolution. This trend is especially pronounced in the differentiating granule cell lineage, where several cadherins (CDH8, CDH20, CDH10, CDH11) exhibit higher expression in mice, while contactins (CNTN5, CNTN4, CNTN1) and glypicans (GPC5, GPC6) are more highly expressed in humans.

**Figure 7.**
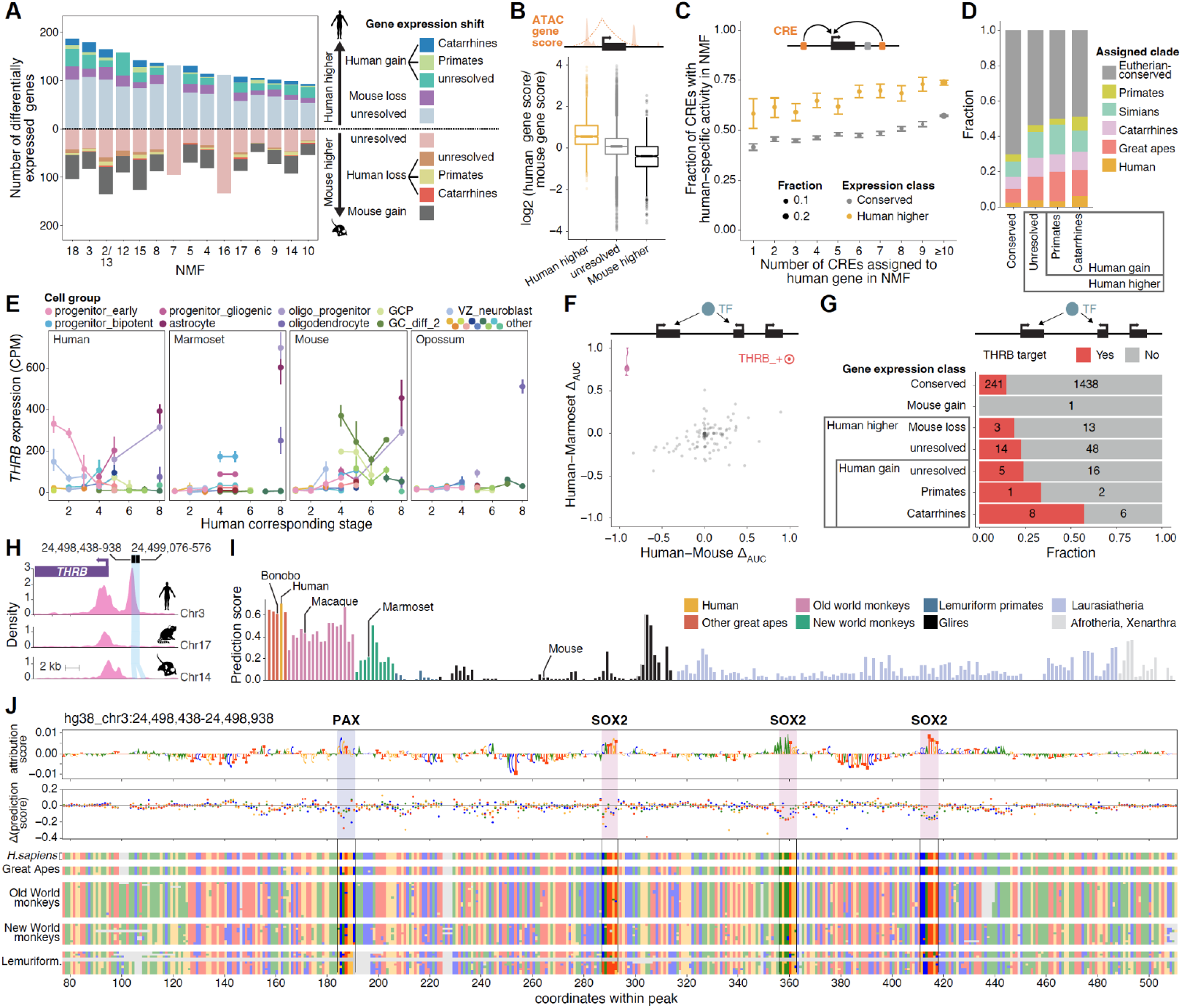
Gene expression shifts and their association with CRE innovations. (A) Evolutionary classification of genes with significantly higher expression in human or mouse in corresponding cell groups and stages across NMFs. (B) Local chromatin accessibility (ATAC-based gene scores) for genes with significantly higher expression in human or mouse across NMFs. Boxes represent the interquartile range and whiskers extend to extreme values within 1.5 times the interquartile range from the box. (C) Fraction of CREs with human-specific accessibility in the same NMF in which a gene is more highly expressed in human, stratified by the number of CREs linked to a gene. Genes with conserved expression in human and mouse are shown for comparison. Error bars indicate 5% and 95% confidence intervals, estimated by sampling genes with replacement 1000 times. (D) Fraction of CREs from different evolutionary clades based on DeepCeREvo’s predictions linked to genes with conserved or higher expression in humans. (E) Spatiotemporal expression of *THRB* in corresponding cell groups and developmental stages across mammalian species. Line denotes the mean and error bars indicate the range across biological replicates. CPM, counts per million. (F) Difference in TF activity (target-gene-based AUC score) between human, mouse and marmoset early cerebellar progenitors across 114 TF activators. (G) Fraction of THRB target genes with conserved and divergent expression in human early cerebellar progenitors. (H) Chromatin accessibility profiles around the TSS of *THRB* in early progenitors across species. Black boxes represent human CREs located ~3 kb upstream of the TSS. (I) DeepCeREvo’s predictions for sequences orthologous to hg38_chr3:24,498,438-24,498,938 across 227 mammals. (J) DeepExplainer and *in silico* mutagenesis profiles for NMF 5 (early progenitors), along with multiple sequence alignment (MSA) across 43 primates for hg38_chr3:24,498,438-24,498,938. A detailed MSA is provided in fig. S18.

When aggregating the chromatin accessibility profiles around genes to infer “gene scores”, we observed significantly higher accessibility around genes with higher expression in a given cell type and species (Fig. 7B and fig. S17, E and F). Besides providing orthogonal validation for these shifts, this observation also suggests that they are associated with changes in the chromatin accessibility landscape around these genes. Indeed, by integrating these data with our GRN analysis, we observed a significant enrichment (Likelihood ratio test, *P* < 10^−15^) of CREs with human-specific activity (compared to mouse) linked to genes with higher expression in humans relative to those with high expression in both species (Fig. 7C). These CREs include both human-specific CREs (i.e., regions not present or accessible in the mouse genome), as well as CREs with repurposed spatiotemporal activity. In total, we identified 8,691 CREs with human-specific activity (compared to mouse), associated with 987 genes with higher expression in humans in the context of the same NMF (table S16). For 23% of these CREs, we were additionally able to infer their evolutionary histories, using DeepCeREvo’s predictions (Fig. 7D).

As an example, we highlight the case of the thyroid hormone receptor beta (THRB), a transcription factor from the family of nuclear receptors with diverse functions in brain development (*60*). In the rodent cerebellum, THRB has been shown to regulate granule cell proliferation and migration, Purkinje cell dendritogenesis, myelination and Bergmann glia differentiation (*61*). While maintaining conserved expression in oligodendrocytes and astrocytes for at least 160 and 90 million years, respectively, the *THRB* gene gained expression in early cerebellar progenitors in the human lineage in the past 40 million years (Fig. 7E). Unlike most other TFs in our GRNs, THRB shows a major shift in transcription factor activity (AUC score) in human compared to marmoset and mouse progenitor cells (Fig. 7F), suggesting that the gain of *THRB* expression in the human lineage has propagated to the expression of downstream target genes. Indeed, THRB targets in the human GRN are significantly enriched amongst the genes we identified as differentially expressed in human cerebellar progenitors (Fisher’s exact *P* = 0.001, OR: 2.09), and especially amongst those that gained expression in the human lineage in the last 25 million years (Fisher’s exact *P* = 0.0003, OR: 7.65; Fig. 7G).

To investigate the regulatory basis of the *THRB* expression shift, we examined the evolutionary histories of CREs assigned to *THRB* in the GRN analysis. We found 5 CREs linked to *THRB* that are accessible in human but not mouse cerebellar progenitors. Of these, the two with the highest importance (hg38_chr3:24,498,438-24,498,938 and hg38_chr3:24,499,076-24,499,576) are about 3 kilobases upstream of the gene’s TSS and directly adjacent to each other, most likely representing a single regulatory unit. We did not observe any accessibility in this locus in mouse or marmoset (Fig. 7H) and our evolutionary classification based on the DeepCeREvo predictions suggests that this region gained accessibility in the ancestor of catarrhines approximately 25 MYA, though we also did observe high prediction scores in individual outgroup species (Fig. 7I). By investigating the sequence features driving high model predictions for cerebellar progenitors in hg38_chr3: 24,498,438-24,498,938, we identified two canonical (CATTGT) SOX2 motifs flanking a third deviant instance (CATTCT). All three motifs are highly conserved within Catarrhini species and in some new world monkeys, both in contrast to other mammalian clades and compared to their flanking sequences in catarrhines, suggesting that they are under functional constraint (Fig. 7J and fig. S18). Additionally, we identified an imperfect PAX motif (GCGTGGC), which was likely established in the common ancestor of the great apes. Collectively, our analyses suggest that sequence changes that occurred 25-40 MYA led to the gain of progenitor-specific CREs upstream of the *THRB* gene, resulting in a gene expression shift, which in turn propagated to the target genes of *THRB*. Thus, the roles of *THRB* in human cerebellum development likely extend beyond those described in rodents and may include evolutionarily novel functions in early progenitors.

## DISCUSSION

By combining cross-species single-cell multiomics datasets, machine learning models, and *ex vivo* measurements of enhancer activity, our study provides a comprehensive view of the evolution of the gene regulatory programs that define mammalian cerebellar cell types.

We reconstructed gene regulatory networks, and identified TFs with conserved activity in developing cerebellum cell types. Given their high network centrality and role in regulating genes with conserved cell type-specificity, we propose that these conserved TFs lie at the core of cerebellar cell type identities and can aid efforts to engineer these cell types *in vitro*. The presence of these core TF regulators explains why CRE sequence codes have remained conserved throughout mammalian evolution, despite the rapid turnover of individual CREs, especially around genes that have changed their expression during evolution. Our sequence-based model of CRE accessibility in cerebellar cell types, DeepCeREvo, learned these CRE codes, which allowed us to predict accessibility in unseen species and infer CRE evolutionary histories. In light of the numerous gene expression differences between species, the tracing of CRE histories is key for the prioritization of changes that might have been driven by positive selection. Notably, we linked recently emerged CRE sequence substitutions around the *THRB* gene to gains in expression and target gene activity of *THRB* in human progenitor cells. We thus hypothesize that the change in *THRB* expression could have contributed to evolutionary innovations of the cerebellum in the human lineage, such as the expansion of progenitor domains (*7*) or the shift in Purkinje cell developmental dynamics (*8*) which we estimate to have occurred in the past 40 million years.

Our study has several limitations. Our measurements of chromatin accessibility are only a proxy for CRE activity, which we tried to address by testing a selection of our predictions with enhancer reporter assays, albeit outside of their endogenous loci. Furthermore, our inference of regulatory interactions between TFs, CREs and their target genes is based on co-variance across cells and thus cannot discern association from causality. We attempted to tackle this by employing a mechanistically-inspired framework (e.g., by requiring the presence of the TF motif to link a TF to a CRE), and expect cross-species conservation to enrich for *bona fide* regulatory interactions. Finally, our analyses of gene expression and chromatin accessibility evolution were primarily focused on changes in cell type-specificity rather than quantitative levels of activity. This was a deliberate choice to limit the impact of known confounders such as differences in the quality of samples, genome assemblies, and annotations between species. We expect that the rapid improvement in machine learning methods will enable quantitative comparisons that explicitly account for these effects in future studies.

Despite its limitations, our study provides a new framework for studying the evolution of gene expression and its regulation that we anticipate to be broadly applicable across organs and species. While recent studies have used sequence-based models to explore the conservation of enhancer codes (*14, 34*) and we have applied them as a means to examine cell type homologies (*37*), this study is, to our knowledge, the first to reconstruct CRE evolutionary histories using such models. Our approach improves accuracy compared to sequence conservation, and allows highlighting individual sequences underlying the changes in CRE accessibility. As illustrated by the case of *THRB*, this facilitates the identification of CREs associated with gene expression shifts that can be further interrogated through functional studies aimed at understanding their roles in phenotypic evolution.

## Acknowledgements

We thank D. Odom, J. Zaugg, S. Anders, O. Stegle, B. Velten, M. Saraswat, P. Pruunsild, C. Mannens and all members of the Kaessmann and Aerts lab for discussions; K. Hall, A. Berenson, E. Wolff, A. Schrod, T. Becker, N. Umland, E. Renner, M. Toronyay-Kasztner, B. Nickel, T. Nath Varma, B. Crespo Lopez, and S. Krasemann for assistance, and the Joint MRC/Wellcome (MR/R006237/1) Human Developmental Biology Resource, Maryland Brain Collection at the Maryland Psychiatric Research Center (NIH NeuroBioBank), Chinese Brain Bank Center, Human Brain Tissue Bank at Semmelweis University, and P. Khaitovich for providing human samples.

## Author contributions

I.S., M.S. and H.K. conceived and organized the study. M.S. collected samples and performed experiments with support from J.S., C.S., R.F. and P.J.. I.S., T.Y., P.S.L.S. and M.S. analyzed data with support from N.T., I.I.T., N.H., C.B.G.-B., N.K. and E.L. C.D. and S.M. performed marmoset experimentation. R.B., S.L., M.P. and S.P. provided samples. S.M.P., L.M.K., M.C.-M., F.A., K.L., P.J. and K.O. provided critical discussions. N.T. developed the web application with input from I.S. and T.Y.. H.K. and S.M.P. provided funding. H.K. and S.A. supervised the study. I.S., M.S. and T.Y. drafted the manuscript, with critical review by P.S.L.S., M.C.M., H.K. and S.A. All authors provided feedback on drafts and approved its final version.

## Diversity, equity, ethics, and inclusion

We aimed for sex-balanced sampling. This study received support from a program fostering scientists from underrepresented backgrounds.

## Data and materials availability

All data generated in this study are freely available in the heiData repository https://heidata.uni-heidelberg.de/previewurl.xhtml?token=bea0c1aa-3110-4f3a-a98c-f10e65a407d0.

Processed data and DeepCeREvo’s predictions can be interactively explored at https://apps.kaessmannlab.org/cerebellum_genreg_evodevo_app.

Genome-wide chromatin accessibility profiles for human, marmoset and mouse are available as UCSC tracks:

https://genome-euro.ucsc.edu/s/ioansarr/hg38_cerebellum_tracks,

https://genome-euro.ucsc.edu/s/ioansarr/calJac4_cerebellum_tracks,

https://genome-euro.ucsc.edu/s/ioansarr/mm10_cerebellum_tracks.

Previously published datasets are available at https://doi.org/10.11588/data/QDOC4E (*8*) and Array Express (E-MTAB-9765 and E-MTAB-10533) (*38*).

Custom code is available at https://gitlab.com/kaessmannlab/mammalian-cerebellum-gene-regulation.

## METHODS

### Sample collection and ethics statements

Animal procedures were performed in compliance with national and international ethical guidelines and regulations, and were approved by the local animal welfare authorities at Heidelberg University Interfaculty Biomedical Research Facility (mouse: T-23/19, T-28/21, T-57/23, T-08/24, T-37/24), Lower Saxony State Office for Consumer Protection and Food Safety (LAVES; marmoset: #42502-04-12/0708 and #42502-04-16/2129). RjOrl:SWISS (RRID:MGI:5603077) adult, postnatal day P7, and time-mated pregnant mice (*Mus musculus*) were purchased from Janvier Labs (France). Adult mice were sacrificed by cervical dislocation and pups by decapitation. Common marmosets (*Callithrix jacchus*) were bred in a colony in the German Primate Center, Göttingen. Timed pregnancies were obtained from animals in which the gestational day (GD) was estimated by tracking the post-ovulatory increase in progesterone levels, and monitoring embryo growth by ultrasonography, as described previously (*62*). The embryos/fetuses were retrieved through a caesarean section procedure, ensuring the mother’s survival, performed by an experienced veterinarian under anaesthesia in sterile conditions, as described (*62*). All animals received appropriate analgesic and antibiotic treatment following surgery. The embryos/fetuses were weighed and measured (table S1). The rhesus macaque (*Macaca mulatta*) sample is from a colony in the German Primate Center, Göttingen. The bonobo (*Pan paniscus*) samples are from Lola Ya Bonobo Sanctuary Congo, Democratic Republic Congo. The macaque and bonobo individuals died for reasons other than their participation in this study (table S1). For bonobos, the entire cadavers were frozen at −20 °C, and the samples were dissected frozen.

The use of human samples included in this study was approved by an ERC Ethics Screening panel (associated with H.K.’s ERC Consolidator Grant 615253, OntoTransEvol) and ethics committees in Heidelberg (authorization S-220/2017), North East-Newcastle & North Tyneside (REC reference 18/NE/0290), London-Fulham (REC reference 18/LO/0822), Ministry of Health of Hungary (No.6008/8/2002/ETT) and Semmelweis University (No.32/1992/TUKEB). The human prenatal samples are from the MRC Wellcome Trust Human Developmental Biology Resource (HDBR; UK). The samples were donated voluntarily by women who had an elective abortion and provided written informed consent to donate fetal tissues for research. The prenatal samples had normal karyotypes, and were categorized into specific Carnegie stages or weeks post conception (wpc) based on their external physical appearance and measurements. The human postnatal samples are from the University of Maryland Brain and Tissue Bank of National Institutes of Health NeuroBioBank (USA), Chinese Brain Bank Center (CBBC) in Wuhan, and Lenhossék Human Brain Program, Human Brain Tissue Bank at Semmelweis University (Hungary). Written informed consent was obtained from donors or their families for the utilization of tissues in research. All postnatal samples were sourced from healthy, non-affected individuals classified as normal controls by the respective tissue bank.

Cerebella or its fragments were dissected as previously described (*8, 38*). If available, samples from both sexes were used for data production. To determine the sex of the developing animals, tail samples were collected and subjected to PCR genotyping using MyTaq Extract-PCR Kit (Meridian Bioscience) and Y-chromosome-specific primers (mouse: 5’-TCATGAGACTGCCAACCACAG-3’ and 5’-CATGACCACCACCACCACCAA-3’) or primers targeting *DDX3* (marmoset: 5’-GGWCGRACTCTAGAYCGGT-3’ and 5′-GTRCAGATCTAYGAGGAAGC-3′), which is found on both the X and Y chromosomes, with variants of different lengths. Of note, due to cellular chimerism in twin marmosets, even females may exhibit a weak male-specific band if their co-twin is male. (*62*). All samples with details about their collection are listed in table S1.

### Primary cultures of mouse granule cells

Cerebella from P7 RjOrl:SWISS mice were used in all experiments. For multiome data production, pups were divided by sex to facilitate multiplexed profiling as described in the section *Preparation of nuclei and sample quality control*. Sex was initially estimated based on the presence (male) or absence (female) of dark pigmentation on the perineum, and morphology of the developing gonads (*63*). Assignments were subsequently confirmed with 100% accuracy through PCR-based genotyping as described above. Granule cells were cultured essentially as described previously (*64–66*). P7 cerebella were dissected under the microscope, meninges was removed, and cells were dissociated using Papain Dissociation System Kit (Worthington) according to manufacturer’s instructions with 2.5 ml papain solution applied to 4-6 cerebella. After papain treatment and one-step discontinuous density gradient centrifugation (100 g 4 min) in albumin-ovomucoid inhibitor solution (following the kit’s protocol), cells were resuspended in 2 ml EBSS (Worthington) supplemented with 0.5 mg/ml DNase (Sigma-Aldrich) and 5 g/l glucose (Thermo Fisher Scientific), strained using 70 μm MACS SmartStrainers (Miltenyi Biotec), and subjected to two-step discontinuous density gradient centrifugation (2000 g 12 min with lowered acceleration/deceleration ramps) in 35% / 60% Percoll (5 ml + 5 ml; Sigma-Aldrich) acidified to pH 7.4 with HCl. Cells at the interface between the 35% and 60% Percoll were collected (ca 3 ml), 3 volumes of HBSS with 6 g/l glucose was added and cells were pelleted at 1100 g 5 min. Cells were resuspended in growth medium containing Neurobasal Plus Medium supplemented with 100 U/ml penicillin-streptomycin, 2 mM GlutaMAX, 4.5 g/l glucose, B-27 Plus (all Thermo Fisher Scientific), SPITE and Linoleic Acid-Oleic Acid-Albumin supplements (Sigma-Aldrich), and 0.16 mg/ml N-Acetyl-L-cysteine (Sigma-Aldrich). Cells were pre-cultured for 2 times 45 minutes on uncoated cell culture plates, non-adherent cells were collected and live cell counts were estimated using Trypan Blue and Countess (Thermo Fisher Scientific). Cells were plated on dishes coated with poly-D-lysine (0.1 mg/ml, Thermo Fisher Scientific) and Matrigel Growth Factor Reduced Basement Membrane Matrix (Corning) diluted 1:75 in HBSS (Thermo Fisher Scientific) at a density of 2-3 × 10^5^ cells/cm^2^. At DIV (days in *vitro*) 0-3, 200 nM InSolution Smoothened Agonist (SAG; Sigma-Aldrich) was added to the growth medium to support proliferation of granule cell progenitors, and full medium changes were performed daily. At DIV3, SAG was omitted and half of the medium was changed at DIV5. Cells were collected for further assays at DIV3 and DIV6. In some experiments, cells cryopreserved in CryoStor CS10 (Stemcells Technologies) after pre-culturing, were used.

### Preparation of nuclei and sample quality control

The nuclei were prepared as described (*8*) and used as input for the production of single-nucleus libraries (table S1). For primary granule cells, nuclei from male DIV3 cells and female DIV6 cells were mixed in a 1:1 ratio and demultiplexed *in silico* as described in the section *Processing and quality control of single-nucleus multiome sequencing data*. Neighbouring tissue or total or cytoplasmic fractions from the preparations were used for RNA extraction to monitor sample quality (table S1). Total and cytoplasm extracts were mixed with 40 mM DTT-supplemented RLT buffer (Qiagen) and 100% ethanol at 2:7:5 ratio, and RNA was purified using the RNeasy Micro kit (Qiagen). The same kit was used for RNA extractions directly from tissues. RNA quality numbers (RQN) were determined on Fragment Analyzer (Advanced Analytical). Most samples had an RQN value above 7, except for a few human postnatal samples (table S1).

### Bulk RNA-sequencing

RNAs were extracted directly from tissue fragments, total lysates, or fractions of nuclei and/or cytoplasm (table S2) using RNeasy Micro or Mini kits (Qiagen). Most extracted RNAs had an RQN value above 7, except for two bonobo samples (table S2). Bulk RNA-sequencing libraries were prepared with the NEBNext Ultra II kit (New England Biolabs) at the Deep Sequencing Core Facility of Heidelberg University. Qubit Fluorometer (Thermo Fisher Scientific) was used to estimate DNA concentrations, and the average fragment size was determined on Bioanalyser 2100 (Agilent). Libraries were sequenced on Illumina NextSeq 550 using High Output Kit v2.5 with 150 cycles (Illumina) and the following setup: 159 cycles for Read 1 (cDNA), 8 cycles for i7 index (sample index).

### Single-nucleus RNA and ATAC sequencing

Our study includes both previously reported single-nucleus libraries (*8, 38*) as well as newly generated libraries (table S1). Human, mouse, and opossum cerebellum data come from separately produced snRNA-seq and snATAC-seq libraries; whereas marmoset, macaque and bonobo cerebellum, and mouse cultured granule cell data are from multiome libraries, where both data types are profiled from the same cells. For mouse and human, we selected previous datasets and produced new datasets with the aim to match the separately produced snRNA-seq and snATAC-seq datasets as closely as possible. Dataset pairs were set up based on the following order of priority: the same nuclei preparation, the same sample, the same litter, and, lastly, the same sex and/or developmental stage (table S1).

Our previously reported snRNA-seq libraries of human, mouse and opossum were prepared using Chromium Single Cell 3’ Reagent kits (10x Genomics; v2 or v3 chemistry) (*8*). Our previously reported snATAC-seq libraries of mouse and opossum were prepared using Chromium Single Cell ATAC Reagent kits (10x Genomics; v1 for mouse and v1.1 for opossum) (*38*). Production of new single-nucleus datasets in this study essentially followed similar procedures as used in our previous studies. Specifically, Chromium Single Cell 3’ Reagent kits (v3 or v3.1) and the Chromium Controller instrument (10x Genomics; RRID:SCR_019326) were used to generate new snRNA-seq libraries for human (n=6) and mouse (n=3). Manufacturer’s protocols were followed, with 12 PCR cycles used for cDNA amplification. Newly added snATAC-seq libraries for human (n=24) and mouse (n=3) were prepared using Chromium Single Cell ATAC Reagent kits (v1 or v1.1). Single-nucleus multiome datasets for marmoset (n=17), bonobo (n=3), macaque (n=3) and mouse cultured granule cells (n=1, nuclei from male DIV3 cells and female DIV6 cell mixed) were produced using Chromium Next GEM Single Cell Multiome ATAC + Gene Expression Reagent kit, with 7 PCR cycles used for cDNA amplification. In most of the experiments, 15,000-17,000 nuclei were loaded per channel (range 1000-25,000; table S1). Libraries were quantified on Qubit Fluorometer (Thermo Fisher Scientific) and the average fragment size was determined on Fragment Analyzer (Advanced Analytical).

Libraries were sequenced on NextSeq 550 using High Output v2.5 kits (Illumina) with 75 (separate snRNA-seq and snATAC-seq libraries) or 150 (multiome libraries) cycles. For the separate snRNA-seq libraries the setup was: 28 cycles for Read 1 (cell barcode), 8 cycles for i7 index (sample index), 56 cycles for Read 2 (cDNA). For the separate snATAC-seq libraries the setup was: 34 cycles for both Read 1 and 2 (gDNA), 8 cycles for i7 index (sample index), 16 cycles for i5 index (cell barcode). For the multiome snRNA-seq libraries the setup was: 28 cycles for Read 1 (cell barcode), 10 cycles for both i7 and i5 indices (sample index), 90 cycles for Read 2 (cDNA). For the multiome snATAC-seq libraries the setup was: 50 cycles for both Read 1 and 2 (gDNA), 8 cycles for i7 index (sample index), 16 cycles for i5 index (cell barcode). A custom recipe that includes 8 dark cycles on i5 was used for the sequencing of multiome snATAC-seq libraries.

### Genome and transcript isoform annotations

For human and mouse, which have been extensively studied and are expected to have mostly complete transcript annotations, we used annotations from Ensembl v92 for the hg38 and mm10 assemblies, respectively (*67*). For gene expression analyses, we excluded non-coding genes overlapping coding genes in the same strand to minimize the loss of reads assigned to more than one gene in full-transcript (including introns) counting mode. For chromatin accessibility analyses, we generated custom ArchR (v1.0.2) annotations based on Ensembl v92, only considering protein-coding genes, as we observed that the inclusion of non-coding genes led to reduced and more noisy gene score estimates.

For species with less complete transcript annotations, such as marmoset, rhesus macaque and bonobo, we used bulk RNA-sequencing data, which we generated in parallel to our single-nucleus experiments, to extend the existing reference annotations (Ensembl v92 for macaque and bonobo, RefSeq annotation calJac4 for marmoset), akin to our previous work (*68, 69*). Bulk RNA-sequencing libraries were demultiplexed using bcl2fastq (v2.20) and aligned to the respective genome assemblies (calJac4 for marmoset, Mmul_8.0/rheMac8 for macaque and panPan1 for bonobo) using STAR (v.2.7.9.a) (*70*). BAM files from the same developmental stage (biological or technical replicates) were merged to improve the sensitivity of new transcript isoform identification. We then used Stringtie (v1.3.3) (*71*) (parameters: -f 0.2 -m 200 -a 10 -j 3 -c 2.5 -v -g 10 -M 0.5) to assemble models of transcripts expressed in each developmental stage. For marmoset, we used cuffmerge (v.2.2.1) from the cufflinks package (*72*) with default settings to combine the individual annotations from each developmental stage into a single consensus annotation. We then used cuffcompare from the same package to combine the new annotation with the previously available reference annotations. As for human and mouse, we excluded non-coding genes overlapping coding genes in the same strand for the annotation used for gene expression counting, and only considered protein-coding genes to estimate gene scores in ArchR (v1.0.2) (*73*).

### Processing and quality control of snRNA-seq data

We applied uniform processing and quality control to all previously published (*8*) and newly sequenced snRNA-seq libraries for human and mouse. After demultiplexing data from newly sequenced libraries with bcl2fastq (v2.20), we used STARsolo (v.2.7.9.a) (*70*) to align reads to the reference genomes (hg38 for human, mm10 for mouse) and to count reads in exons and full-length transcripts, allocating multimapping reads based on an expectation-maximisation algorithm (−-soloMultiMappers EM). We then applied Gaussian mixture models with two groups on the distributions of UMI counts (full-length counting mode) and the fraction of intronic reads, as implemented in the R package mclust (v5.4.7), to identify barcodes corresponding to cells. To this end, we only considered barcodes included in the group with higher values for both metrics, and additionally required the number of UMIs to be at least 40% of the median across all putative cells in that sample. We used scrublet (v0.2.3) (*74*) to remove the 10% of cells with the highest doublet score from each sample, as well as cells with more than 2.5 times the median number of UMIs (full-length counting) in that sample.

For each sample, we used Seurat (v4.0) (*75, 76*) to regress out cell cycle scores (only correcting for the difference between S and G2/M phases), applied SCTransform and projected the data into 50 principal components, which we utilized for low-resolution Louvain clustering (resolution=0.2). We then used these clusters as input to SoupX (v1.5.2) (*77*) to correct the expression of transcripts associated with ambient RNA or cellular debris. Contamination estimates appeared much higher when considering exonic counts, suggesting that most contaminating transcripts are already spliced. Thus, we only corrected exonic counts, and subsequently reconstructed full-length expression counts by adding the corrected exonic counts to the uncorrected intronic count values. Additionally, we limited the correction to genes estimated to contribute more than 0.05% of the total contamination to avoid introducing noise in the expression of genes for which the background contamination level could not be reliably estimated. We then used the corrected full-length count values to repeat the Seurat analysis described above (cell cycle regression, SCTransform, PCA). Next, we integrated samples from the same developmental stage using the Seurat function IntegrateData() based on the SCT-corrected counts in the 3,000 most highly variable features across biological replicates. These stage-wise integrated objects were used for integration with the corresponding snATAC-seq datasets, as described below.

### Processing and quality control of snATAC-seq data

Data from newly sequenced snATAC-seq libraries were demultiplexed and converted to fastq format using cellranger-atac mkfastq (v1.1.0) (*78*). The command cellranger-atac count (v1.1.0) was used to align reads to the reference genomes (hg38 for human, mm10 for mouse) and generate position-corrected tabular fragment files. Fragment files from both new and previously published (*38*) snATAC-seq datasets were used as input to ArchR (v1.0.2) (*73*). Barcodes corresponding to cells were identified by applying Gaussian mixture models (mclust, v5.4.7) with two groups on the distributions of the number of fragments and TSS enrichment scores, estimated using ArchR (v1.0.2). Only barcodes assigned to the group with higher values for both metrics, and additionally having at least 2,500 fragments and a minimum TSS enrichment of 2.5 were considered for downstream analyses. ArchR (v1.0.2) was then used to estimate doublet scores per cell. The top 10% of cells with the highest doublet scores in each sample as well as cells with more than 2.5 times the median number of fragments in that sample, were removed.

Samples from the same stage were then jointly analyzed using ArchR (v1.0.2), counting accessibility in 500 bp-wide windows and inferring gene scores (*73*). Single-cell chromatin profiles were projected into 50 latent dimensions based on an iterative LSI with gradually increasing clustering resolution (0.1, 0.2, 0.4, 0.8). These dimensions were then corrected using Harmony (v.0.1.0) (*79*) to facilitate integration between biological replicates.

### Processing and quality control of single-nucleus multiome sequencing data

Single-nucleus multiome (ATAC and gene expression) libraries for marmoset, rhesus macaque, bonobo, and mouse primary granule cells were demultiplexed and processed using cellranger-arc (v2.0.1). The alignment and counting of gene expression libraries was performed independently using STARsolo (v.2.7.9.a) (*70*) allocating multimapping reads based on an expectation-maximisation algorithm (−-soloMultiMappers EM) for consistency with the processing of human and mouse snRNA-seq libraries. Barcodes corresponding to cells were identified based on a combination of the metrics we used for independently processed snRNA-seq and snATAC-seq libraries. We used Gaussian mixture models (mclust, v5.4.7) with two groups on the distributions of the number of UMIs (full-length counting), the fraction of intronic reads, the number of ATAC fragments, and the TSS enrichment scores (as estimated by ArchR, v1.0.2). Only barcodes corresponding to the group with higher values for all four metrics, as well as having at least 40% of the median number of UMIs across putative cells in that sample, were considered as cells.

We used scrublet (v0.2.3) and ArchR (v1.0.2) to estimate doublet scores for the gene expression and chromatin accessibility modalities, respectively. To facilitate integration between these two metrics, we standardized these scores within each sample as Z-scores and additionally estimated a consensus doublet score by taking the mean of the two scores for each barcode. We then removed barcodes that ranked in the top 10% when considering the consensus doublet score or in the top 5% when considering the doublet score in either modality. Gene expression counts were corrected for the effect of ambient RNA or cellular debris using SoupX (v1.5.2) on the exonic counts, focusing on the genes with the highest contribution (more than 0.05%) to the estimated soup, as described above for snRNA-seq datasets. To demultiplex the data from a library in which male DIV3 and female DIV6 primary granule cells were mixed, we used the R package cellXY (v0.99.0). Based on gene expression profiles in each cell, we identified 34 male/female doublets (0.8%) using the findMfDoublet() function and removed them from subsequent analysis. We then assigned the sex of each cell using the classifySex() function, identifying 1,201 male DIV3 cells (30.1%), 2,784 female DIV6 cells (69.8%), and 2 unassigned cells (0.05%; excluded from further analyses).

### Integration across developmental stages and cell type annotation based on gene expression

We used liger (v0.5.0) (*80*) to integrate all cells from the same species based on their gene expression profiles. We first identified highly variable genes using a variance threshold of 0.1 within each sample. We then considered genes identified as highly variable in at least two samples (since every developmental stage had at least two biological or technical replicates). We additionally excluded genes with extreme sex-bias (more than 0.8 Pearson’s correlation coefficient with *XIST* across all cells in the dataset), a set of manually curated cell cycle-related genes provided by the Linnarsson lab (*40*), as well as a set of 25 genes recently reported as associated with activation-signatures induced by differences in sample preparation (*81*). After creating a single liger object for all samples from the same species and manually setting the highly variable genes as described above, we used optimiseALS() with lamda=5 to project the cells into 100 components for human and 75 components for mouse and marmoset. These embeddings were used for constructing a neighbor graph (20 NNs), Louvain clustering at a high resolution (3.0) and UMAP projection (metric = “cosine”, min.dist = 0.1, n.neighbors = 20) as implemented in Seurat (v4.0).

For human and mouse, 71% and 78.8% of the cells in the snRNA-seq dataset were assigned a cell type and state annotation from our previous work (*8*). To annotate newly added cells, we applied a neighbor-voting procedure, coupled with manual curation of clusters with a fraction of unannotated cells. First, for each newly added cell, we considered its 20 nearest neighbors obtained from the global liger embedding. If at least 50% of these neighbors for which an annotation was available (and with a minimum of 5 cells) shared the same cell type label, the new cell was also assigned that label. Based on this approach, we were able to annotate 18,979 (14.5%) human and 13,720 (11.7%) mouse cells. Using previously annotated cells as a control, we observed high concordance in cell type labels (98% and 96% for human and mouse respectively; 95% and 88% at the precisest level of cell state/subtype annotation with many of the mismatches driven by adjacent differentiation states, such as GC_diff_1 and GC_diff_2). To account for the possibility of new cell types or states being present in our newly profiled samples, we next considered clusters with a high fraction of cells (more than 50%) that remained unannotated. These clusters were annotated on the basis of the expression of known marker genes and include oligodendrocyte and deep nuclei neurons from white-matter enriched adult samples. At this stage, we were able to annotate an additional 2,323 (1.8%) human and 8,542 (7.3%) mouse cells. Cells that remained unannotated at this stage were subjected to a final round of neighbor-voting. At this stage, for each cell we identified the 20 nearest neighbors that already had an annotation (i.e., which might not be in the 20 neighbors considering all cells). If more than 50% of these neighbors had the same label at the precisest level of our cell type annotation, we assigned this label to the newly profiled cell. At this final stage, we were able to annotate 16,391 (12.6%) human and 1420 (1.2%) mouse cells. All remaining cells were not annotated and excluded from downstream analyses. In total, we were able to provide a confident annotation for 99.6% and 98.6% mouse cells profiled with snRNA-seq.

To annotate the 79,702 cells in the newly generated marmoset dataset, we used Seurat (v4.0) to predict cell type and state labels using the mouse and human cell type annotations as a reference. For each pairwise integration, we used 1:1 orthologous genes, highly variable in at least two samples in both species (4,532 and 3,076 genes for integrations with human and mouse, respectively). We used the function TransferData() with k=30 and weight.reduction=“cca” to transfer cell type and state labels from each reference annotation (human, mouse) to marmoset cells. We observed high concordance in the predictions obtained using the human or mouse reference annotation (93% agreement at the level of broad cell types, with the most prominent mismatches representing developmental transitions, such as from progenitors to ventricular zone neuroblasts). Given this agreement, we relied primarily on the predictions obtained by using the mouse as a reference, the species that we were previously able to annotate at the highest granularity (*8*). We additionally complemented the mouse predictions with human predictions for cell type labels that were not distinguished in the mouse, such as the progenitors of the anterior ventricular zone. To also accommodate the detection of cell types and states that are absent from both the human and mouse annotations, we additionally performed Louvain clustering (resolution: 3.0) followed by subclustering of individual cell type groups (broadly corresponding to astroglia, ventricular zone-derived neurons, rhombic lip-derived neurons and other/non-neural cell types). We manually curated these clusters and modified cell type labels in the few cases for which we observed a mismatch between the predicted label and the actual label as inferred by the expression of cell type-specific marker genes. This procedure allowed the identification of rare populations of ependymal progenitors, contaminating cells from the midbrain and the lower brainstem regions, as well as the removal of putative residual doublets that were not removed during the initial processing of the data. In total, we were able to provide confident cell type and state annotations for 76,508 (96%) marmoset cerebellar cells profiled for both gene expression and chromatin accessibility.

For rhesus macaque, bonobo, and mouse primary granule cells, we followed the same approach as for marmoset using the mouse cell type annotation as a reference, with the following modifications to account for the overall lower complexity of the datasets which only covered a limited number of developmental stages and cell states. We selected 1:1 orthologous genes that were highly variable in at least two samples in both species, yielding 1,403 genes in macaque and 1,129 genes in bonobo. For the mouse granule cell culture, we intersected highly variable genes from both *in vitro* and *in vivo* datasets, resulting in 1,641 genes.. Using these genes as features, we transferred cell type and state labels from mouse reference annotations using weight.reduction=“pcaproject”, as this method is efficient when applying large reference datasets or classifying query datasets with relatively homogeneous cell populations (*75*). We manually curated these transferred labels by inspecting the expression of cell type-specific marker genes when necessary. For rhesus macaque, we further refined the cell type and state annotations by using a k-nearest neighbor graph (k = 20). Each cell was assigned the most common label among its neighbors, provided that at least 5 cells shared the same annotation and represented more than 50% of the annotated neighbors.

### Integration between paired snRNA-seq and snATAC-seq data

For human and mouse, we used Seurat (v4.0) (*75, 76*) to integrate the snRNA-seq and snATAC-seq modalities. Starting from the stage-wise integrated embeddings for each modality (generated using Seurat v4.0 and ArchR v1.0.2 as described above), we identified the 5,000 most highly variable genes in the snRNA-seq dataset. These were used to construct transfer anchors between the reference set (snRNA-seq data) and the query (snATAC-seq data). We used the function TransferData() with k.weight=30 in the snATAC-seq LSI embedding to weigh the predictions. We transferred labels for cell types and states and additionally imputed coordinates for the stage-wise (Seurat v4.0) and global (liger 0.5.0) integrated snRNA-seq embeddings. This imputation step allowed us to co-embed cells profiled with snRNA-seq an snATAC-seq.

### CRE identification and characterization

We used ArchR (v1.0.2) to identify peaks of open chromatin (as proxy for putative CREs) within each species in a cell type-specific and replicate-aware manner. We grouped cells with the same cell type label (at the most precise level of our annotation, i.e., also considering subtypes and developmental states) and from the same replicate into pseudobulks using the function addGroupCoverages(). We required at least 50 cells in each pseudobulk setting the sample ratio (i.e., fraction of cells from other replicates that can be sampled to complete the required number for groups that have fewer than 50 cells) to 80%. We constructed between 2 and 10 pseudobulks per cell type, allowing each pseudobulk to contain up to 500 cells and 50 million fragments to cap the contribution of very abundant cell types. We then used the function addReproduciblePeakSet(), which internally calls MACS2 (v2.1.2) (*82*), to identify peaks of open chromatin in each group. We used parameters “--shift -75 --extsize 150 --nomodel --call-summits --nolambda --keep-dup all -q 0.01”, retaining up to 200,000 peaks per group or up to 1,000 peaks per cell. ArchR collapsed the peak annotations from each pseudobulk group into a single consensus annotation based on its iterative overlap peak merging procedure, which retains the most significant summit around a 500 bp region discarding any less significant overlapping peaks (*73*). Only peaks that could be independently detected in at least two pseudobulk groups (i.e., reproducible) were retained in the final peak annotations.

Orthologous CREs between species were identified based on reciprocal syntenic alignments in a pairwise manner, as described previously (*38*). Briefly, we used liftOver with -minMatch=0.1 -multiple -minSizeQ=50 -minSizeT=50 to identify syntenic regions from species A to species B. These regions were then overlapped with peak annotations in species B using bedtools intersect (v.2.28) (*83*). Peaks with reciprocal and unique matches between species were considered orthologous. Additionally, since we used a fixed 500 bp width for the peak annotations in each species, we considered cases where one peak from species A matched to two peaks from species B that were up to 500 bp from each other. In these cases, we retained the peak with the highest overlap as the ortholog in species B. Additional one-to-many and many-to-many matches were excluded from downstream analyses.

### Developmental correspondences

We applied a dynamic time-warping algorithm, as implemented in the R package dtw (v1.22.3) on four different metrics of dissimilarity, incorporating both gene expression and chromatin accessibility modalities, to reevaluate our previously reported correspondences between human and mouse developmental stages (*8*) and to infer corresponding stages for the development of the cerebellum in marmoset.

First, we used canonical correlation analysis (CCA) as implemented in Seurat (v.4.0) to transfer snRNA-seq developmental stage annotations from one species to another based on 1:1 orthologous genes with highly variable expression in at least two samples in each species. Smoothing across the 30 nearest neighbors, we predicted developmental stage annotations from the reference species for each cell from the query species. Then, for each developmental stage in the query species, we calculated the mean prediction score across all cells for every developmental stage in the reference species. Finally, we subtracted these scores from 1 to convert the estimates into a metric of dissimilarity.

As a second metric, we considered similarities in cell type composition. For each species and for each developmental stage, we estimated the fraction of cells belonging to every possible cell type (at the second level resolution, i.e., cell states). Then, we used the fractions of cells belonging to each label to estimate Manhattan distances between every possible combination of developmental stages from the two species.

For the third metric, we directly compared gene expression profiles of 1:1 orthologous genes between species. We aggregated gene expression counts (estimated using full-length counting mode) across all cells from a sample, scaled by counts per million (CPM) and applied quantile normalization as implemented in the R package preprocessCore (v.1.54). We averaged expression profiles across biological and technical replicates from the same species and developmental stage. Considering genes reaching at least 25 CPM in at least one developmental stage, we identified within each species the 5000 genes with the highest temporal variance based on their variance/mean ratio. We then converted expression values into Z-scores (i.e., measuring the time-specificity of their expression). Finally, we estimated Spearman’s correlations across species and developmental stages using the Z-score standardized expression values of 1:1 orthologous genes with temporally variant expression in both species.

For the fourth metric, we compared chromatin accessibility profiles of 1:1 orthologous CREs between species. We used the same procedure as with gene expression, but this time considering CREs with at least 1 CPM in at least one developmental stage (the feature space for CREs is approximately 25 times larger than for protein-coding genes). We estimated Spearman’s correlations for orthologous CREs belonging to the 100,000 most highly variable features in both species.

In the rhesus macaque and bonobo, we applied the same methodology as used for the mouse, marmoset, and opossum to infer developmental correspondences to humans. However, due to the limited number of developmental stages sampled in these species, we omitted the direct prediction of developmental stages (first metric) and modified the computation of the third and fourth metrics. Since the limited number of stages in macaque and bonobo limited our ability to detect highly variable features in these species, we focused exclusively on the 5,000 genes and 100,000 CREs exhibiting the highest temporal variance in human. Additionally, since within-species scaling is only reliable in datasets with multiple developmental stages and cell types, we calculated Spearman’s correlations using CPM instead of Z-score normalized CPM values.

While inferring developmental correspondences between marmoset and the other species in the dataset, we observed that the sample dated as GD80 (sm043) appeared to be substantially less developed than expected, showing the highest similarity to samples from GD73. This apparent developmental delay is also supported by our morphological measurements, as this embryo/fetus weighted 40% of its littermate (for which we were unfortunately unable to generate data due to a technical failure). Developmental differences between marmoset embryos/fetuses are not uncommon, with 24% of twin pregnancies resulting in only one birth (*84*). Given this uncertainty about the developmental staging of the GD80 embryo/fetus, for which we also lack a biological replicate, we decided to exclude this sample from all comparative analyses which require the confident identification of a developmental stage and of its corresponding stages in other species. However, reasoning that despite its developmental delay this sample might still capture aspects of normal development in earlier stages, we decided to provide the molecular profiles and annotations of these cells for further exploration by the scientific community.

### Gene regulatory network (GRN) inference

We constructed gene regulatory networks (GRNs) for human, mouse, and marmoset using the SCENIC+ pipeline (*43*) on metacells generated by aggregating single cells in similar cell states as described below. There were three primary motivations for aggregating single cells into metacells. Firstly, since our gene expression and chromatin accessibility measurements were obtained independently for human and mouse, linking profiles at the single-cell level is unrealistic. Using metacells allows us to aggregate at the level of granular cell states, which can be linked with much higher confidence across modalities. Secondly, even for species for which gene expression and chromatin accessibility were profiled from the same cell (e.g., marmoset), we found that metacell aggregation improved the analysis by reducing the sparsity of single-cell measurements. Finally, aggregating cells into metacells improved the computational efficiency of the analysis, allowing us to perform some of the more computationally intensive steps of the SCENIC+ pipeline.

The main steps of our GRN inference approach are described below.

#### 1. Motif enrichment analysis

To infer putative TF binding sites in our CREs, we relied on TF motif enrichment as implemented in pycisTarget (*43*). To this end, we first obtained sets of co-accessible CREs for each species using two complementary approaches, differential accessibility and topic modeling.

To identify CREs specific to different cell states, we performed differential accessibility analysis across cell states (annotation level 3) using the pycisTopic function find_diff_features(). Each cell state was contrasted against all others, using the primary fragment matrix with standard parameters. Differentially accessible regions (DARs) were identified based on a *P*-value threshold of 0.1 and a log fold-change threshold of 0.32.

To additionally capture more complex patterns of co-accessibility (for example CRE accessibility shared across cell states or changing along developmental trajectories), we also considered a cell annotation-free approach, topic modeling, as implemented in pycisTopic. For each species, we considered models with a varying number of topics, from 30 to 140 in intervals of 10. Based on several metrics incorporated within the package, we ascertained the optimal topic count as 130 for human, 100 for mouse, and 140 for marmoset. CREs associated with each topic were identified by binarizing the topic-region matrix using the Otsu method.

For each species and for each set of co-accessible CREs (DARs and binarised topics) we estimated the enrichment of more than 49,000 TF motifs collapsed into more than 8,000 clusters (https://resources.aertslab.org/cistarget/motif2tf/). We used the v10 motif annotations for human and mouse, and considered the human motif annotation for marmoset. To ensure that all our CREs were considered for TF motif enrichment, we constructed custom cisTarget databases for each species using the create_cistarget_databases code snipped from https://github.com/aertslab/create_cisTarget_databases. We utilized both the cisTarget and differential enrichment of motifs (DEM) methodologies using default parameters.

#### 2. Aggregating single cells into metacells

We aggregated cells into metacells based on the stage-specific RNA and ATAC co-embeddings for human and mouse, as outlined in the section *Integration between paired snRNA-seq and snATAC-seq data*. We randomly selected 50% of the cells as seeds and aggregated up to 40 cells around each seed in the embedding, identified via the K-Nearest Neighbor algorithm, for both the RNA and ATAC modalities to create metacells. Aggregates that did not consist of at least two-thirds of their cells (i.e., a minimum of 26 cells) with a consistent cell-type label across the embeddings were discarded. To minimise redundancy among metacells, we only considered metacells with less than 30% overlap in their single cells. Finally, we scaled the gene expression and chromatin accessibility profiles of each metacell to 10,000 counts and 100,000 counts respectively. Genes and CREs expressed/accessible in less than 0.5% of metacells were excluded from the analysis.

For marmoset, we followed the same aggregation, filtering and scaling procedure, omitting the integration step between snRNA-seq and snATAC-seq data, as both modalities were profiled from the same cell. Instead, we aggregated neighbors using the RNA embedding for each developmental stage. Altogether, our dataset consisted of 6,852 metacells for human, 5,153 for mouse, and 3,604 for marmoset. Importantly, the relative representation of cell states was largely preserved between the single cell and metacell datasets. UBC_diff, UBC_defined, and isth_N_diff were not considered in the marmoset dataset due to their low abundance.

#### 3. SCENIC+ analysis

Utilizing the aggregated multiome profiles, we calculated CRE-gene and TF-gene associations, using Gradient Boosting Machine regression with default parameters. Based on the TF motif enrichment, CRE-gene and TF-gene links, we constructed the GRN for each species using the build_grn function with default parameters. For each eRegulon, enrichment scores (AUC: area under the recovery curve) of target genes and CREs per cell were calculated using the per-cell rankings of gene expression and imputed accessibility scores, respectively. In addition to the default filtering step, we required eRegulons to have at least 50 target genes and to show correlations of at least 0.4 between gene- and CRE-based AUCs, as well as between TF expression and gene-based AUC. This resulted in 420 eRegulons for human, 324 for mouse, and 347 for marmoset.

### Benchmarking mouse GRN

We assessed the quality of our GRNs at three levels: TF-CRE, CRE-gene and TF-gene links. For TF-CRE links (i.e., the ability of our model to infer the binding of a TF on a CRE), we relied on previously published ChIP-seq data for TFs detected in the mouse GRN within cell types observed during cerebellum development (*85–92*) (table S7). We downloaded the fastq files from each ChIP-seq experiment from the NCBI Sequence Read Archive (SRA). Adapter trimming, low-quality sequence removal, and quality control were performed using Cutadapt and FastQC, respectively, both of which are incorporated within Trim Galore (v0.6.6) (*93*). The trimmed and filtered reads were then aligned to the mouse mm10 (GRCm38) assembly using BWA-MEM (*94*). For peak calling we used MACS3 (v3.0.1) with default parameters (*82*). ChIP-seq datasets for Mef2a, Mef2d, and Olig2, in which fewer than 1,000 peaks were called, were excluded from subsequent analyses. We verified that motifs of the TFs under investigation were enriched in the ChIP-seq peaks by employing motifmatchR (*95*) in conjunction with the JASPAR2022 database (*96*). For each ChIP-seq experiment, we used rtracklayer (*97*) and BRGenomics (*98*) to calculate the coverage in regions extending ±500 bp from the target CREs of Atoh1, Nfia, Nfib, Nfix, Nkx2-2, Spi1, and Zic2 activators in the mouse GRN, where TF and target gene expression are positively correlated. To account for differences in the number of target CREs between TFs, we downsampled CREs to match the smallest number of target CREs, which was for Nkx2-2 (n = 928). The coverage was then smoothed by taking the rolling mean over 100 bp windows.

We next sought to assess the quality of the CRE-gene links identified by SCENIC+. Since most enhancer-promoter interactions tend to occur within the same topologically associating domain (TAD), we reasoned that true CRE-gene links would be enriched for sharing the same TAD. We considered the TAD annotations derived from Hi-C profiling of neural progenitor cells isolated from the developing mouse neocortex (*99*). After intersecting TAD coordinates with CREs and transcription start sites, we determined the fraction of CRE-gene links in which the CRE and gene’s TSS are co-located within the same TAD. This was done for two sets: (i) CRE-gene pairs present in the GRNs and (ii) all CREs within 250 kb of each gene that were not incorporated in the GRNs.

Finally, we evaluated the quality of TF-gene links in the GRN using Gene Regulatory Network Performance Analysis (GRaNPA) (https://git.embl.de/grp-zaugg/GRaNPA/-/tree/865c0013) (*100*). GRaNPA uses gene expression samples not included in the GRN inference step (i.e., holdout samples) to assess how well differential expression of target genes can be predicted based on the differential expression of TFs. To this end, GRaNPA develops a random forest regression model grounded on the TF-gene adjacency matrix. This model predicts the fold changes of genes that display differential expression between samples that were not part of the GRN inference. We used mouse cerebellum samples previously profiled with snRNA-seq but not included in this study at E14 (SN088 and SN101) and P4 (SN038 and SN044) stages (*8*). We aggregated gene expression counts across all cells in each sample and determined differential expression between developmental stages (E14 vs P4) using DESeq2 (*101*). Filtering criteria of an absolute log2 fold change ≥ 1.0 and an adjusted P-value < 0.05 resulted in the identification of 1,035 differentially expressed genes. After excluding genes absent in the mouse GRN, 743 genes remained. We then provided these differentially expressed genes and a TF-gene adjacency matrix based on the mouse GRN as inputs to GRaNPA_main_function in a 10-fold cross validation setting. We compared the distributions of R^2^ values between actual and predicted log_2_ fold expression changes to those obtained using a randomized GRN or permuted DE gene lists. Additionally, we visualized the relationship between actual and predicted log2 fold changes for one cross-validation set using plot_GRaNPA_scatter.

### Comparisons of GRNs across species

We compared the GRNs between human, marmoset and mouse with the focus on activating TF-gene interactions GRNs. First, we converted the genes and CREs in the marmoset and mouse GRNs to their 1:1 orthologous counterparts in human. We then intersected TFs controlling GRNs across species and identified 114 TFs shared among human, marmoset, and mouse. To investigate the association between conservation and network centrality, we computed the PageRank centrality of the TFs in the human TF-gene network using the networkx package (v2.8.6) and compared centrality estimates across conservation levels. For subsequent analyses, we focused on GRNs composed of these conserved TFs (i.e., recalled in the GRNs of all species) to facilitate comparison between species.

The conservation of different regulatory layers in the GRNs (i.e., TF-CRE, TF-gene, and CRE-gene) was assessed by calculating the ratio of connections conserved between species to the total number of connections in the human GRN at each layer. To examine how marker genes conserved across species are regulated in the GRNs, we used the list of marker genes identified in our previous study (*8*). Marker genes conserved between human, mouse, and opossum, or between human and mouse, were classified as conserved markers, while all other marker genes were classified as unique markers. We further stratified the target genes based on whether they are TFs, using a previously published human TF list (*102*).

To compare the cell type-specific activity of the GRNs controlled by the conserved TF regulators across species, we calculated regulon specificity scores (RSSs) for each cell type and regulon. The RSSs were then z-normalized for each cell type, and we intersected the top 5 cell types with the largest normalized RSSs for each regulon across human, marmoset, and mouse to identify 128 out of 138 regulons with conserved cell type specificity. Due to the absence of certain cell type labels (isth_N_diff, UBC_diff, and UBC_defined) in our marmoset dataset, we required conservation between only human and mouse for these specific cell types. Additionally, we excluded regulons with gene-based AUC values smaller than 0.5 in any cell types, retaining 110 regulons with conserved cell type-specific activity across species.

Finally, to identify the cell type in which each regulon is most specifically active, we selected the cell type with the largest mean RSS x AUC value across species among the top 5 RSS-ranked cell types in all of the three species. We used PyComplexHeatmap (v1.5.0) to visualize the conserved cell type-specific activity of these regulons. To plot the human TF networks for the GC and Purkinje cell lineages, we extracted TF-gene pairs with a Pearson correlation greater than 0.4. The network plots were generated using CytoScape (v3.10.0).

### Summarizing gene expression and chromatin accessibility by cell type and developmental stage

For each species in our dataset, we aggregated gene expression and chromatin accessibility profiles across cells from the same sample (biological or technical replicate), cell type (annotation level 2) and developmental stage. We introduced the following two modifications to our cell type annotation levels for this analysis, aiming to achieve the maximum granularity while in parallel retaining adequate cell numbers to reliably estimate molecular profiles: 1) Progenitor cells were further separated into “gliogenic”, “bipotent” and “early” progenitors, the latter containing all remaining progenitor groups that are most prevalent in embryonic development. 2) Ventricular zone and NTZ neuroblast states (originally designated as 1,2,3 based on their progression along their developmental trajectories) were collapsed into a single group.

Initially focusing on human and mouse, the best captured species in our dataset, we required at least 50 cells in at least two samples for the same cell type and developmental stage in both modalities (RNA and ATAC). We then aggregated the counts of each gene/CRE across all cells in that group (pseudobulks by sample). For human, mouse and opossum, for which RNA and ATAC were profiled separately, the contribution of each cell profiled with ATAC to the pseudobulk was weighted by the prediction score of its cell type label, allowing cells with more confident annotation to contribute more to the bulk profiles. This step was not necessary for bonobo, macaque and marmoset, for which both modalities were captured from the same cell.

We then scaled by sequencing depth (CPM) and additionally applied quantile normalization within each species and modality as implemented in the R package preprocessCore (1.54). For the gene expression matrix, we considered full-length transcripts, as we observed that these show higher correlation between species compared to counting reads in exons only. Prior to scaling, we only retained protein-coding genes within each species, to avoid diluting out CPM levels for species with more extensive non-coding transcript annotations. To estimate gene expression and chromatin accessibility profiles by cell type and stage, we estimated the mean quantile-normalized CPM value of each feature across replicates.

### Non-negative matrix factorisation of CRE accessibility in human and mouse

We used the pseudobulks described above to construct the feature space for the non-negative matrix factorisation (NMF) analysis by selecting highly accessible and highly variable CREs separately in human and mouse. This analysis was limited to 45 groups (cell type and developmental stage) that had at least 50 cells in at least two replicates in both modalities (RNA and ATAC) in both human and mouse. To match human and mouse stages we relied on the developmental stage correspondences described above (e.g., a 11 wpc human sample was considered corresponding to a E15.5 mouse sample). To identify highly accessible reproducible CREs in each species, we filtered for at least 5 CPM in at least two replicates within the same group (cell type and developmental stage). Amongst these CREs, we identified the most highly variable by estimating a variance/mean ratio across all cell types and developmental stages. For each species, we retained the 100,000 CREs with the highest variance/mean ratio. We next standardized the accessibility of each CRE within each species, by scaling to its maximum value across all groups (i.e., after this step the accessibility of each CRE ranged from 0 to 1).

We used non-negative matrix factorisation (NMF) to summarize the standardized accessibility of human and mouse CREs along the 45 corresponding samples into a set of factors. In NMF, the original matrix [CREs x samples] is approximated by the multiplication of two new matrices that correspond to the loadings of a predetermined number of factors on CREs [CREs x factors] and samples [factors x samples] respectively. Thus, samples with similar accessibility patterns are grouped together into the same factor, along with CREs with the highest accessibility in these samples.

To determine the optimal number of factors for our dataset, we considered multiple values in a range between 2 and 30. After each factorization step, we estimated the reconstruction error between the original [CRE x samples] matrix and the one inferred by the multiplication of the two factor-based matrices. Additionally we evaluated the degree of mixing between species by calculating the Euclidean distance between human and mouse CREs in the [CREs x factors] matrix. Naturally, the reconstruction error decreases with the addition of more factors, albeit with a smaller rate after 18 factors (fig. S7B). Similarly adding more factors, especially above 25, leads to a greater separation of the two species in the factor space (fig. S7B). Based on these two metrics, and by considering the biological relevance of the identified associations between samples and factors, we determined the optimal number of factors to be 18.

To associate samples (cell types x developmental stages) with factors, we considered the [samples x factors] matrix (sample loadings). For each factor, we considered samples with a loading at least equal to 40% of the maximum loading as associated with that factor. In practice, this led to 1-5 samples assigned to each factor, and to each sample being associated with up to 2 factors. Reassuringly, samples from the same cell type or developmental stage were grouped together in the same factors.

Similarly, we assigned human and mouse highly variable CREs to each factor based on the [CREs x factors] matrix (CRE loadings). For this we used elbow plots to determine the optimal cutoff, adapting a procedure introduced by Gerrard et al. 2020 (*103*). For each factor, we plotted the CRE loadings in ascending order and determined the elbow point as the CRE that minimized the Manhattan distance from the intercept of the maximum loading with the x-axis. CREs with loadings greater or equal to the elbow point were assigned to that factor. In practice, this led to 24,569-38,529 of the 200,000 human and mouse highly variable CREs assigned to each factor and to 90% of these CREs being assigned to up to 5 factors (30% to a single factor).

We used pycisTarget (v1.0.1) (*43*) to identify the TF motifs enriched in sets of human and mouse highly variable CREs associated with each NMF factor. We used the custom cisTarget databases constructed on our peak sets, as detailed in the section *Gene regulatory network (GRN) inference*. To enable quantitative comparisons of TF motif enrichments between human and mouse CREs, we first used a lenient normalized enrichment score (NES) cutoff of 0.1. We then identified significantly enriched motifs as those that reached a NES of at least 3 in at least one species. For these TF motifs, we computed the Pearson correlation in NES scores across NMF factors in human and mouse, as a metric of similarity in the TF code of corresponding cell types between species.

To project marmoset CREs on the previously identified NMFs for human and mouse, we first identified highly accessible CREs as those reaching at least 5 CPM in at least two replicates within the same group (cell type and developmental stage). We then subsetted the previously estimated sample loadings [samples x factors] for the 34 groups (cell type and developmental stage) that were shared between human, marmoset and mouse and used scipy’s (v1.7.1) implementation of non-negative least squares (scipy.optimisation.nnls) to estimate the [CREs x factors] matrix. This allowed us to estimate the loadings of marmoset CREs on the NMFs learned from the human and mouse CREs. We applied the same procedure to project all highly accessible human and mouse CREs irrespective of their variability, which allowed us to also identify CREs with high loadings in many/all NMFs.

### Sequence-based models of CRE accessibility

We trained a multiclass multilabel classifier to predict CRE assignment to NMF factors based on their DNA sequence, akin to previous studies (*29, 32*). For each species and for each factor, we considered the 3000 highly variable CREs with the highest loadings as assigned to that factor. To allow the model to generalize beyond cell type-specific CREs, we additionally incorporated two more sets of CREs. First, we selected 3000 random intergenic regions that did not overlap CREs from our study, Ensembl annotated exons, genome assembly gaps or CREs in other tissues (ENCODE v3) (*104*). These regions were considered putatively inactive and assigned zero membership to all NMFs. Second, we selected 3000 lowly variable CREs assigned in all 18 NMFs (enriched for promoters). We used 80% of these CREs for training models, 10% for validation and 10% for testing. Since the model considers individual CREs, the training-validation-test splits included CREs from all chromosomes. To increase the number of sequences used for training and to allow the model to focus on relevant sequence features, augmentation was performed by extending the training sequences by 100 bp towards each side, then using a sliding window of 500 bp with a stride of 50 bp to generate partially overlapping sequences with the same label. CRE sequences were extracted using bedtools getfasta and converted to a one-hot encoding format.

The models use a hybrid architecture of convolutional and recurrent neural networks, akin to that used in previous studies (*29, 32*). Briefly, one-hot encoded sequences are used as input for a convolutional layer with 512 kernels of size 24, followed by a max-pooling layer with size and stride of 16. Of the 512 kernels, 285 were initialized with motifs from the JASPAR 2020 database (*105*). The output of the convolutional network is fed into a time-distributed dense layer together with a bidirectional long short-term memory (LSTM) layer with 256 neurons. Finally, the output of the LSTM layer is passed on to a flattened and then to a dense layer, which in turn uses a sigmoid activation function to estimate prediction probabilities for each of the 18 possible classes (NMFs). The remaining activation functions were constructed using Leaky ReLU. To prevent overfitting, dropout layers are introduced after the max-pooling, LSTM and dense layers with dropout rates of 0.5, 0.2 and 0.5, respectively. The model architecture was implemented in TensorFlow (v2.9.1). Training was performed on NVIDIA A40 GPUs using the Adam optimizer with a batch size of 128 and a learning rate of 0.001 for a minimum of 10 and a maximum of 100 epochs, allowing for early stopping. The best epoch was selected as the one that minimized the binary crossentropy loss in the validation set. Model performance was evaluated using the auROC and auPR metrics, estimated based on the testing dataset using the average_precision_score and roc_auc_score functions from the scikit-learn package. To compare model performance in single-species versus mixed-species models, we used the same training-validation-test split. To control for the impact of the training set size on model performance, we first generated a human-mouse mixed dataset that contained 50% of CREs from each species. After establishing the higher performance of multispecies models for multispecies predictions, we trained DeepCeREvo by combining the full training set from both human and mouse.

### Interpretation of sequence-based models

We used three complementary approaches to identify the sequence features driving our model’s predictions. For individual CREs from the test set with high prediction scores in the factor of interest, we estimated the contribution of each nucleotide to the model’s prediction using the package DeepExplainer SHAP (v0.37.0) (*106*). The explainer was initialized by shuffling the CRE 100 times while preserving dinucleotide frequencies, akin to previous approaches (*107*)). Hypothetical contribution scores were multiplied with the one-hot encoded matrix and visualized based on the viz_sequence function of the package DeepLift (*107*).

We additionally employed *in silico* saturation mutagenesis to mutate each position to every other possible nucleotide and measure the effect on the model’s prediction. Here, sequence features important for the model’s predictions were identified as nucleotide positions that lead to a large decrease in the model’s predictions when mutated, akin to previous approaches (*29*).

Finally, to comprehensively investigate the sequence code of each NMF factor, we used TF-MoDIsco (v0.5.6.5)(*51*). For each species and NMF, we identified the 2500 CREs with the highest prediction scores amongst those that belonged to this factor. We additionally required CREs to be assigned to a maximum of 5 NMFs, to avoid prioritizing broadly accessible regions that tend to receive high prediction scores across all factors. This led to a total set of 36,934 human and 36,760 mouse CREs (the theoretical maximum of 2500 CREs across 18 factors was 45,000). We used DeepExplainer SHAP as described above to estimate nucleotide importance scores per CRE across all factors. For each species and factor, we provided the SHAP values as input to TF-MoDIsco with the following parameters: n_sample_null=5000, trim_to_window_size=15, sliding_window_size=15, flank_size=5, initial_flank_to_add=5, final_flank_to_add=5, final_min_cluster_size=100, target_seqlet_fdr=0.05, kmer_len=8, num_gaps=3 and num_mismatches=2.

Briefly, TF-MoDIsco uses the SHAP scores to identify regions (seqlets) within each CRE that have high importance in the model’s prediction towards a certain class/NMF. Subsequently, seqlets are clustered based on their activity across tasks. In this case, since we run the algorithm separately for each factor, there are only two activity patterns (metaclusters), +1 for positive importance and −1 for negative importance. Within each metacluster, seqlets are then clustered based on their sequence similarity, with the goal of identifying motifs (patterns) with non-redundant sequence and non-redundant activity. The precise pattern boundaries are further refined by trimming, expanding and centering based on all seqlets in this group. The precise steps performed by TF-MoDIsco are detailed in Shrikumar et al. 2018 (*51*).

Since we run TF-MoDIsco separately for each species and factor (following a recommendation by the original developers and to make the problem computationally feasible with available resources), we had to further integrate the patterns identified in each run. For example, the same TF motif could be identified as human_NMF_2_metacluster0_pattern2 and as mouse_NMF_5_metacluster1_pattern5. To identify pattern correspondences across independent TF-MoDIsco runs, we extracted the trimmed motifs, converted them into the MEME format and used Tomtom (v5.5.1) (*108*) to quantify sequence similarity between patterns identified across species and factors, specifying the following parameters: dist=kullback, motif-pseudo=0.1, min-overlap=1. We clustered patterns based on the Pearson’s correlation between the −log_10_ transform of the reported q-value to identify patterns with similar sequence. We cut the dendrogram at a height of 0.7, considering patterns falling into the same cluster as corresponding (though we note that our results are consistent across a range of height thresholds). We additionally used Tomtom with threshold=0.3 to match patterns against known TF motifs from the JASPAR 2022 database (*96*).

### Regulatory grammar conservation across mammals

To assess similarity in CRE sequence codes between species and NMFs, we extracted the number of seqlets detected per pattern, as stored in the modisco output: modisco_hdf5[‘metacluster_idx_to_submetacluster_results’][metacluster_name][‘seqlets_to_patterns_result ‘][‘patterns’][pattern_name]. We only considered metaclusters with positive contributions for each class, as negative contributions mainly correspond to activators of other classes rather than repressors due to our model being a multiclass and multilabel classifier. For patterns matching the same motif cluster, we used the maximum number of seqlets instead of aggregating across patterns, reasoning that most patterns with such high sequence similarity would predominantly contain overlapping rather than distinct seqlets. This procedure resulted in a [motif_cluster x class] count matrix, where each class corresponds to a distinct combination of species and NMF. We note that the counts in this matrix do not merely correspond to occurrence of a motif in a set of sequences but to these motifs being considered important for prediction in this class by DeepCeREvo, after taking into account local context such as flanking sequences and TF combinatorics. Reasoning that such a matrix represents a reasonable approximation of CRE sequence code similarity, we next performed hierarchical clustering across NMF classes and motifs using Pearson’s correlation as a metric and “average” as a clustering method. To assess the robustness of our clustering of different classes, we used the R package pvclust (v2.2-0) to perform bootstrapping with 10,000 iterations. All motif clusters were used for clustering, but for visualisation we filtered the matrix for motifs with at least 1000 seqlets across all classes and capped at the 99th quantile.

To examine the patterns assigned to the same motif cluster across different classes for more subtle differences in their sequences, we used the function view_motifs() from the R package universalmotif (v1.10.2) to align (and when needed, reverse complement) patterns. We also used the function merge_motifs() from the same package to extract the consensus motif of each motif cluster for visualization purposes.

Individual seqlet occurrences in human and mouse CREs were detected based on TF MoDISco’s density adapted hit scoring function (densityadapted_hitscoring.MakeHitScorer() followed by hit_scorer.set_coordproducer() with max_seqlets_total=np.inf (i.e., infinite). We then scored each peak for the presence of at least one seqlet belonging to a motif cluster of interest. This binarization step allowed us to account for overlaps between seqlet instances and similar patterns assigned to the same motif cluster.

### Inference of CRE evolutionary history from model’s predictions

We inferred the evolutionary history of human CREs by applying DeepCeREvo to their orthologous loci in over 200 mammalian species. First, we liftovered the sequences of 554,237 human CREs to the genomes of 240 mammalian species using halLiftover (*109*) on the Zoonomia version 2 alignment (*20*), followed by halLiftover Post-processing for the Evolution of Regulatory elements (HALPER) (*110*). HALPER builds contiguous orthologues from halLiftover outputs by extending liftovered 1 bp focal positions to both sides until all liftovered fragments on the same contig are covered or until the pre-defined maximum length is reached. To ensure that a focal position was both important for the accessibility of a human CRE and conserved across as many species as possible, we identified the position within 50 bp from the center of each human CRE that was mapped to the largest number of species using halAlignmentDepth with default parameters (*109*). We then extended or trimmed the output regions in each species, maintaining the same distance ratio from the focal position, to set their length to 500 bp. On average, 392,167 (70.8%) human CRE sequences were liftovered to each species, and the number of liftovered loci in each species was well correlated with their evolutionary distance from human (fig. S11C). We removed 13 species (5.3%) from subsequent analyses which had fewer liftovered regions compared to other species with similar evolutionary distance from human, by setting a threshold based on the 95% confidence interval of the linear regression between the number of liftovered regions and the evolutionary distance. We manually confirmed that these excluded species had lower assembly quality than the remaining species. Finally, for each species we extracted the sequences of regions orthologous to human CREs and applied DeepCeREvo to obtain prediction scores across NMFs.

Based on the DeepCeREvo’s prediction scores in orthologous loci across mammals, we inferred the evolutionary histories of human CREs. We grouped 227 mammalian species (228 genomes, including 2 *Canis lupus* genomes) into 8 non-overlapping categories, named by representative clade in the category: Human (1 species); Non-human hominid (4 species), Old World monkeys (17 species); New World monkeys (10 species); Prosimians (11 species); Glires (59 species); Laurasiatheria (112 species); and Afrotheria/Xenarthra (14 species). We then combined these categories to make hierarchical clades containing human: Human (1 species); Great apes (5 species); Catarrhini (22 species); Simiiformes (32 species); Primates (43 species); Euarchontoglires (102 species); Boreoeutheria (213 species); and Eutheria (227 species).

For each NMF, we identified human CREs whose prediction score arrays across 227 mammalian species showed signals specific to any of the various clades defined above. For missing values (i.e., where no orthologous regions to human CREs were found), we imputed zero prediction scores. To this end, we applied three criteria based on: (1) the fold-change of prediction scores between species within the clade of interest and those outside of it, (2) empirical *P*-values using the fold-change of prediction scores as a test statistic, and (3) a minimum prediction score values in species within the clade of interest.

1. We set the threshold of the fold-change of prediction scores between ingroup and outgroup species as 1.5. To alleviate species representation bias in ingroups and outgroups, we computed median prediction scores for each of the 8 non-overlapping species categories and required that any combination of categories in the ingroup and outgroup meets the fold-change threshold.
2. To evaluate the probability of observing clade-specific signals by random chance, we computed empirical *P*-values. Traditional permutation—shuffling values across samples (in this case, species)—is not optimal because species are not independent, but rather related to each other based on evolutionary distances. Therefore, we employed the recently proposed approach of phylogenetic ‘permulations’ (*53*). In this approach, the Brownian motion model of continuous trait evolution is applied to the phylogenetic tree to assign simulated phenotype values. The observed values (in this case, prediction scores) are then assigned to the simulated values based on their rankings. This ensures that species close to each other have similar phenotypic traits (here, predictions), reflecting evolutionary distances, while maintaining the distribution of the original observed values. For the test statistic, we used the fold-change of mean prediction scores between ingroup and outgroup species. The mean scores were calculated by averaging the mean prediction scores of non-overlapping species categories belonging to ingroups and outgroups. We then conducted 10,000 permulations for each array of prediction scores across 227 mammals and computed the *P*-value as the fraction of simulations in which the fold-change statistic exceeded the observed value.
3. To determine the prediction score threshold for ingroup species, we identified the thresholds that optimized the performance of predicting accessible regions in human for each NMF factor. We built a binary classifier to predict accessible regions using prediction scores from the same NMF factor and calculated precision and recall for different thresholds, ranging from 0.00 to 1.00 in increments of 0.01. The threshold was then set as the score which maximized the F1 score, the harmonic mean of precision and recall, for each of the 18 NMF factors (table S11). We required that the median prediction scores of each non-overlapping category in ingroups exceeds this threshold for the corresponding NMF factor.

### Evaluation of CRE evolutionary history predictions

We evaluated our evolutionary classification by examining the accessibility of orthologous loci to human CREs in bonobo, macaque, marmoset, and mouse. We assessed all NMFs in marmoset and mouse, and additionally considered bonobo and macaque for mature granule cells (NMF 14), which were robustly detected in all species. When considering all NMFs, we computed the fraction of human CREs assigned to each NMF that had an orthologous CRE assigned to the same NMF in marmoset or mouse. Since we mapped marmoset data to the calJac4 assembly whereas Zoonomia used an earlier assembly (ASM275486v1), we liftovered marmoset regions from ASM275486v1 to calJac4 using UCSC liftOver with a custom chain file, which we generated using nextflow-LiftOver (nf-LO) with the --blat --distance near options (*111*). Liftovered regions that were more than 25 bp shorter or longer than the original regions were discarded. As a result, 494,914 out of 500,582 (98.9%) marmoset loci orthologous to human CREs were retained for the analysis.

In the analysis focusing on mature granule cells (NMF 14), we examined the accessibility of orthologous loci to human CREs assigned to this NMF. To this end, we used the addPeakSet() and addPeakMatrix() functions in ArchR to compute the number of fragments detected in loci liftovered from human CRE sequences in each species. For marmoset, we used the coordinates liftovered to calJac4 from ASM275486v1, as described above. We then aggregated the fragment counts of each liftovered locus for cells annotated as GC_defined in each sample to generate a pseudobulk count matrix, which was scaled for sequencing depth to obtain CPM profiles. Pseudobulk fragment counts were averaged across samples within each species, and the resulting CPM values were compared based on the predicted evolutionary clades of the human CREs.

### Selection and cloning of CREs for luciferase reporter assays

For testing CREs’enhancer activity in mouse primary granule cells we focused on CREs accessible in granule cell progenitors or differentiating granule cells (NMF 18, 2, 13 and 8), because these are very abundant in early postnatal mouse development and robust protocols for their culturing *ex vivo* were available. We constructed three sets of CREs to be tested: conserved CREs, divergent CREs and shuffled sequences.

For the conserved CREs, we selected CREs classified as eutherian-shared based on DeepCeREvo’s predictions. We additionally prioritized CREs that were within 250 kilobases (kb) of a gene with conserved granule cell-specific expression in human, mouse and opossum, as identified previously (*8*). We further prioritized CREs for conserved and specific accessibility in the granule cell lineage in our dataset. Finally, we manually investigated DeepCeREvo’s SHAP attribution and *in silico* mutagenesis profiles of these CREs to ensure that they contained motifs recognised by TFs active in the granule cell lineage (primarily E-boxes and NFI motifs). We included orthologous sequences from human and mouse.

Divergent CREs were selected based on DeepCeREvo’s classification into a clade with an evolutionary age of 40 million years or younger (simians, catarrhines, hominoids or human-specific), and with accessibility profiles in human, marmoset and mouse that were consistent with the model’s predictions. We additionally prioritized CREs within 250 kb of a gene with divergent expression between human and mouse in a relevant NMF (and when possible matching the direction and timing of the CRE gain). Finally, we manually investigated DeepCeREvo’s SHAP attribution and *in silico* mutagenesis profiles of these CREs across species and prioritized those that gained motifs recognised by TFs active in the granule cell lineage. For selected divergent CREs we included sequences from up to 5 species in addition to the human sequence. The selection of species was in some cases influenced by the provider’s (IDT) restrictions on sequence features for DNA fragment synthesis

Shuffled sequences were generated by randomly shuffling the nucleotides in human sequences of eutherian-shared CREs, preserving GC content and dinucleotide frequencies. We repeated the shuffling 1,000 times and used DeepCeREvo to generate predictions across all 18 NMFs. We then selected sequences with the lowest maximum prediction across NMFs.

As a backbone for enhancer reporter constructs we used pNL1.2[NlucP] vector (Promega) that encodes for NanoLuc-PEST (NlucP) reporter protein. All cloning steps were performed using In-Fusion Snap Assembly Master Mix (Takara) in miniaturized reaction volumes (1-2 **μ**l). First, we designed adapter sequences that would allow insertion of the CREs in forward and reverse orientations (table S13). Overlapping oligos (Sigma-Aldrich) for these sequences were subjected to 3 cycles of PCR with KAPA HiFi HotStart ReadyMix (Roche), the obtained fragments were cleaned up using 1.8x SPRIselect beads (Beckman Coulter), and inserted to the pNL1.2[NlucP] vector linearized with KpnI and EcoRV to produce pNL1.2_adaptersF_NlucP and pNL1.2_adaptersR_NlucP vectors. Next, SV40 promoter sequence was amplified from pHRdSV40-scFv-GCN4-sfGFP-VP64-GB1-NLS vector (*112*), which was a gift from Ron

Vale (Addgene plasmid #60904), using KAPA HiFi HotStart ReadyMix (Roche) and primers with overhanging homology arms (table S13). The fragment was inserted into pNL1.2_adaptersF_NlucP and pNL1.2_adaptersR_NlucP vectors, linearized with HindIII and NcoI. Finally, the obtained pNL1.2_adaptersF_SV40_NlucP and pNL1.2_adaptersR_SV40_NlucP vectors were linearized with KpnI and SacI, and used for the insertion of CRE sequences in front of the SV40 promoter in either forward or reverse orientation. CRE sequences with suitable homology arms (table S13) were synthesized as eBlocks at IDT. Constructs were purified using Monarch Plasmid Miniprep Kit (NEB), and verified by Sanger sequencing (Azenta Life Sciences). Firefly luciferase vector pGL4.15[luc2P/EF1α/Hygro], a gift from Priit Pruunsild, was obtained by inserting the BamHI and HindIII fragment with EF1α promoter sequence (hg38 chr6:73,520,038-73,521,250) from pGL4.83[hRlucP/EF1α/Puro] (*113*) into pGL4.15[luc2P/Hygro] (Promega) between BglII and HindIII. Purity of the plasmids was evaluated by Nanodrop (Thermo Fisher Scientific) and concentrations were determined by Qubit (Thermo Fisher Scientific) measurements.

### Transfections and luciferase reporter assays

Primary cultures of mouse granule cells grown on 96-well plates were transfected at DIV2 using FuGENE HD transfection reagent (Promega) at a DNA:reagent ratio of 1:4. The medium was replaced with optiMEM for the duration of transfection (4-5 hours). For each well, we co-transfected 95 ng of different pNL1.2[NlucP]-based vectors and 30 ng of pGL4.15[luc2P/EF1α/Hygro] that expresses firefly luciferase used as a normalizer. All transfections were performed in triplicates or quadruplicates. 28-30 hours after transfection the cells were lysed in Passive Lysis Buffer (Promega; 50 **μ**l per well) and subjected to a freeze-thaw cycle. Nano-Glo Dual-Luciferase Reporter Assay System (Promega) was used for the detection of firefly and NanoLuc luciferases, following the manufacturer’s instructions. Luminescence signals were measured on Spark Cyto plate reader (Tecan) using 3 seconds integration times.

For the estimation of enhancer activities, only wells with firefly luciferase signals at least 3-fold above background were included. Signals from untransfected wells were subtracted, NanoLuc luciferase signal values were normalized to firefly luciferase signals, and the ratios were log_2_-transformed. To normalize the spread across independent experiments, we estimated experimental standard deviations, based on data points present in all independent experiments. We divided values by experimental standard deviations and multiplied by the global standard deviation to adjust the spread to a common scale. For statistical inference, we applied linear mixed models using the R packages lme4 (v1.1-36) (*114*), lmerTest (v3.1-3)(*115*), and pbkrtest (v.0.5-0.1) (*116*). Interaction model was fitted twice using Restricted Maximum Likelihood (REML) to contrast each element with the no-enhancer control with either forward or reverse orientation adapter sequences. The model included the element, orientation, and their interaction as fixed variables (CRE * orientation), and independent experiments (n = 7) as a random variable (1 | experiment; experiment variance = 0.5531, residual variance 0.1149). The residuals were normally distributed (*P* = 0.3181, Shapiro-Wilk test), supporting the model’s fit. The effect of orientation was significant (*P* < 9.32 × 10^−5^, no-enhancer control log_2_(fold change) = 0.37), reflecting a mild impact of construct orientation. Compared to the additive model (CRE + orientation), including the interaction term significantly improved model fit (*P* < 10^−15^, Likelihood Ratio Test based on ML models), indicating that the effect of orientation varies per CRE (table S13). The Kenward-Roger approximation was used for computing the degrees of freedom and t-tests. *P*-values from the model, including both main CRE effects (forward and reverse) and interaction effects (forward versus reverse), were corrected for multiple comparisons using the Benjamini-Hochberg method. For clarity, in Fig. 5, F and G, and fig. S13G, *P*-values are reported only for the bars with log_2_(fold change) ≥ 0.5. All values are listed in table S13.

### Sequence conservation of human CREs across evolutionary clades

We first estimated the age of human CRE loci based on the species with orthologous loci, as identified in the section *Inference of CRE evolutionary history from model predictions*, and compared it with the evolutionary clades defined by prediction scores. For each human CRE, we calculated the fraction of species with orthologous regions within each of the eight non-overlapping species categories. The minimum evolutionary age was defined as the smallest clade that includes all species categories in which at least 20% of species have a 1:1 orthologous region corresponding to the human CRE.

We then assessed the sequence conservation levels of human CREs assigned to different evolutionary clades using genome-wide phyloP and phastCons scores across mammals and primates, respectively (*20, 21, 117, 118*). Conservation scores were extracted using pyBigWig (v0.3.18), and for each CRE, the highest sum of conservation scores within any 100 bp window was used as the conservation level. We then compared these conservation levels between human CREs from different evolutionary clades.

To evaluate whether the model’s predictions perform better than sequence conservation in predicting the conservation of CREs within an evolutionary clade, we examined the accessibility of human CREs in marmoset (primate-conserved) and mouse (eutherian-conserved). To assess the performance of our evolutionary classification rather than DeepCeREvo’s predictions in a single species, we averaged prediction scores of human CREs across primates and mammals. Similarly, we used phastCons scores across primates and phyloP scores across mammals, as a metric of the overall evolutionary conservation of human CREs within these clades. We then developed binary classifiers based on the mean prediction and sequence conservation scores to predict whether human CREs would have an orthologous CRE assigned to the same NMF in marmoset or mouse. We computed true and false positive rates at different thresholds and compared the area under the receiver-operating curve (AUROC) between the prediction and conservation score classifiers.

### Overlap with human accelerated regions

We compared human CREs with different evolutionary histories to previously identified non-coding regions that have undergone human-specific accelerated substitutions or deletions. We downloaded bed files for HARs (*57*), hCONDELs (*25*), and HAQERs (*24*). We then intersected these genomic regions with human CREs using bedtools intersect (v2.30) and compared the fraction of overlap with these elements across CREs with different evolutionary histories.

### Estimation of selection parameters from sets of human genomic regions

To infer the direction and magnitude of selective pressure acting on human CREs assigned to different evolutionary clades, we applied a Bayesian method to estimate selective pressure on a set of regions using allele frequency data from human populations, as described in previous studies (*24, 58*). We downloaded allele frequency datasets of five African populations (Gambian, Mende, Esan, Yoruba and Luhye in Webuye, Kenya) and sets of variants (random and missense) (*24*).

Given that DeepCeREvo indicated only a subset of nucleotides within 500-bp CRE regions are important for its prediction, we extracted the 10 nucleotides with the highest attribution scores for each CRE inferred to be human-specific or eutherian-conserved. As a background, we selected 10 nucleotides with the lowest absolute attribution scores. We obtained up to 1,000 polymorphic sites per nucleotide set in each population and computed selection parameters based on the allele frequency spectrum of each set of polymorphic sites using the intervalOverlap and selectionMcmc functions in gonomics (v1.0.0) with default parameters (*24, 119*).

### Identification of radical gene expression changes between human and mouse

We extended our previous approach to detect changes in the expression of 1:1 orthologous protein-coding genes between species (*8*) to be able to identify cell type and state-specific gene expression changes in a common framework to our analyses of CRE accessibility described above. To this end, we made use of the aggregated gene expression profiles across cell groups, developmental stages and replicates described in the section *“Summarizing gene expression and chromatin accessibility by cell type and developmental stage”*. We identified differentially expressed genes between human and mouse for each factor in our CRE-focused NMF analysis by considering the samples that were assigned to that NMF. NMFs 2 and 13 were considered jointly due to their high similarity in gene expression profiles, whereas NMFs 1 and 11 were excluded as they represent heterogeneous groups of neuroblasts, and any detected gene expression differences could be attributed to differences in the cellular composition of these groups.

Akin to our previous work (*8*), we strived to account for the challenges associated with detecting differentially expressed genes between species with 3’ data by jointly considering changes in absolute (CPM scaled and quantile normalized) and relative (fraction of maximum within species) expression. Considering the relative expression of a gene within a species is important as absolute expression levels can arise from differences in gene annotations (loss of reads if UTR is unknown) as well as in gene/transcript length (longer genes are more likely to contain polyA stretches and thus get captured during reverse transcription) (*120*). These factors are expected to affect all cell types in the same way (with the exception of alternative splicing, which we cannot account for with our 3’ datasets). Thus, additionally requiring differentially expressed genes to show differences in their relative expression can minimize the risk of false positives in such an analysis. On the other hand, relying exclusively on relative expression is sensitive to overestimating the degree of dissimilarity for lowly expressed transcripts, which is why we opted for considering both metrics (fig. S16A).

An additional consideration is whether differences in expression are reproducible across biological replicates and different 10x technologies (some of our samples were profiled with 10x 3’ v2 and others with v3). Since our goal in this analysis was to identify the most striking gene expression differences between human and mouse, which we could then examine in the context of changes in their chromatin accessibility environment, we opted for a conservative approach that favors false negatives over false positives. To achieve this goal, we required at least two replicates in species A to show higher expression than the highest replicate in species B to determine a gene as more highly expressed in species A (fig. S16A).

In light of the considerations described above, we implemented the following procedure to identify differentially expressed genes between human and mouse. First, we identified genes that are robustly expressed in the cerebellum in both species, requiring them to reach at least 100 (quantile normalized) CPM in at least two replicates from the same sample (cell group and developmental stage). We only considered genes passing this filter (4,421 1:1 ortholog pairs), for our downstream analysis. For each gene, and for each set of samples associated with an NMF within each species, we estimated the gene’s relative expression as the fraction of its maximum expression within the NMF over its maximum expression across all NMFs. We also estimated the highest and second highest expression across replicates for a given sample (cell group and developmental stage). We then identified genes with higher expression in species A over species B as those showing at least 2.5 fold increase in absolute expression (quantile normalized CPM) between the second highest replicate in species A versus the highest replicate in species B (i.e., at least two replicates in species A show 2.5 fold higher expression than the highest replicate in species B). We additionally required the expression in species A to be above 100 CPM in at least two samples. Finally, we required the relative expression in species A (fraction of the gene’s maximum expression across all groups) to be at least 0.3 and at least 2.5 fold higher than that of species B. These conservative criteria identified a total of 1,883 and 1,325 cases of genes with higher expression in human/mouse respectively out of a total of 66,315 comparisons (4.8% of all comparisons).

To obtain an estimate of the statistical significance of these comparisons, we randomly swapped 50% of our human and mouse samples within each corresponding cell group and developmental stage (to ensure that all comparisons could still be performed). We repeated the shuffling 10 times with different seeds. For each gene, we estimated empirical *P*-values as the fraction of shuffled comparisons showing equal or greater difference in the combination of metrics used to detect differential expression (jointly considering the absolute and relative expression cutoff and fold change). We additionally adjusted our comparisons for multiple testing using the Benjamini-Hochberg procedure as implemented in the R function p.adjust. Unsurprisingly, considering how conservative our approach was, all adjusted *P*-values identified as differentially expressed in our analyses were smaller than 0.01.

To additionally validate our gene expression differences in an independent dataset, we considered publicly available single-cell data for the human and mouse adult cortex profiled with 10x, as well as the full-length protocol SMART-Seq v4 (*11, 59*). For both technologies we used the trimmed-means normalized data as provided in https://portal.brain-map.org/atlases-and-data/rnaseq. For each species and technology, we estimated the maximum expression of a gene across oligodendrocyte, astrocyte and microglia clusters respectively. We then calculated the log_2_(fold change) in human versus mouse expression levels across 1:1 orthologs within each species and technology.

For gene ontology enrichment analysis, we used the R package clusterProfiler (v4.0.5) to query the database org.Hs.eg.db_3.13.0. For each NMF, we estimated enrichments for the genes showing higher expression in human or mouse, using all genes expressed with at least 100 CPM in at least two replicates of at least one sample in the same NMF and species as a background set. Only gene sets with at least three genes were considered and *P*-values were adjusted using the Benjamini-Hochberg procedure. We report enrichments with an adjusted *P*-value < 0.1.

### Polarization and timing of gene expression changes

Having identified genes with significantly higher expression in human or mouse, we sought to use our datasets for marmoset and opossum to determine whether these changes corresponded to expression gains or losses and at which point in evolution they occurred. As our analysis relied on both absolute and relative expression estimates, we did not include bonobo and macaque here due to the limited number of cell groups and developmental stages that would not allow accurate estimation of relative expression levels. Since our analysis only included four species, we decided against using a phylogenetic approach and instead opted for classifying each gene as reliably expressed, non-expressed or uncertain in each species and NMF (fig. S16B). To this end, we started from gene expression estimates for each cell group and stage (averaged across replicates) for human and mouse and performed min-max scaling for both the absolute (CPM) and relative (fraction of maximum expression) levels. This allowed us to construct a “coordinate system” ranging from 0 (species with lower expression) to 1 (species with higher expression) on which we could project the expression levels of marmoset and opossum. If expression in marmoset or opossum was lower/higher than the minimum/maximum of human and mouse, the value was capped to 0/1 respectively.

We collapsed the two estimates into a single score by taking the mean between the absolute and relative expression scores. Genes with scores above 0.8 or below 0.2 were classified as expressed and non-expressed respectively, with everything in between assigned as unresolved. We limited this analysis to genes that were already identified as differentially expressed between human and mouse, and which we could detect with at least 50 CPM in at least one cell group in the third species (we lowered this threshold from 100 CPM used for human and mouse to be able to consider more genes). Since some cell groups and developmental stages were less well captured in marmoset and opossum compared to human and mouse, we additionally excluded genes that had a low expression score in a third species but were also lowly expressed in the corresponding samples of the species in which we originally detected high expression (i.e., they were only highly expressed in human/mouse samples not captured in the third species). We then used this classification scheme to determine the type of evolutionary shift based on the principle of maximum parsimony (fig. S16C). Genes highly expressed in human or mouse and in the outgroup opossum were classified as losses in the non-expressed species, whereas genes reliably non-expressed in opossum were classified as gains. Similarly, for genes we could reliably estimate expression levels in marmoset, we were able to distinguish between higher expression in catarrhines (human high, marmoset low) or primates (human high, marmoset high).

### Chromatin accessibility of genes with expression changes

To assess whether gene expression differences between human and mouse are reflected in local chromatin accessibility profiles, we use ArchR derived “gene scores”. As with our gene expression analysis, we first averaged gene score estimates across cells from the same species, cell group and developmental stage. Then, for each NMF, we estimated the maximum gene score estimate across all samples assigned to that NMF. Finally, we compared these estimates between species for genes assigned to different expression shift classes.

To identify specific CREs associated with the expression changes, for each NMF, we compared genes with higher expression in human compared to mouse to those with high expression in human (at least 100 CPM and 30% of their maximum expression across the dataset). For each human gene, we extracted associated CREs from the SCENIC+ GRN. Since our GRN inference was performed at the level of the entire dataset (i.e., across cell types and stages), we pruned these CREs by requiring them to reach at least 5 CPM in at least one sample assigned to the NMF of interest. This allowed us to identify a set of CREs that putatively regulate the expression of each human gene in each NMF. We next considered whether these human CREs had an orthologous CRE in mouse (conserved vs human-specific). Additionally, for those CREs with a counterpart in mouse, we estimated the log_2_ fold change in their accessibility (quantile normalized CPM, maximum across samples assigned to that NMF) between human and mouse. Finally, for each NMF we combined human CREs that have no mouse CRE ortholog (human-specific CREs) with human CREs reaching at least 5 CPM in that factor that have a mouse ortholog not reaching that cutoff (repurposed CREs) into a single set termed “CREs with human-specific activity”.

## SUPPLEMENTARY FIGURES

**Fig. S1.**
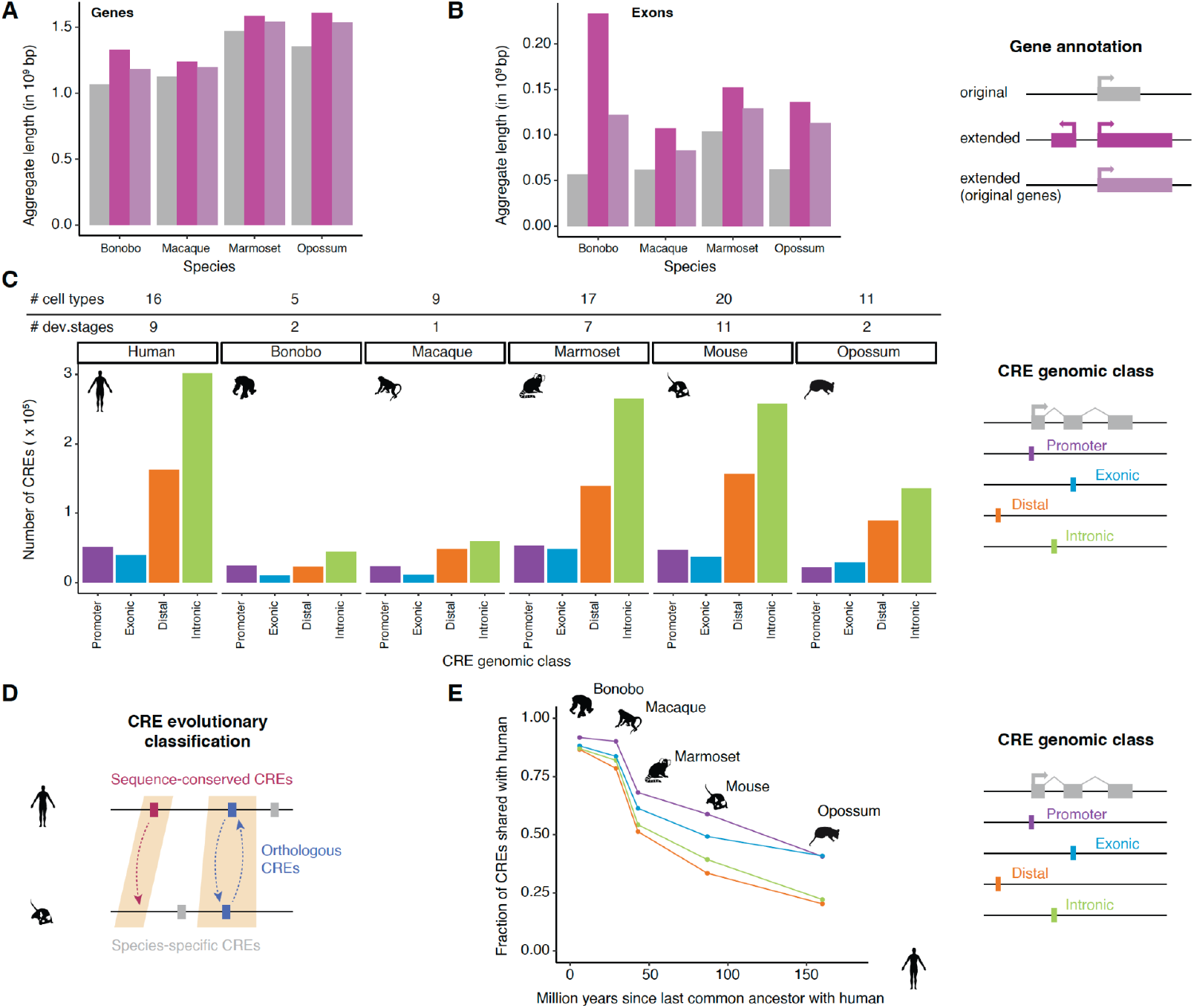
Transcriptomic and regulatory annotations of mammalian genomes. (A, B) Total coverage (in 10^9^ base pairs; bp) of genes (A) and exons (B) in original and extended annotations for bonobo, macaque, marmoset and opossum. (C) Identification and genomic characterisation of CREs in developing cerebellar cell types across mammalian species. Top: number of ATAC-profiled cell types (minimum of 50 cells) and developmental stages per species. (D) Schematic depiction of the identification of orthologous CREs between species based on reciprocal syntenic alignments. (E) Fraction of CREs of non-human species with an orthologous CRE in human. CREs are grouped by genomic class.

**Fig. S2.**
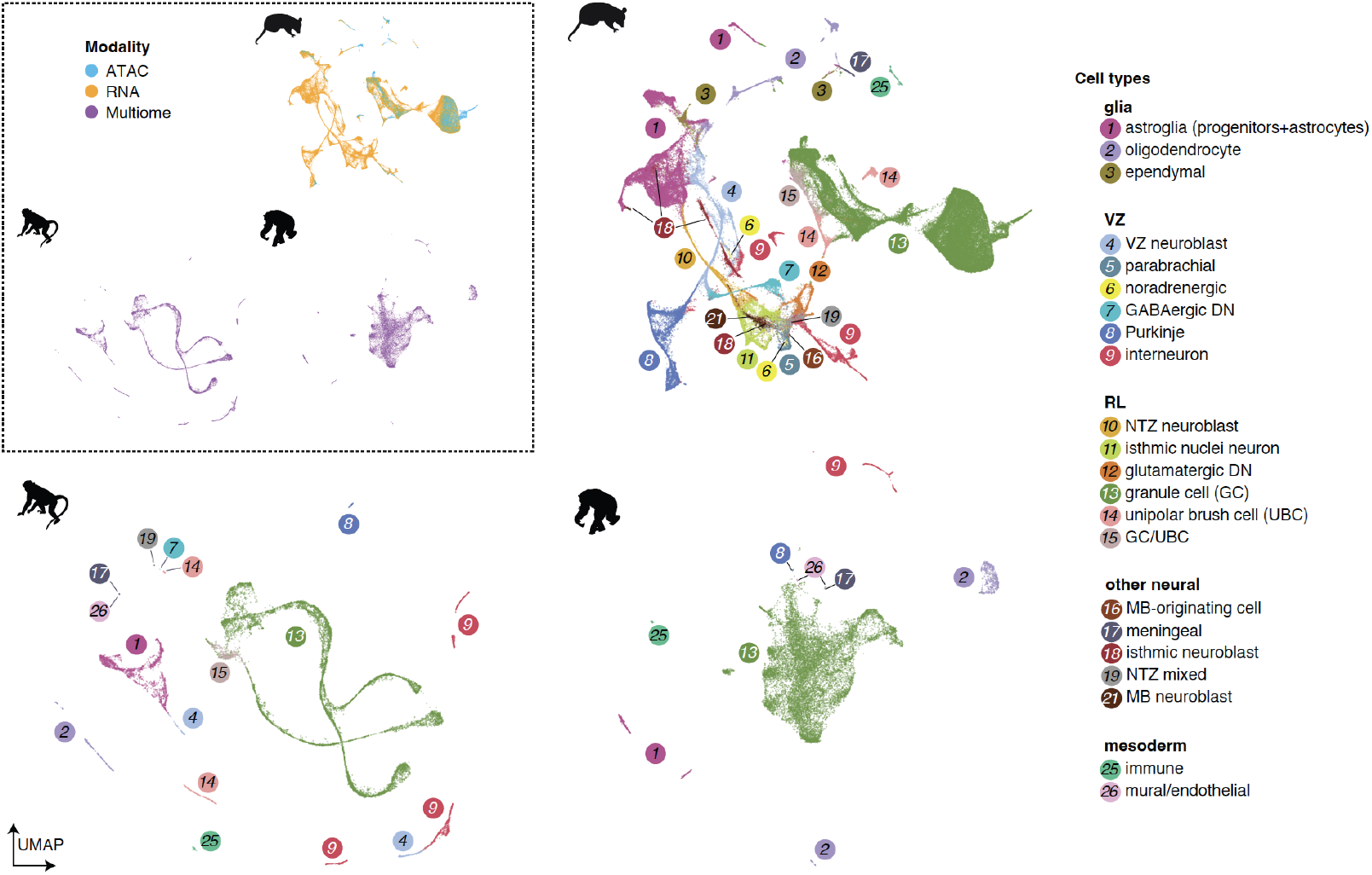
Multiomic atlases of cerebellum development in opossum, macaque and bonobo. UMAPs of opossum, macaque and bonobo single-nucleus profiles integrated across developmental stages and data modalities, coloured by modality (inset) or cell type.

**Fig. S3.**
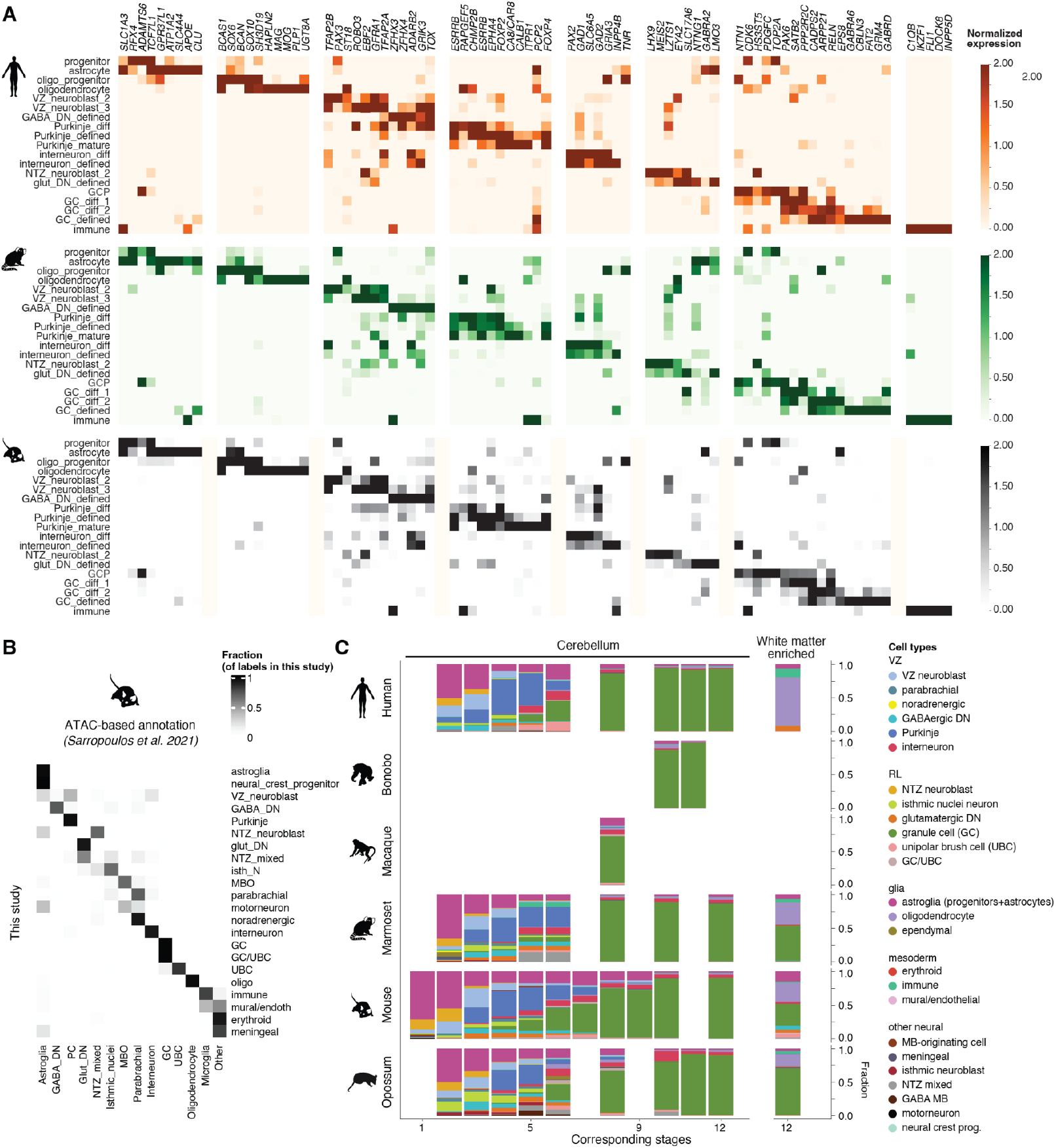
Cell types of the developing cerebellum across mammalian species. (A) Expression of marker genes across cell states in human, marmoset, and mouse. For each state top 5 therian-conserved markers (*8*) - prioritised based on enrichments in human, mouse, and opossum - are shown. Genes marking multiple states are displayed once. (B) Overlap between annotations in this study (rows) and original annotations from (*38*) (columns) for mouse cells profiled with snATAC-seq. (C) Broad cell type (level 1) fractions across species and developmental stages.

**Fig. S4.**
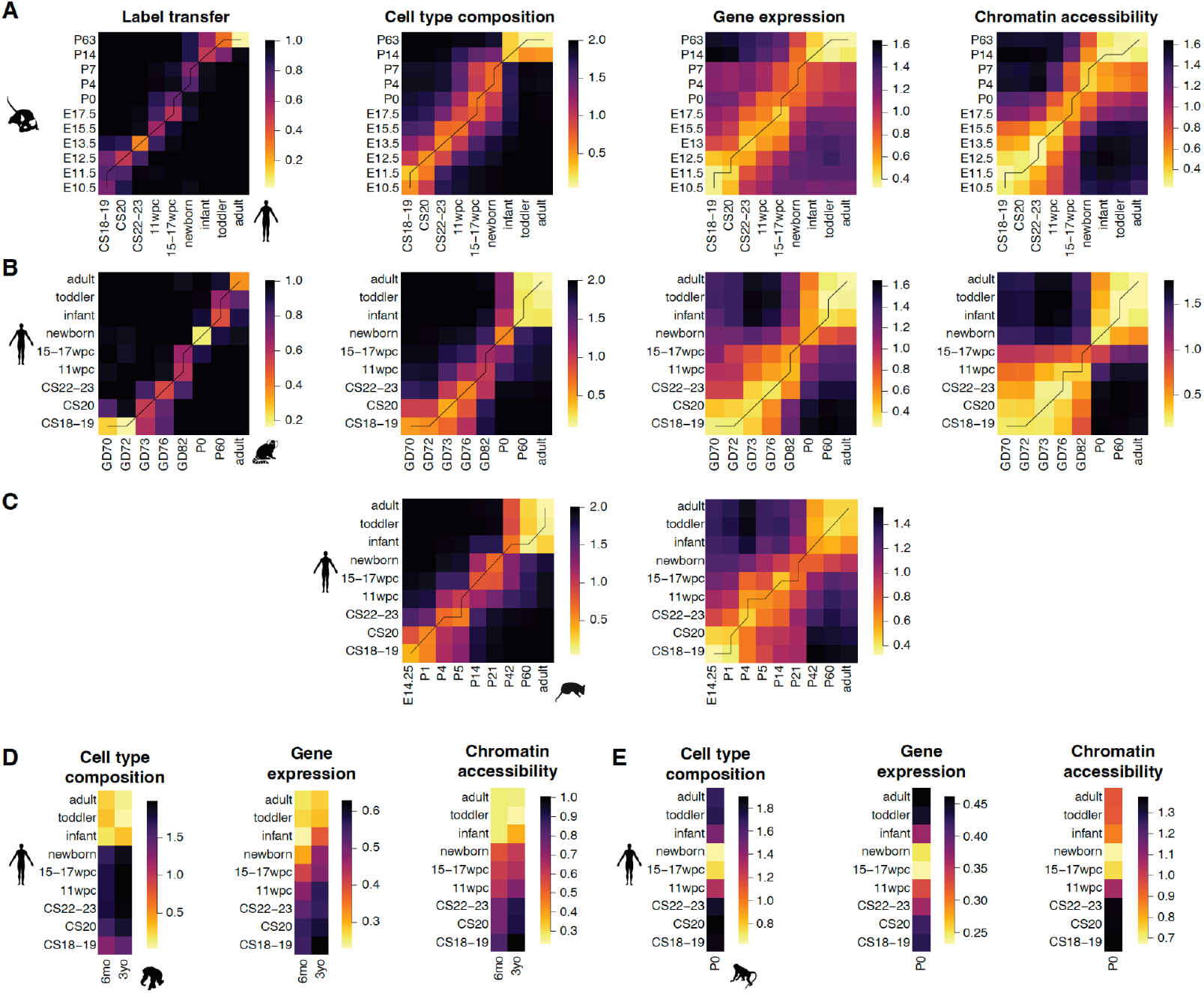
Corresponding stages of cerebellum development across mammalian species. (A-E) Dynamic time-warping alignment between human and mouse (A), marmoset (B), opossum (C), bonobo (D) or macaque (E) developmental stages based on different dissimilarity metrics. dynamic time-warping applied on different dissimilarity metrics. CS, Carnegie stage; E, embryonic/prenatal day; GD, gestational day (counted from ovulation/conception); mo, months old; P, postnatal day; wpc, weeks post conception; yo, years old.

**Fig. S5.**
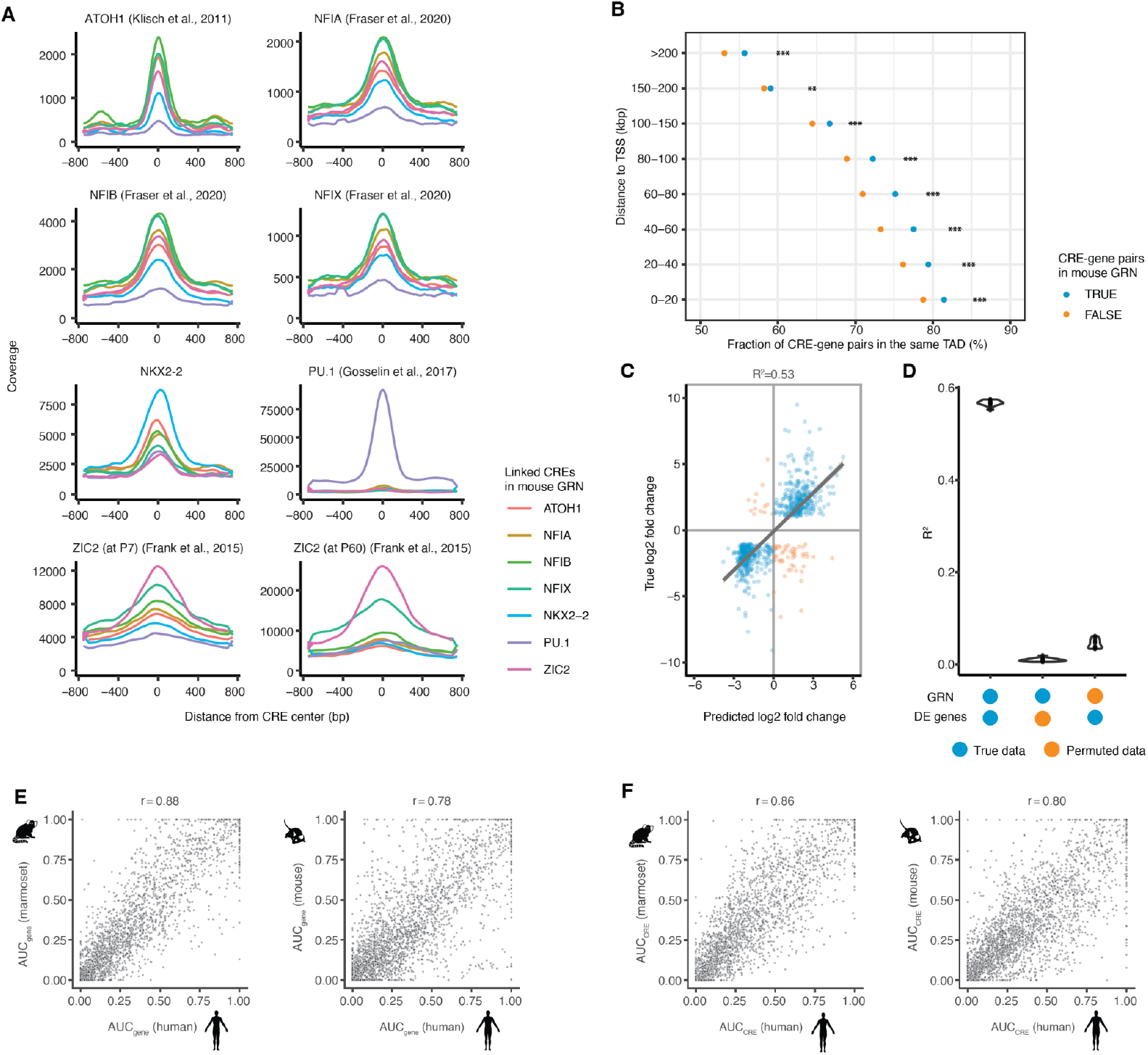
Benchmark of gene regulatory networks (GRNs) of mouse cerebellar cell types. (A) ChIP-seq coverages in CREs predicted to be bound by TFs measured by the experiments. (B) Fraction of CRE-gene pairs in the mouse GRN (blue) or not in the GRN (red), located within the same topologically associating domain (TAD) in mouse neural progenitors (*99*). *** *P* < 0.001, ** *P* < 0.01, Fisher’s exact test. (C) True versus predicted log2 fold-changes for E14 and P4 mouse cerebellum, using a random forest model trained on the mouse GRN with GRaNPA (*100*). (D) Density distribution of R2 values for 10 random forest models trained on the mouse GRN to predict differential expression, compared to two permuted controls. (E, F) Target gene-based (E) and CRE-based (F) AUC scores computed for each TF regulator using AUCell (*121*) across human, marmoset, and mouse.

**Fig. S6.**
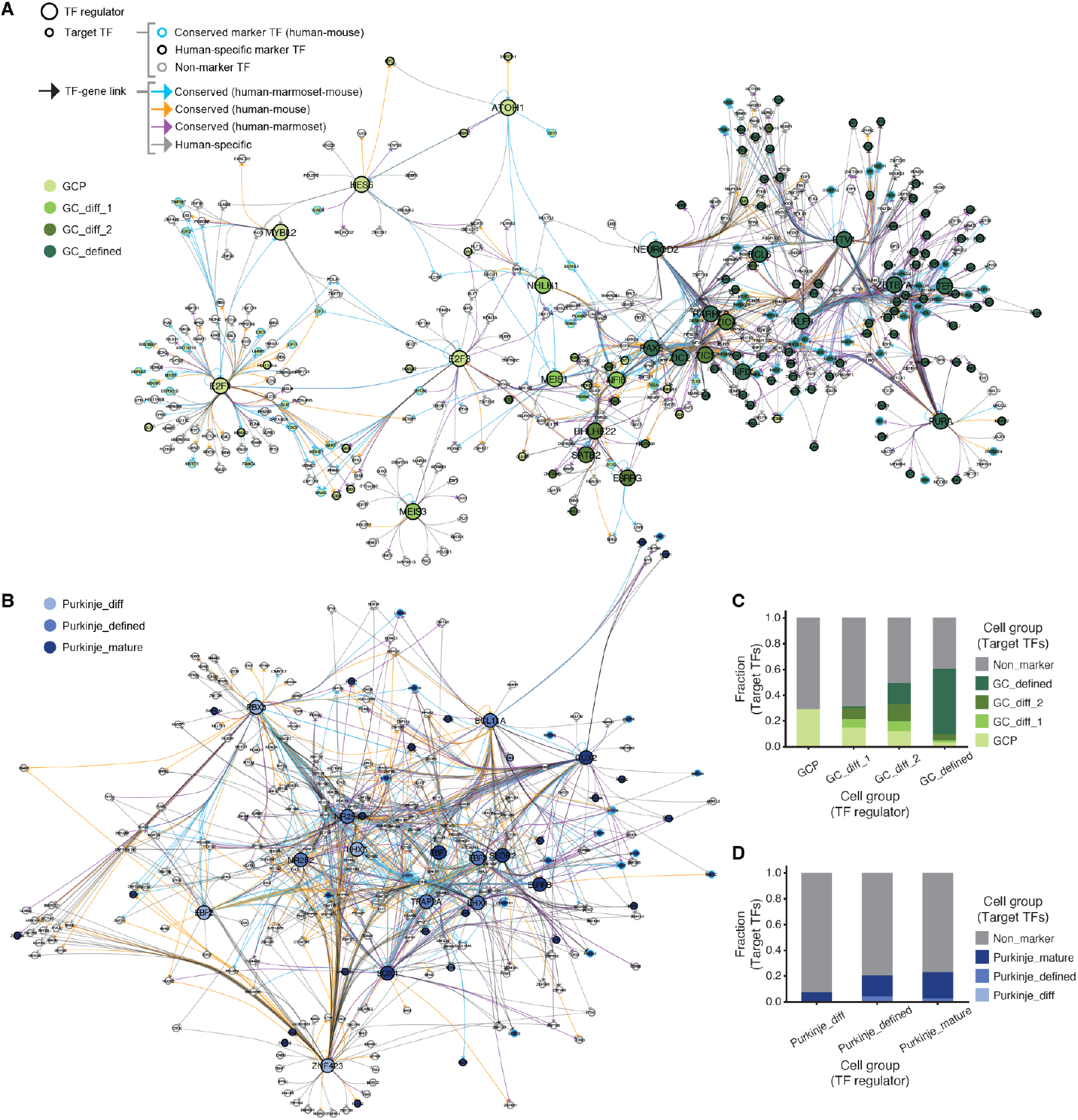
TF networks of human cerebellar cell types. (A, B) TF regulators and their target TF genes in the granule cell (A) and the Purkinje cell lineage (B) within the human gene regulatory network. (C, D) Fraction of target TF genes linked to TF regulators specific to each cell group in the granule cell (C) and the Purkinje cell lineage (D).

**Fig. S7.**
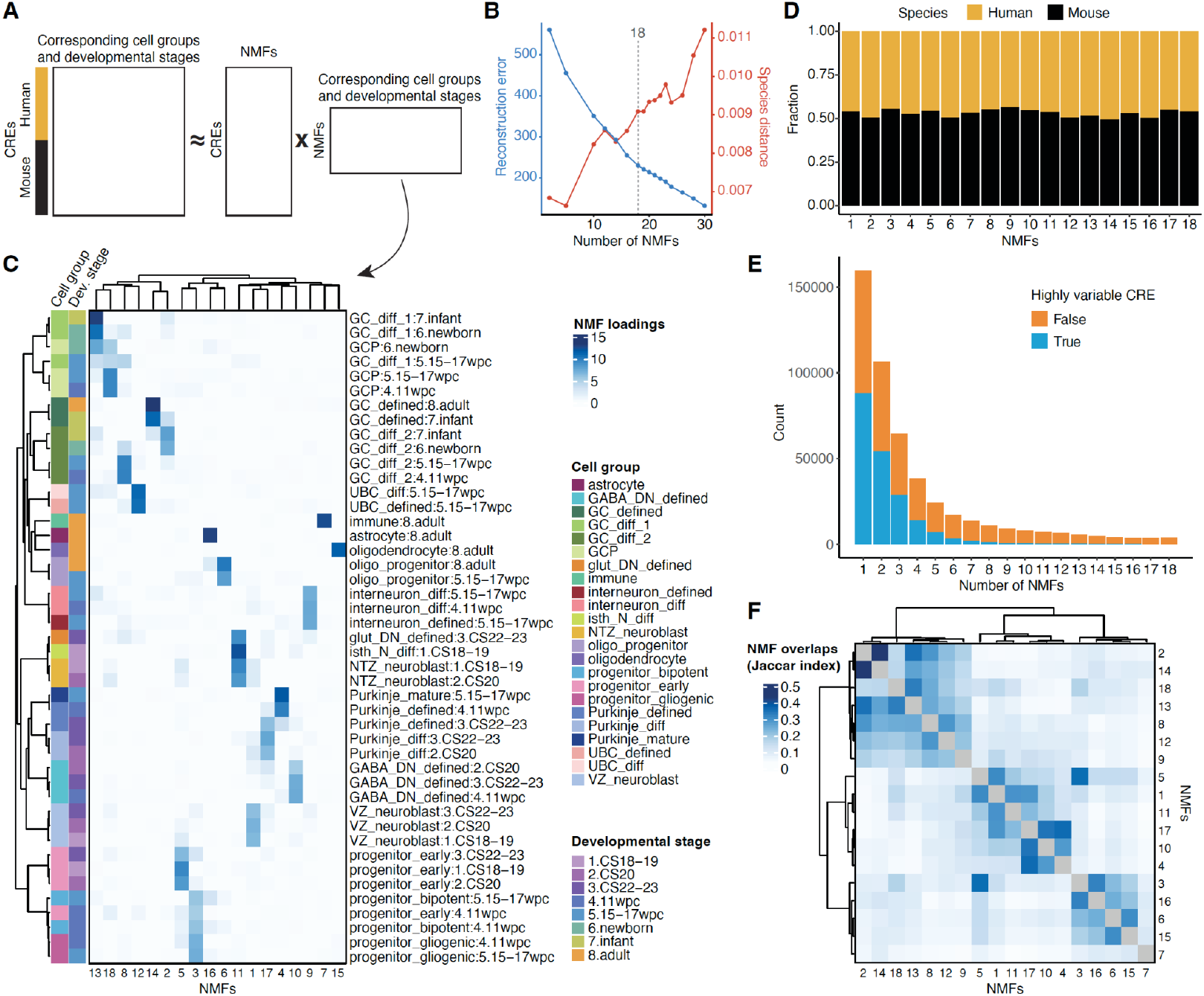
Non-negative matrix factorization of human and mouse CRE spatiotemporal accessibility. (A) Schematic representation of the non-negative matrix factorization. (B) Reconstruction error and distance between species in NMF space across a range of factors. (C) NMF loadings across corresponding cell groups and developmental stages. Rows and columns are grouped based on hierarchical clustering. (D) Fraction of human and mouse CREs assigned to each NMF. (E) Number of NMFs associated with each human and mouse CRE. (F) Overlap of assigned CREs between NMFs.

**Fig. S8.**
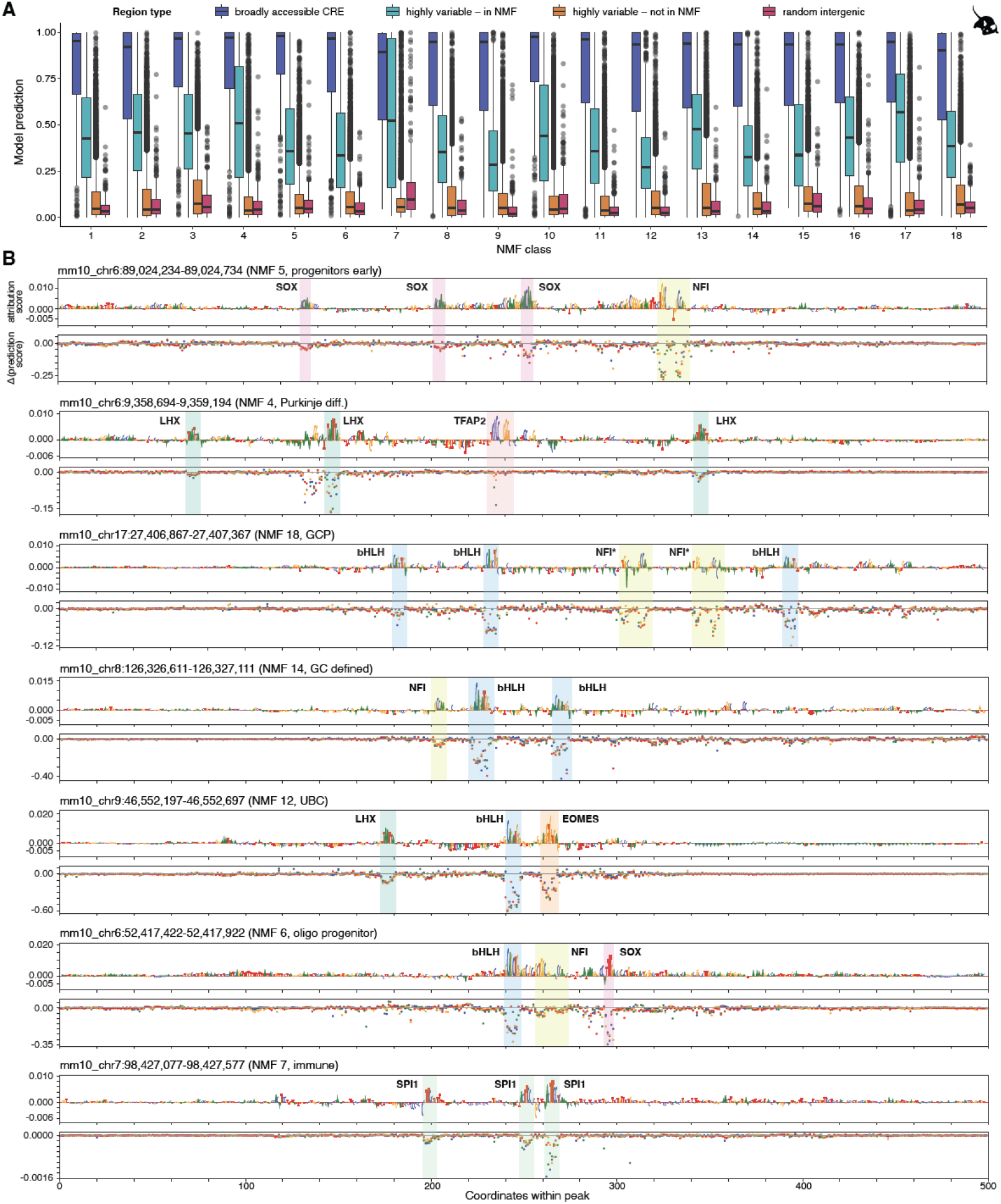
Sequence-based models of CRE accessibility in mouse cerebellar cell types. (A) Predictions in test set mouse regions across NMF classes. Boxes represent the interquartile range and whiskers extend to extreme values within 1.5 times the interquartile range from the box. (B) DeepExplainer (top) and *in silico* mutagenesis (bottom) profiles of mouse CREs in the test set with high predictions in specific NMFs. Relevant TF motifs are highlighted. *, imperfect motif.

**Fig. S9.**
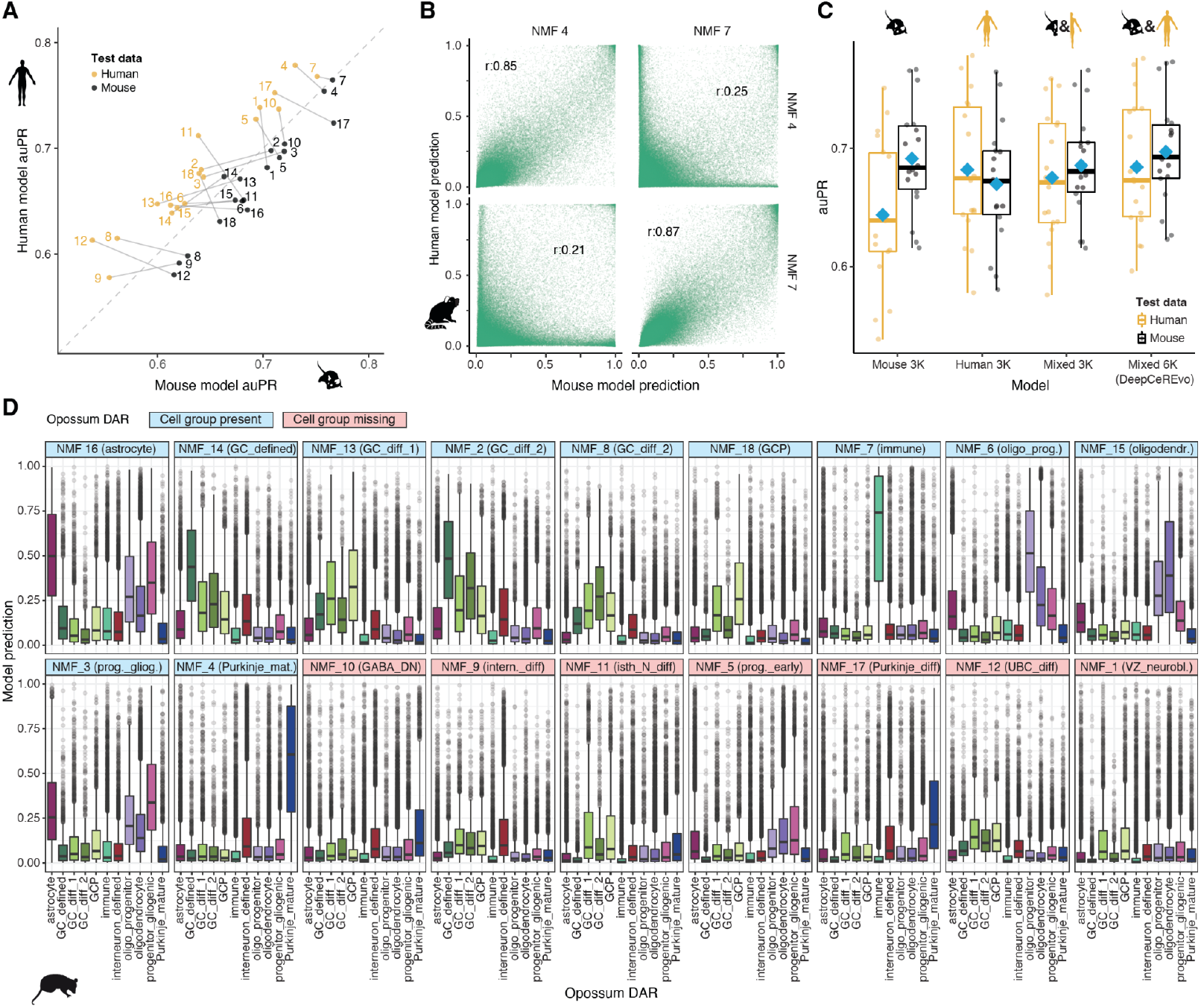
Across species predictions of sequence-based models. (A) Accuracy (auPR) of sequence-based models trained on human or mouse sequences in predicting the accessibility of unseen sequences from both species. (B) Human and mouse model predictions for marmoset sequences across the same or different NMF pairs. (C) DeepCeREvo achieves improved performance by training on both human and mouse sequences. Boxplots show auPR metrics per NMF class in human and mouse unseen sequences for models trained on CREs from human, mouse or both species. (D) DeepCeREvo’s predictions for opossum DARs. NMF annotations are based on the group with the highest loading. The presence/absence of relevant groups in the opossum dataset is shown with blue/red respectively. In C and D, boxes represent the interquartile range and whiskers extend to extreme values within 1.5 times the interquartile range from the box.

**Fig. S10.**
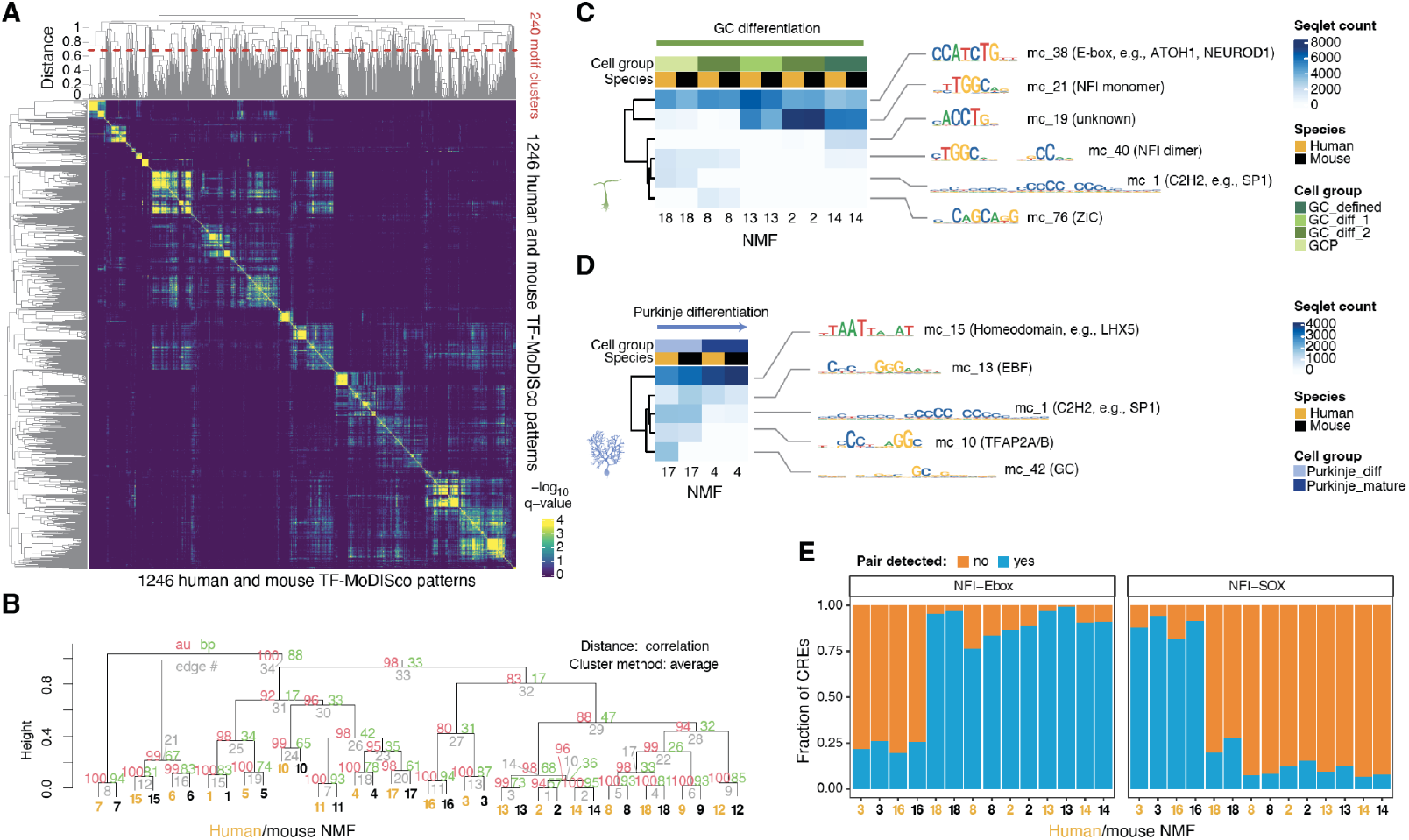
CRE sequence grammar across species and cell types. (A) Hierarchical clustering based on correlations in sequence similarity scores (−log_10_ q-value from Tomtom) across human and mouse TF-MoDISco patterns identified separately for each species and NMF. The red line indicates the distance (0.7) used to identify motif clusters. Patterns belonging to the same motif cluster were collapsed for subsequent analyses. (B) Hierarchical clustering of human and mouse NMFs based on the number of motif instances with high attribution scores (seqlets) for each NMF and motif cluster (mc). Dendrogram as in Fig. 4E with approximately unbiased (au) probability values estimated after bootstrapping with pvclust (v2.2-0) shown in red. (C, D) Seqlet counts for motif clusters with at least 1000 seqlets for human and mouse CREs assigned to NMFs associated with granule cell (C) and Purkinje cell (D) differentiation, along with their respective consensus logos. (E) Fraction of human and mouse CREs with high predictions for NMFs associated with astroglia and granule cells with one or more NFI (mc_21, mc_40) and E-box (mc_38) or SOX (mc_12) seqlets.

**Fig. S11.**
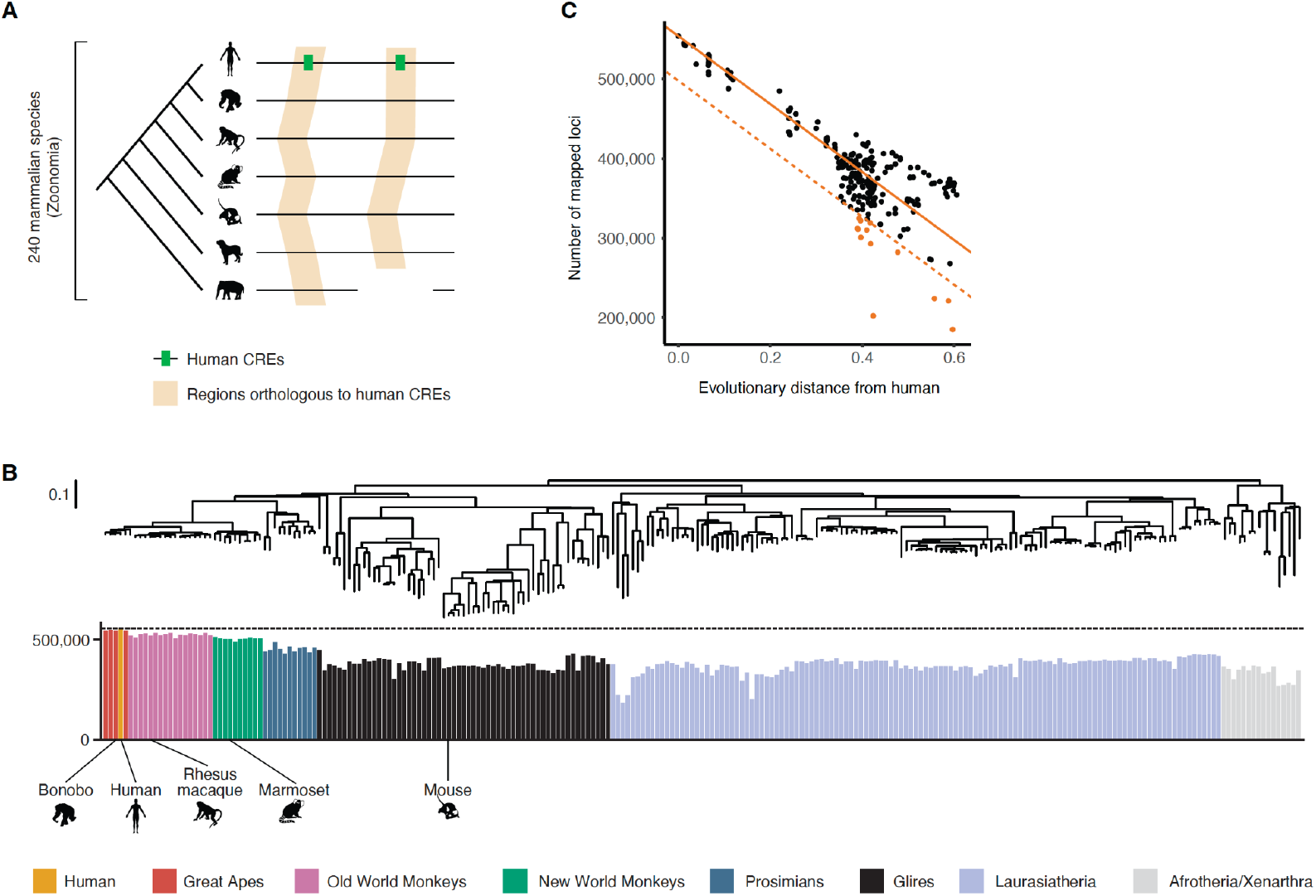
Identification of orthologous sequences of human CREs in 240 eutherian mammals. (A) Schematic illustration of regions orthologous to human CREs across eutherian mammals. (B) Phylogenetic tree of 240 mammalian species, with branch lengths scaled by substitution rate per site (top), and the number of loci in each species mapped from 554,237 human CREs (bottom). (C) Association between the number of mapped loci in each species and its evolutionary distance from humans. Thirteen species were excluded from subsequent analyses due to low mappability.

**Fig. S12.**
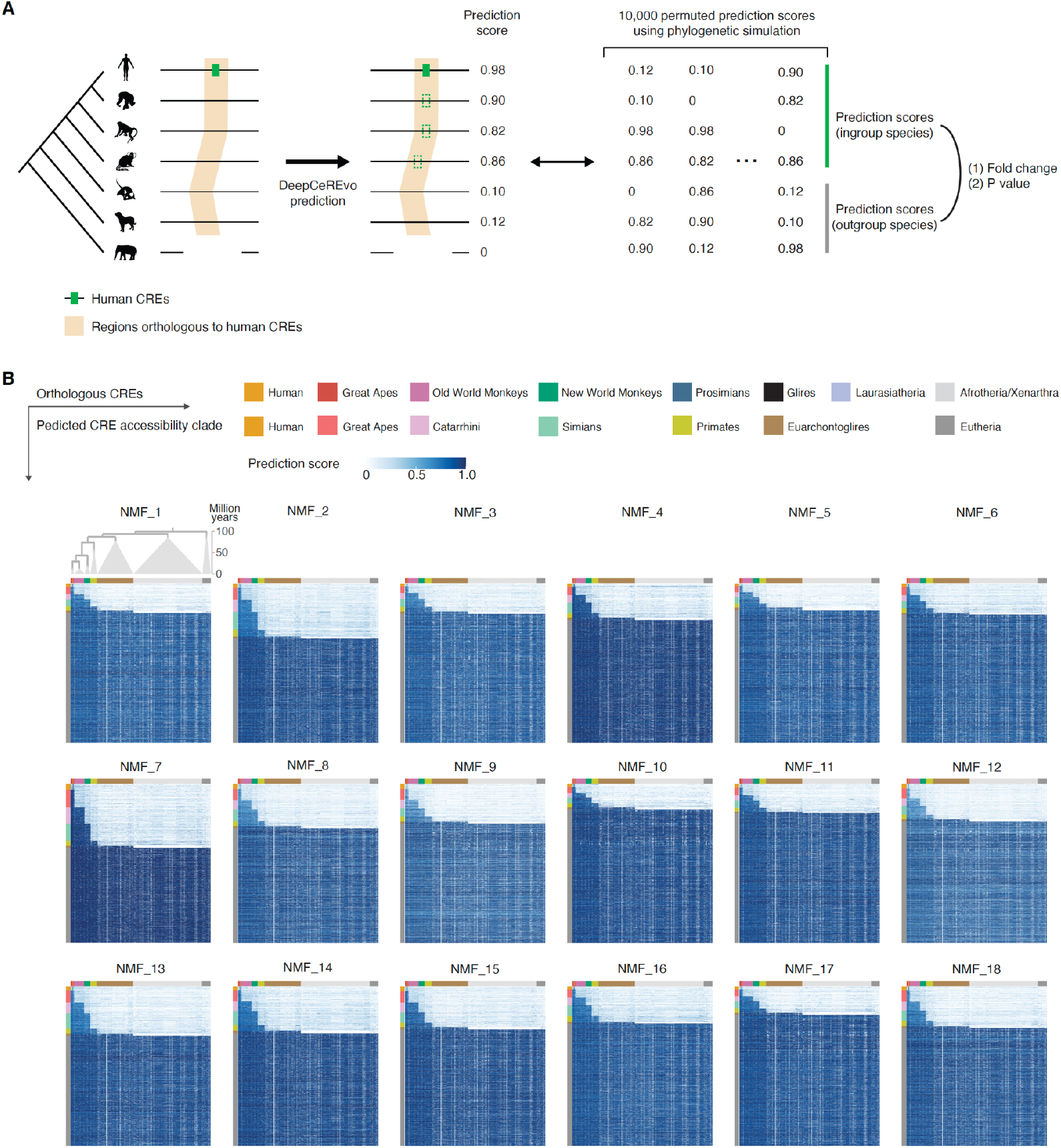
Inferring the evolutionary histories of human CREs. (A) Schematic illustration of the process used to infer the evolutionary histories of human CREs using DeepCeREvo’s predictions on orthologous regions across eutherian mammals via a phylogenetic approach. (B) Human CREs inferred to be preserved in different evolutionary clades for each NMF. Data for NMFs 5, 7, and 14 are also shown in Fig. 5A.

**Fig. S13.**
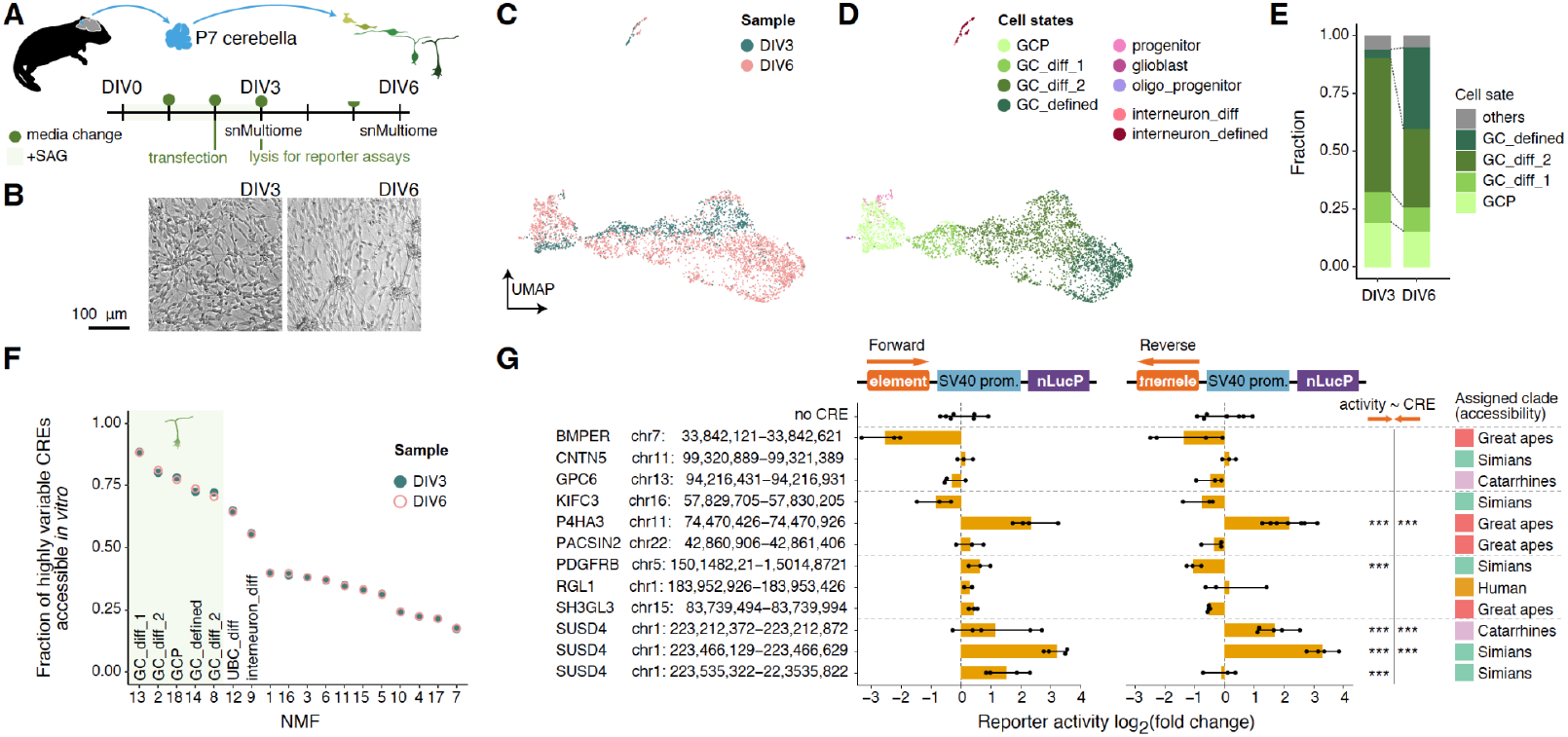
Primary cultures of mouse granule cells for testing CRE activities in reporter assays. (A) Schematic illustration of isolation and culturing of mouse P7 granule cells. DIV, days *in vitro*; SAG, Smoothened Agonist. (B) Phase contrast micrographs of granule cells at DIV3 and DIV6. (C, D) UMAPs of 1,167 DIV3 and 2,784 DIV6 granule cells, coloured by sample (C) or cell state (D). (E) Relative cell type abundances in the DIV3 and DIV6 granule cell datasets. (F) Fraction of highly variable CREs identified in the mouse (*in vivo*) snATAC-seq data that are also accessible in the mouse granule cells cultured *in vitro*. (G) Luciferase reporter assays in mouse primary granule cells testing the enhancer activity of human CREs predicted to have emerged in the last 43 million years. CREs were placed in front of the SV40 promoter in forward (left) or reverse (right) orientation. Bars and error bars display the mean normalized and scaled reporter activity and its range; points denote biological replicates. *P*-values relative to the constructs without an enhancer were estimated using linear mixed-effects models, corrected for multiple testing using the Benjamini-Hochberg method, and are shown only for bars with log_2_(fold change) ≥ 0.5. ***, *P* < 0.001.

**Fig. S14.**
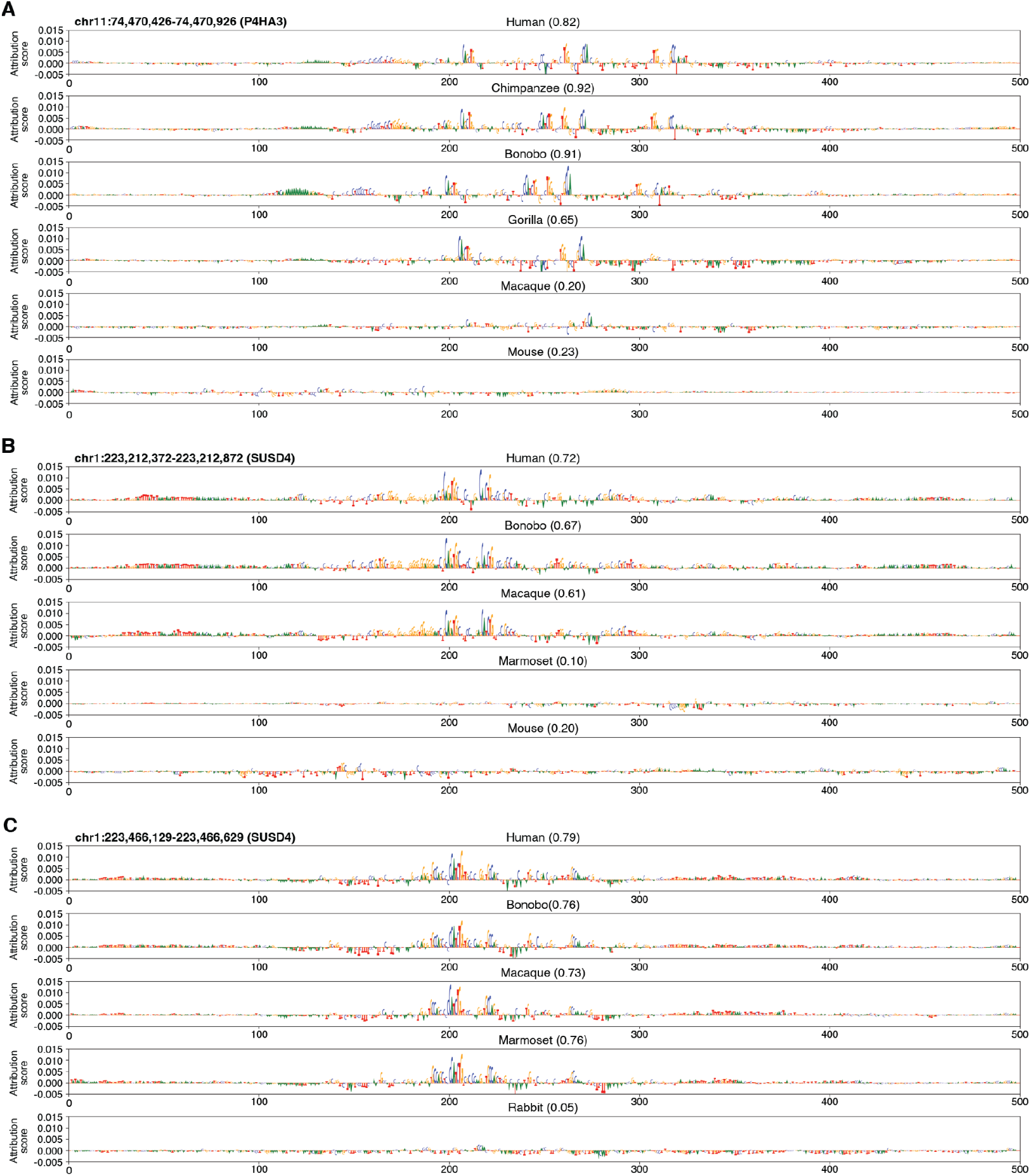
Sequence features of orthologous genomic regions tested in reporter assays. (A-C) DeepExplainer profiles of genomic regions from other mammalian species that are orthologous to the three human CREs tested in Fig. 5G. Prediction scores are shown next to each species label.

**Fig. S15.**
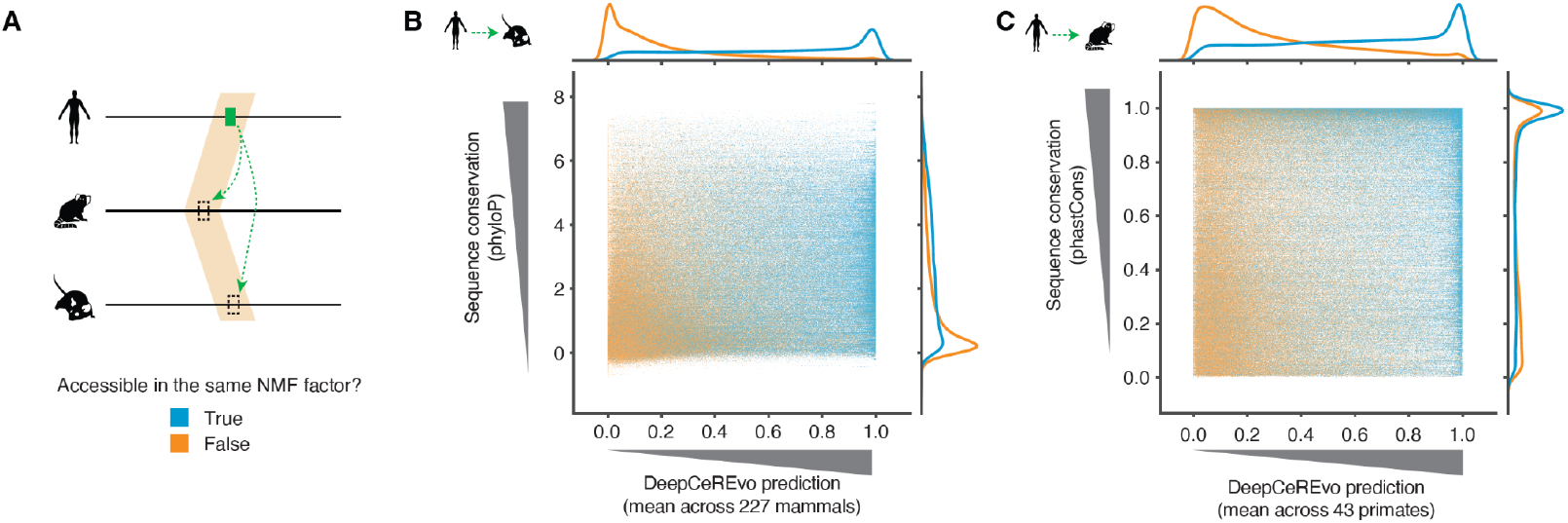
Performance of DeepCeREvo and sequence conservation in predicting chromatin accessibility in mouse and marmoset. (A) Schematic illustration of predicting chromatin accessibility in mouse and marmoset genomic loci based on orthologous human loci containing CREs. (B, C) DeepCeREvo prediction scores and sequence conservation scores for mouse (B) and marmoset (C) regions orthologous to human CREs, displayed for each combination of human CRE and NMF factor. Points are colored based on whether the orthologous regions in mouse (B) or marmoset (C) are accessible.

**Fig. S16.**
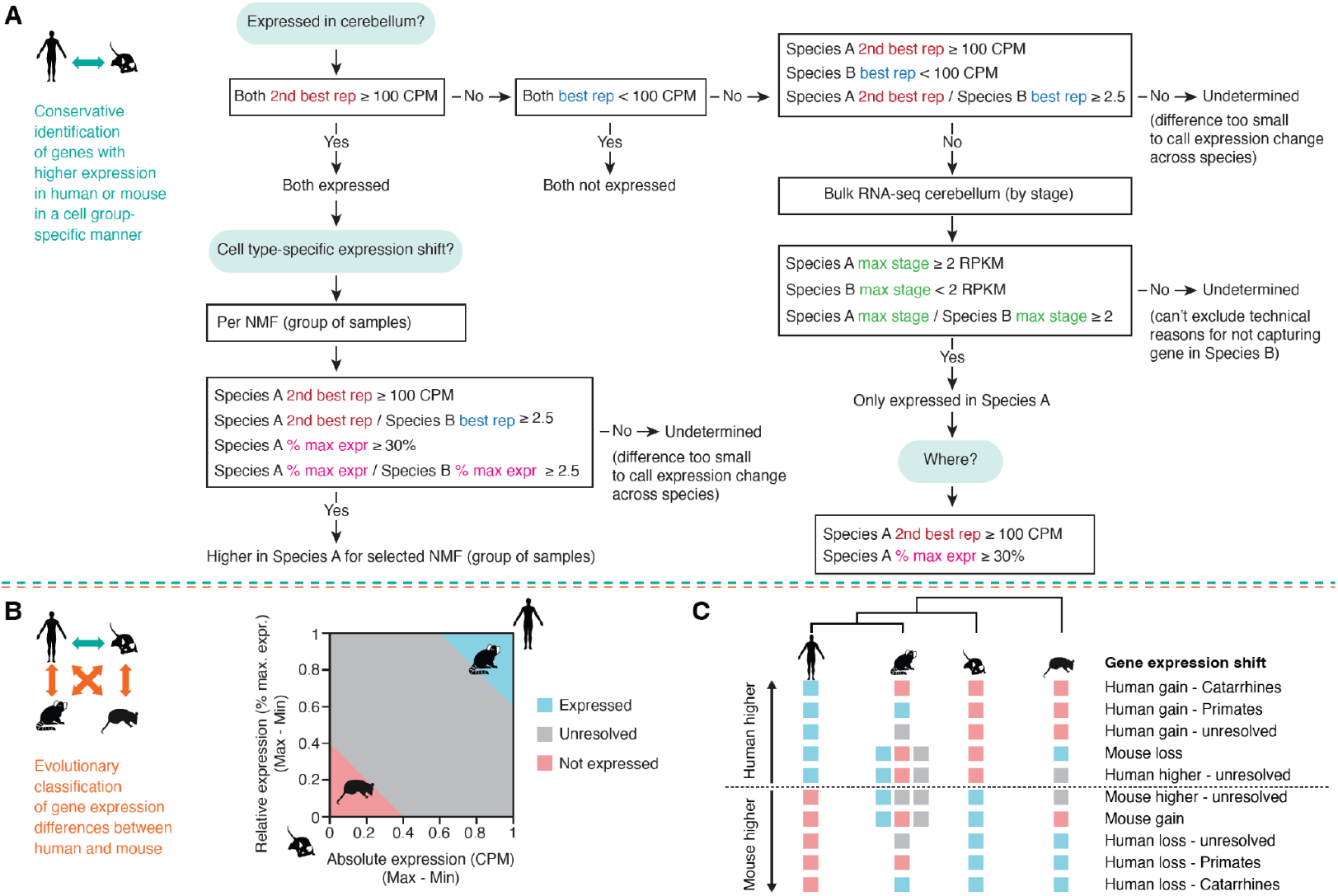
Schematic overview of evolutionary characterization of gene expression differences between human and mouse. (A) Pipeline developed for the detection of cell-group-specific radical gene expression differences between human and mouse. Genes are broadly divided into those expressed in neither, one or both species. For those expressed in one species, additional evidence from bulk RNA-seq data is used to minimize the risk that lack of detection in the other species is due to technical limitations of 3’ technologies. For genes detected in both species, comparisons are performed per NMF (cell group) considering both absolute and relative expression levels. (B) Expression classification of human-mouse differentially expressed genes in marmoset and opossum datasets. In this example, a gene is more highly expressed in human compared to mouse. Absolute (quantile-normalized CPM) and relative (fraction of maximum across cell groups) expression values in each species are scaled based on the maximum (human) and minimum (mouse) values. The mean scaled absolute and relative expression for marmoset/opossum is greater/smaller than 0.8/0.2 and the gene is classified as expressed/non-expressed, respectively. (C) Evolutionary classification of human-mouse differentially expressed genes based on the expression class assigned in marmoset and opossum.

**Fig. S17.**
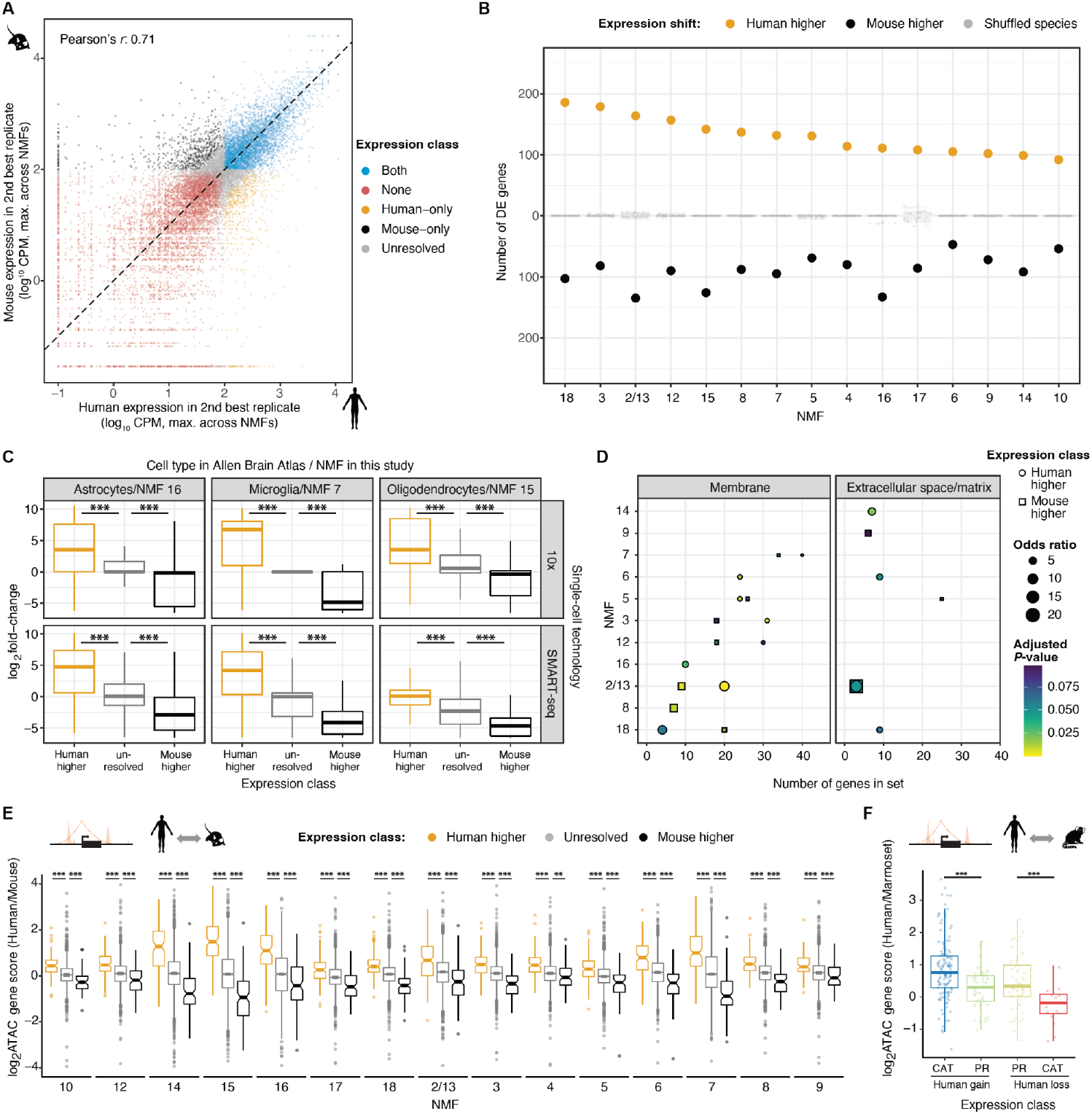
Evolutionary shifts in cell type-specific gene expression. (A) Maximum expression across NMFs of human and mouse 1:1 orthologs. Genes reaching at least 100 CPM in at least 2 replicates of at least one cell group in both species (expression class: both) were considered for the detection of cell type-specific shifts. (B) Number of differentially expressed (DE) genes detected per NMF compared to 100 random permutations in which species labels were swapped in half of the cell groups. (C) Log_2_ fold-change in the expression of genes with conserved or divergent expression in independent datasets from the Allen Brain Atlas resource across two distinct single-cell sequencing technologies. (D) Enrichment of genes with higher expression in human or mouse, compared to all genes with high expression in the same NMF and species, for localisation to the membrane (left) and extracellular space (right). (E) Comparison of local chromatin accessibility (ATAC-based gene scores) between human and mouse for genes with significantly higher expression in either species across NMFs. Data as in Fig. 6B but split across NMFs. (F) Comparison of local chromatin accessibility (ATAC-based gene scores) between human and marmoset for genes with a human-specific (CAT) or shared with marmoset (PR) gain or loss of expression. In C, E, F, boxes represent the interquartile range and whiskers extend to extreme values within 1.5 times the interquartile range from the box. *** *P* < 0.001, ** *P* < 0.01, Mann-Whitney *U* test.

**Fig. S18.**
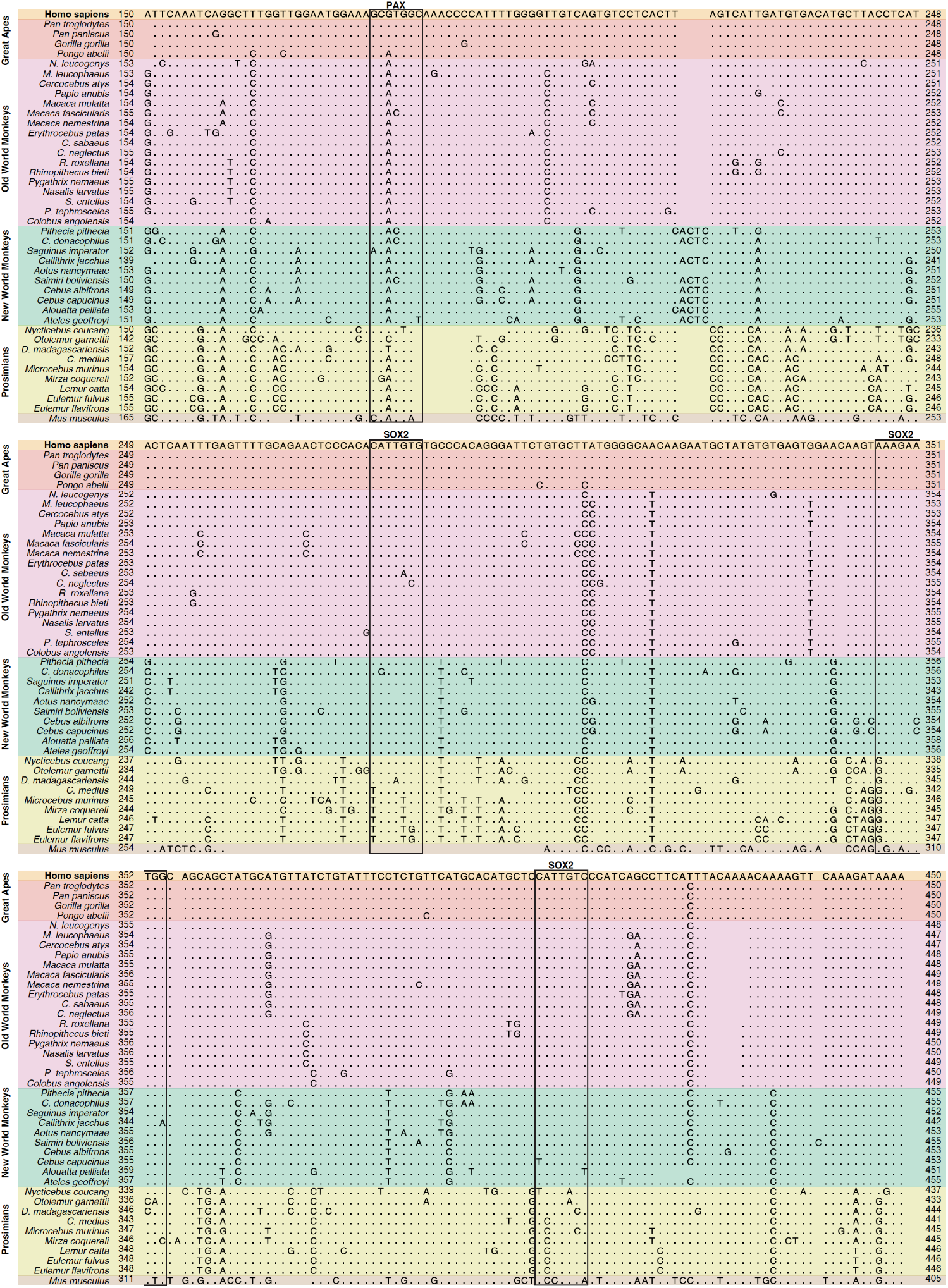
Sequence conservation of the ~3kb upstream CRE in *THRB*. Multiple sequence alignment for human chr3:24,498,438-24,498,938 across 43 primates and mouse is shown.

## LIST OF SUPPLEMENTARY TABLES

**Table S1**. Single-cell RNA- and ATAC-seq libraries

**Table S2**. Bulk RNA-seq libraries

**Table S3**. Developmental correspondences between species

**Table S4**. AUC scores of target genes for each regulon in each cell group of the human GRN.

**Table S5**. AUC scores of target genes for each regulon in each cell group of the marmoset GRN.

**Table S6**. AUC scores of target genes for each regulon in each cell group of the mouse GRN.

**Table S7**. Public ChIP-seq datasets used for benchmarking mouse gene regulatory networks.

**Table S8**. Cell group and developmental stage with the highest loading per NMF factor.

**Table S9**. Performance metrics of deep-learning models across NMF factors and species.

**Table S10**. Species included in the Zoonomia dataset.

**Table S11**. Prediction score thresholds for ingroup species based on F1 scores.

**Table S12**. Number of human CREs assigned to different evolutionary clades per NMF factor.

**Table S13**. Sequences used for cloning reporter constructs and results from reporter assays.

Log_2_(fold change) values of reporter activity relative to the no-enhancer control constructs are reported for elements in forward (F) and reverse (R) orientation. *P*-values and t values are from linear mixed model, including both main CRE effects (forward and reverse) and interaction effects (forward versus reverse). *P*-values were corrected for multiple comparisons using the Benjamini-Hochberg method.

**Table S14**. Evolutionary classification of differentially expressed genes between human and mouse.

**Table S15**. Gene ontology enrichments for genes differentially expressed between human and mouse.

**Table S16**. Human CREs linked to genes with higher expression in human compared to mouse.

